# The GTEx Consortium atlas of genetic regulatory effects across human tissues

**DOI:** 10.1101/787903

**Authors:** François Aguet, Alvaro N Barbeira, Rodrigo Bonazzola, Andrew Brown, Stephane E Castel, Brian Jo, Silva Kasela, Sarah Kim-Hellmuth, Yanyu Liang, Meritxell Oliva, Princy E Parsana, Elise Flynn, Laure Fresard, Eric R Gaamzon, Andrew R Hamel, Yuan He, Farhad Hormozdiari, Pejman Mohammadi, Manuel Muñoz-Aguirre, YoSon Park, Ashis Saha, Ayellet V Segrć, Benjamin J Strober, Xiaoquan Wen, Valentin Wucher, Sayantan Das, Diego Garrido-Martín, Nicole R Gay, Robert E Handsaker, Paul J Hoffman, Seva Kashin, Alan Kwong, Xiao Li, Daniel MacArthur, John M Rouhana, Matthew Stephens, Ellen Todres, Ana Viñuela, Gao Wang, Yuxin Zou, The GTEx Consortium, Christopher D Brown, Nancy Cox, Emmanouil Dermitzakis, Barbara E Engelhardt, Gad Getz, Roderic Guigo, Stephen B Montgomery, Barbara E Stranger, Hae Kyung Im, Alexis Battle, Kristin G Ardlie, Tuuli Lappalainen

## Abstract

The Genotype-Tissue Expression (GTEx) project was established to characterize genetic effects on the transcriptome across human tissues, and to link these regulatory mechanisms to trait and disease associations. Here, we present analyses of the v8 data, based on 17,382 RNA-sequencing samples from 54 tissues of 948 post-mortem donors. We comprehensively characterize genetic associations for gene expression and splicing in *cis* and *trans*, showing that regulatory associations are found for almost all genes, and describe the underlying molecular mechanisms and their contribution to allelic heterogeneity and pleiotropy of complex traits. Leveraging the large diversity of tissues, we provide insights into the tissue-specificity of genetic effects, and show that cell type composition is a key factor in understanding gene regulatory mechanisms in human tissues.

## Introduction

A pressing need in human genetics remains the characterization and interpretation of the function of the millions of genetic variants across the human genome. This is essential for identifying the molecular mechanisms of genetic risk for complex traits and diseases, mainly driven by non-coding loci with largely unknown regulatory functions. To address this challenge, several projects have built comprehensive annotations of genome function across tissues and cell types (*1, 2*), and mapped the effects of regulatory variation across large numbers of individuals, primarily from whole blood and blood cell types (*3-5*). The Genotype-Tissue Expression (GTEx) project provides an essential intersection where variant function can be studied across a wide range of both tissues and individuals.

The GTEx project was launched in 2010 with the aim of building a catalog of genetic effects on gene expression across a large number of human tissues in order to elucidate the molecular mechanisms of genetic associations with complex diseases and traits, and improve our understanding of regulatory genetic variation (*6*). The project set out to collect biospecimens from ∼50 tissues from up to ∼1000 postmortem donors, and to create standards and protocols for optimizing postmortem tissue collection and donor recruitment (*7, 8*), biospecimen processing (*7*), and data sharing (www.gtexportal.com).

Following the earlier publication of the GTEx pilot (*9*) and mid-stage results (*10*), we present a final analysis from the GTEx Consortium based on the v8 data release. We provide the largest catalog to date of genetic regulatory variants affecting gene expression and splicing in *cis* and *trans* across 49 tissues, and describe patterns and mechanisms of tissue- and cell type specificity of genetic regulatory effects. Through integration of GTEx data with genome-wide association studies (GWAS), we characterize mechanisms of how genetic effects on the transcriptome mediate complex trait associations.

### QTL discovery

The GTEx v8 data set consists of 948 donors and 17,382 samples from 52 tissues and two cell lines, with 838 donors and 15,253 samples having both RNA sequence (RNA-seq) and genotype data from whole genome sequencing (WGS) (figs. 1a, S1–2). The 838 donors were 85.3% European American, 12.3% African American, and 1.4% Asian American. Of the 54 tissues, 49 had samples from at least 70 individuals and were used for analyses of quantitative trait loci (QTL) (15,201 samples total). WGS was performed for each donor to a median depth of 32x, resulting in the detection of a total of 43,066,422 single nucleotide polymorphisms (SNPs) after QC and phasing (10,008,325 with MAF ≥ 0.01) and 3,459,870 small indels (762,535 with MAF ≥ 0.01) (fig. S3, table S1, (*11*)). The mRNA of each of the tissue samples was sequenced to a median depth of 82.6 million reads, and alignment, quantification and quality control were performed as described in (*11*) (figs. S4–5).

**Figure 1.**
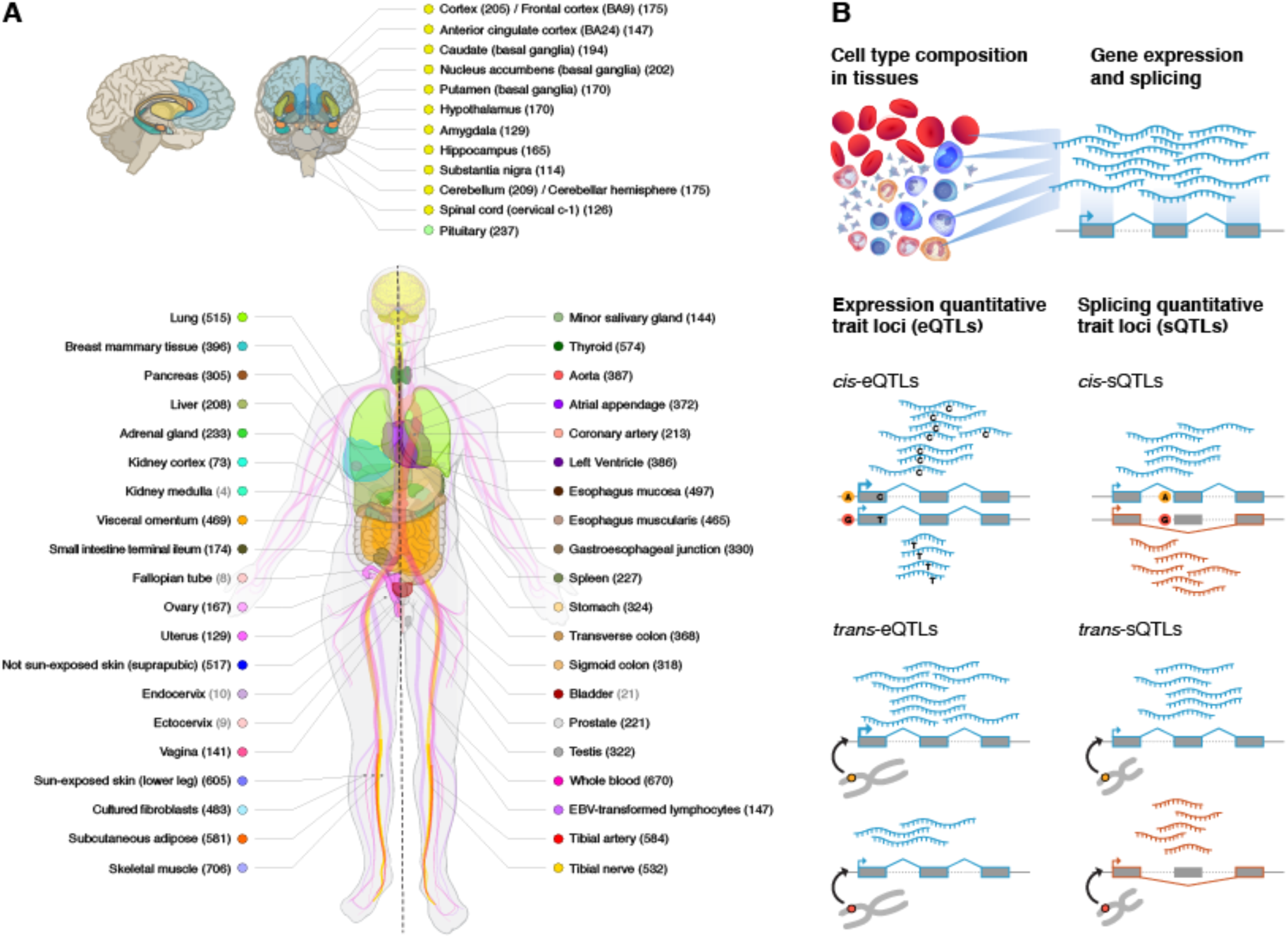
Sample and data types in the GTEx v8 study. (A) Illustration of the 54 tissue types (including 11 distinct brain regions and 2 cell lines), with sample numbers from genotyped donors in parentheses and color coding indicated in the adjacent circles. Tissues with ≥70 samples were included in QTL analyses. (B) Illustration of the core data types used throughout the study. Gene expression and splicing were quantified from bulk RNA-seq of heterogenous tissue samples, and local and distal genetic effects (cis-QTLs and trans-QTLs, respectively) were quantified across individuals for each tissue.

The resulting data provide the broadest survey of individual- and tissue-specific gene expression to date, enabling a comprehensive view of the impact of genetic variation on gene expression and splicing (fig. 1b). Across all tissues, we discovered *cis*-eQTLs (5% FDR, per tissue (*11*)) for 18,262 protein coding and 5,006 lincRNA genes (23,268 total *cis*-eGenes, corresponding to 94.7% of all protein coding and 67.3% of all detected lincRNA genes detected in at least one tissue), with a total of 4,278,636 genetic variants (43% of all variants with MAF ≥ 0.01) that were significant in at least one tissue (*cis*-eVariants) (figs. 2a, S6, table S2). *Cis*-eQTLs for all long non-coding RNAs (lncRNAs) are characterized in a companion analysis (*12*). The genes lacking a *cis*-eQTL were enriched for those lacking expression in the tissues analyzed by GTEx, including genes involved in early development (fig. S7). While most of the discovered *cis*-eQTLs had small effect sizes measured as allelic fold change (aFC), across tissues an average of 22% of *cis*-eQTLs had an over 2-fold effect on gene expression (fig. S10). We mapped splicing QTLs in *cis* with intron excision ratios from LeafCutter (*11, 13*), and discovered 12,828 (66.5%) protein coding and 1,600 (21.5%) lincRNA genes (14,424 total) with a *cis*-sQTL (5% FDR, per tissue) in at least one tissue (*cis*-sVariants) (fig. 2a, table S2). As expected (*10*), *cis*-QTL discovery was highly correlated with the sample size for each tissue (Spearman’s rho = 0.95 for *cis*-eQTLs, 0.92 for *cis*-sQTLs).

**Figure 2.**
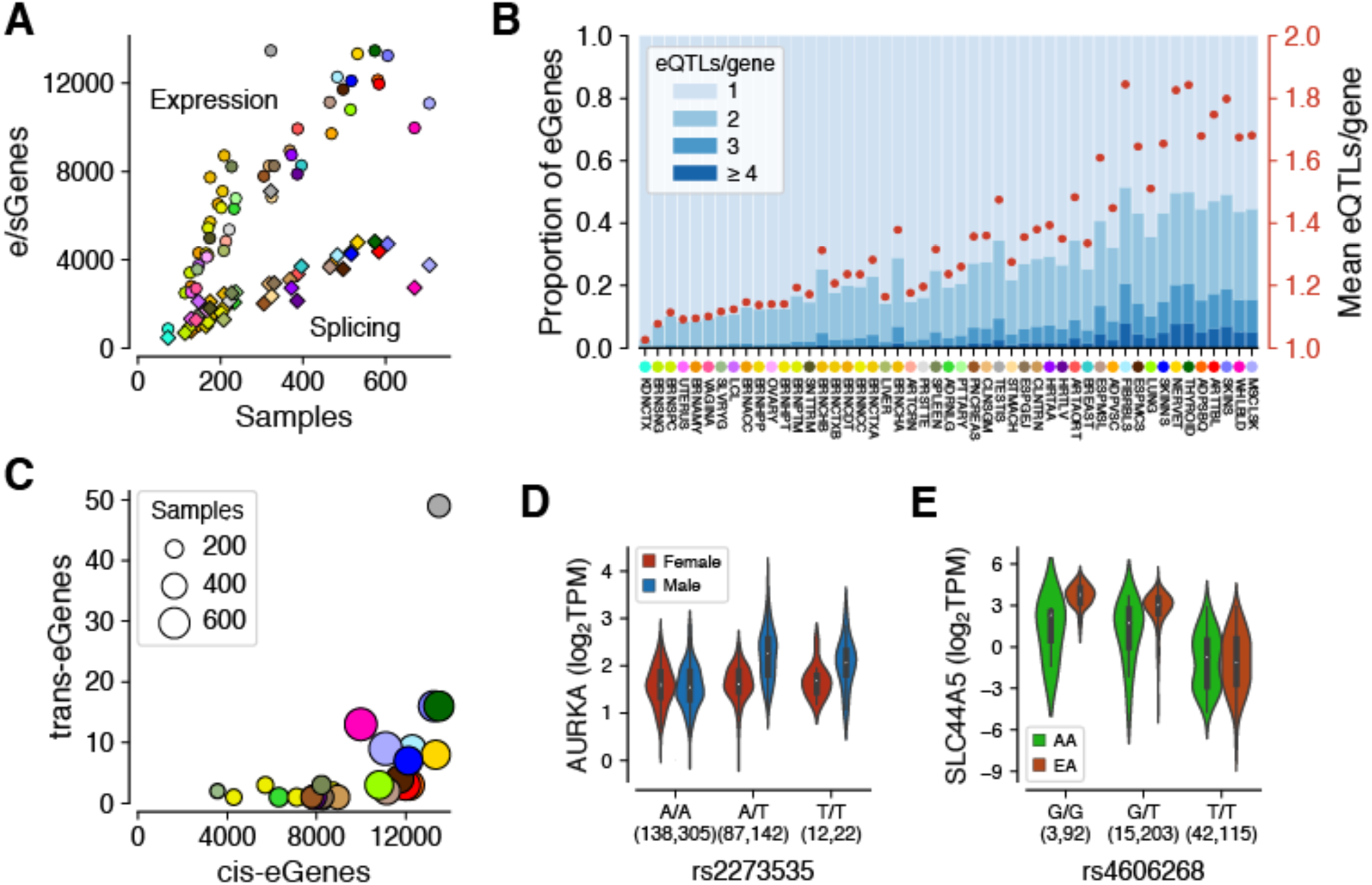
QTL discovery. (**A**) The number of genes with a *cis*-eQTL (eGenes) or *cis*-sQTL (sGenes) per tissue, as a function of sample size. See Fig. 1A for the legend of tissue colors. (**B)** Allelic heterogeneity of *cis*-eQTLs depicted as proportion of eGenes with ≥1 independent *cis*-eQTLs (blue stacked bars; left y-axis) and as a mean number of *cis*-eQTLs per gene (red dots; right y-axis). The tissues are ordered by sample size. **(C)** The number of genes with a *trans*-eQTL as a function of the number of *cis*-eGenes. **(D)** Sex-biased *cis*-eQTL for *AURKA* in skeletal muscle, where rs2273535-T is associated with increased *AURKA* expression in males (p = 9.02×10^−27^) but not in females (p = 0.75). (**E**) Population-biased *cis*-eQTL for *SLC44A5* in esophagus mucosa (allelic fold change = −2.85 and −4.82 in African Americans (AA) and European Americans (EA), respectively; permutation p-value = 1.2×10^−3^).

Previous studies have shown widespread allelic heterogeneity of gene expression in *cis*, i.e., multiple independent causal eQTLs per gene (*4, 14, 15*). We used two approaches to characterize this: 1) stepwise regression to identify conditionally independent *cis*-eQTLs, where the threshold for significance was defined by the single *cis*-eQTL mapping (*10*), and 2) a Bayesian approach where the posterior probability of linked variants was used to control the local FDR (*11, 16*). Both methods showed concordant results of widespread allelic heterogeneity, with up to 50% of eGenes having more than one independent *cis*-eQTL in the tissues with the largest sample sizes (figs. 2b, S8). Our analysis captured a lower rate of allelic heterogeneity for *cis*-sQTLs, which can be a result of both underlying biology and lower power in *cis*-sQTL mapping (fig. S8). These results highlight continued gains in *cis*-eQTL mapping with increasing sample sizes even when the discovery of new eGenes in specific tissues starts to saturate.

*Trans*-eQTL mapping yielded 143 *trans*-eGenes (121 protein coding and 22 lincRNA at 5% FDR assessed at the gene level, separately for each gene type), after controlling for false positives due to read misalignment (*11, 17*) (table S13). The number of *trans*-eGenes discovered per tissue is correlated with sample size (Spearman’s rho = 0.68), and to the number of *cis*-eQTLs (rho = 0.77), with outlier tissues such as testis contributing disproportionately to both *cis* and *trans* (fig. 2c). We identified a total of 49 *trans*-eGenes in testis, with 47 found in no other tissue even at FDR 50%. Over two-fold effect sizes on *trans*-eGene expression were observed for 19% of *trans*-eQTLs (fig. S10). *Trans*-sQTLs mapping yielded 29 *trans*-sGenes (5% FDR, per tissue), including a replication of a previously described trans-sQTL (*3*) and visual support of the association pattern in several loci (*11*) (fig. S9, table S14). These results suggest that while trans-sQTL mapping is challenging, we can discover robust genetic effects on splicing in *trans*.

We produced allelic expression (AE) data using two complementary approaches (*11*). In addition to the conventional AE data for each heterozygous genotype, we produced AE data by haplotypes, integrating data from multiple heterozygous sites in the same gene, yielding 153 million gene-level measurements (≥8 reads) across all samples (*18*). Allelic expression reflects differential regulation of the two haplotypes in individuals that are heterozygous for a regulatory variant in *cis*; indeed, *cis*-eQTL effect size is strongly correlated with allelic expression (median rho = 0.82) (*10*). We hypothesized that *cis*-sQTLs could also partially contribute to allelic imbalance even if only for parts of transcripts. However, there is drastically less signal of increased allelic imbalance among individuals heterozygous for *cis*-sQTLs (median Spearman’s rho = −0.05) (fig. S11). This indicates that allelic expression data captures primarily *cis*-eQTL effects and genetic splicing variation in *cis* is not strongly reflected in gene-level AE data.

### Genetic regulatory effects across populations and sexes

Variability in human traits and diseases between sexes and population groups is likely to partially derive from differences in genetic effects (*19-21*). To study this, we analyzed variable *cis*-eQTL effects between males and females, as well as between individuals of European and African ancestry. Since external replication data sets are sparse, we use a novel allelic expression approach for validation with an orthogonal data type from the same samples (*18*): allelic imbalance in individuals heterozygous for the *cis*-eQTL allows individual-level quantification of the *cis*-eQTL effect size (*22*), and can be correlated with the interaction terms used in *cis*-eQTL analysis to validate modifier effects of the *cis*-eQTL association.

To characterize sex-differentiated genetic effects on gene expression in GTEx tissues, we mapped sex-biased *cis*-eQTLs (sb-eQTLs). Analyzing the set of all conditionally independent *cis*-eQTLs, we identified eQTLs with significantly different effects between sexes by fitting a linear regression model and testing for a significant genotype-by-sex (G×S) interaction (*11*). Across the 44 GTEx tissues shared among sexes, we identified 369 sb-eQTLs (FDR ≤ 25%), characterized further in (*23*). Sex-biased eQTL discovery had a modest correlation with tissue sample size (Spearman’s rho = 0.39, p = 0.03), with most sb-eQTLs discovered in breast but also in muscle, skin and adipose tissues. In some cases, the cis-eQTL signal — identified with males and females combined — seems to be driven exclusively by one sex. For example, the *cis*-eQTL association of rs2273535 with the gene *AURKA* in skeletal muscle (cis-eQTL p = 6.92×10^24^) is correlated with sex (p_G×S_ = 9.28×10^−12^, Storey q_G×S_ = 1.07×10^−7^, AE validation p = 1.15×10^−11^) and present only in males (figs. 2d, S12). *AURKA* is a member of the serine/threonine kinase family involved in mitotic chromosomal segregation that has been widely studied as a risk factor in several cancers (*24-27*) and has been recently shown to be involved in muscle differentiation (*28*).

We also characterized population-biased *cis*-eQTLs (pb-eQTLs), where a variant’s molecular effect on gene expression differs between individuals of European and African ancestry, controlling for differences in allele frequency and Linkage Disequilibrium (LD) (*11*). Analyzing 31 tissues with sample sizes >20 in both populations, we mapped genes with a different eQTL effect size measured by aFC. After applying stringent filters to remove differences potentially explained by LD or other artifacts (fig. S13a), we identified 178 pb-eQTLs for 141 eGenes (FDR ≤ 25%) that show a moderate degree of validation in allele-specific expression data (fig. S13, table S10). While some of the pb-eQTL effects are tissue-specific, there are also effects that are shared across most tissues (fig. S13). Figure 2e shows an example of a pb-eQTL for the *SLC44A5* gene involved in transport of sugars and amino acids, and expressed at different levels between epidermis of lighter and darker skin (reconstructed *in vitro*) (*29, 30*). In Europeans, the derived allele of rs4606268 decreases expression of the gene in esophagus mucosa (aFC = −4.82), but this effect is significantly lower in African Americans (aFC = −2.85, permutation p-value = 1.2×10^−3^, AE validation p = 0.002, fig. S13)

This relative paucity of both sex- and population-biased *cis*-eQTLs reflects the fact that they are challenging to identify and there are few with large effects, but that they can provide insights in to sex- or population-specific regulatory effects on gene expression.

### Fine-mapping

A major challenge of all genetic association studies is to distinguish the causal variants from their LD proxies. We applied three different statistical fine-mapping methods — CaVEMaN (*31*), CAVIAR (*32*), and dap-g (*16*) — to infer likely causal variants of *cis*-eQTLs in each tissue (fig. 3a) (*11*). For many *cis*-eQTLs the causal variant can be mapped with a high probability to a handful of candidates: the 90% credible set for each *cis*-eQTL consists of variants that include the causal variant with 90% probability; using dap-g, we identified a median of 6 variants in the 90% credible set for each *cis*-eQTL (fig. S14). Furthermore, 9.3% of the *cis*-eQTLs have a variant with a posterior probability > 0.8 according to dap-g, indicating a single likely causal variant for those *cis*-eQTLs. We defined a consensus set of 24,740 *cis*-eQTLs across all tissues (7,709 unique variants), for which the posterior probability was >0.8 across all three methods (fig. S15). Fine-mapped variants were significantly higher enriched among experimentally validated causal variants from MPRA (*33*) and SuRE (*34*) data, compared to the lead eVariant across all eGenes. The highest enrichment was observed for the consensus set although with overlapping confidence intervals (fig. 3b). This demonstrates how careful fine-mapping facilitates the identification of likely causal regulatory variants.

**Figure 3.**
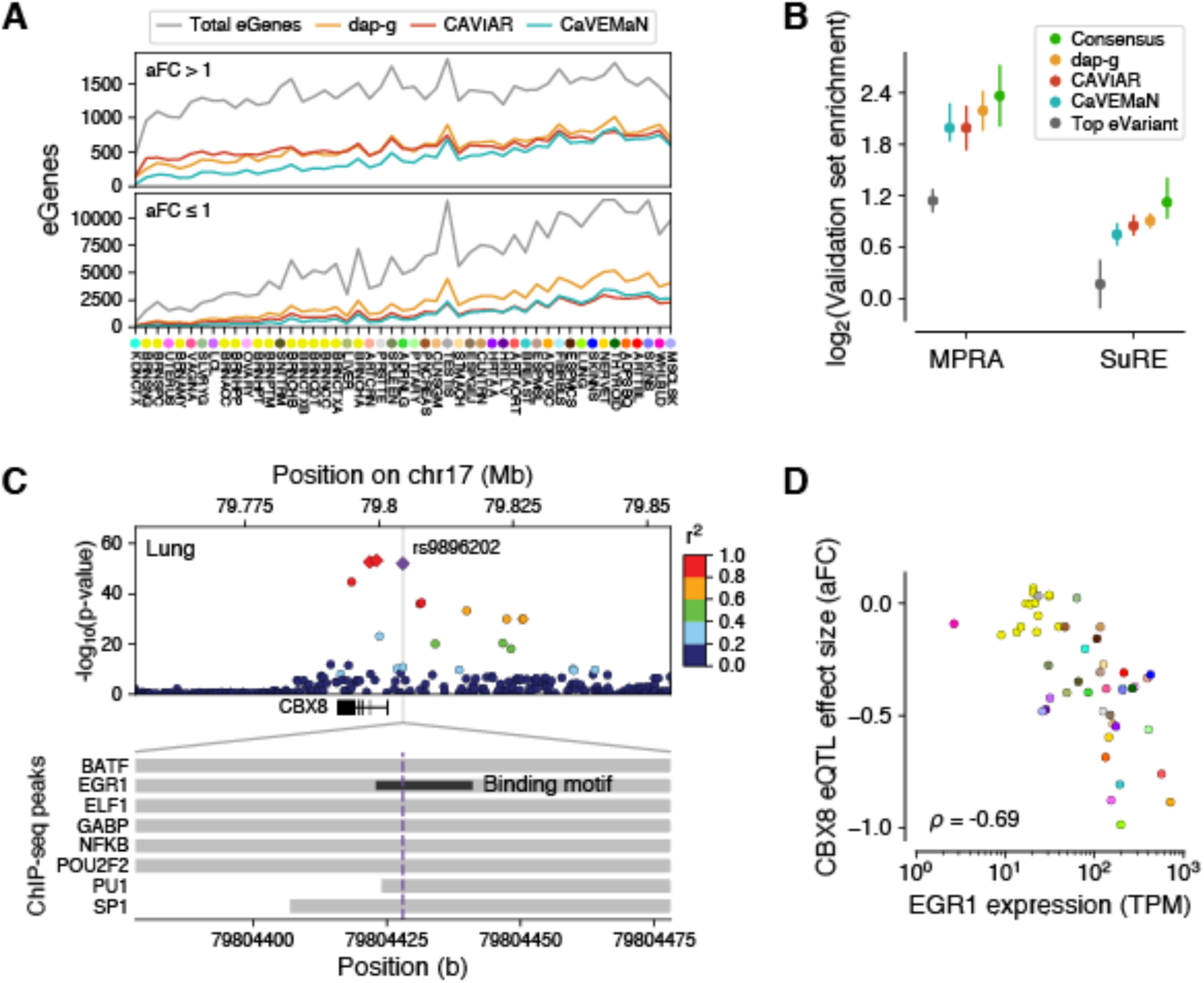
Fine mapping of *cis*-eQTLs. **(A)** Number of eGenes per tissue with variants fine-mapped with >0.5 posterior probability of causality, based on three methods. The overall number of eGenes with at least one fine-mapped eVariant increases with sample size for all methods. However, this increase is in part driven by better statistical power to detect small effect size *cis*-eQTLs (aFC or allelic fold change ≤1 in log2 scale) with larger sample sizes, and the proportion of well fine-mapped eGenes with small effect sizes increases more modestly with sample size (bottom vs. top panels), indicating that such *cis*-eQTLs are generally more difficult to fine-map. (**B)** Enrichment of variants among experimentally validated regulatory variants, shown for the *cis*-eVariant with the best p-value (top eVariant), and those with posterior probability of causality >0.8 according to each of the three methods individually or all of them (consensus). Error bars: 95% CI **(C**) The *cis*-eQTL signal for *CBX8* is fine-mapped to a credible set of three variants (red and purple diamonds), of which rs9896202 (purple diamond) overlaps a large number of transcription factor binding sites in ENCODE ChIP-seq data and disrupts the binding motif of *EGR1*. (**D**) The potential role of EGR1 binding driving this *cis*-eQTL is further supported by correlation between *EGR1* expression and the *CBX8 cis*-eQTL effect size across tissues.

Knowing the likely causal variant enables greater insights into the molecular mechanisms of individual eQTLs, including the mechanisms of their tissue-specific effects. Figure 3c shows an example of an eQTL for the gene *CBX8* that colocalizes with breast cancer risk and birth weight (posterior probability 0.68 for both in lung). One of the three variants in the confident set overlaps the binding site and disrupts the motif of the transcription factor *EGR1* (*1*) (fig. S16). The role of *EGR1* as an upstream driver of this eQTL is further supported by a cross-tissue correlation of the effect size of the eQTL and the expression level of *EGR1* (Spearman’s rho = −0.69) (fig. 3d).

### Functional mechanisms of QTL associations

Quantitative trait data from multiple molecular phenotypes, integrated with the regulatory annotation of the genome and GWAS data (table S3), offer a powerful way to understand the molecular mechanisms and phenotypic consequences of genetic regulatory effects. As expected, *cis*-eQTLs and *cis*-sQTLs are significantly enriched in functional elements of the genome (fig. 4a). While the strongest enrichments are driven by variant classes that lead to splicing changes or nonsense-mediated decay, these account for relatively few variants. *Cis*-sQTLs have significant enrichments almost entirely in transcribed regions, while *cis*-eQTLs are significantly enriched in transcriptional regulatory elements as well. Previous studies (*4, 35*) have indicated that *cis*-eQTL and *cis*-sQTL effects on the same gene are typically driven by different genetic variants. This is corroborated by the GTEx v8 data, where the overlap of *cis*-eQTL credible sets of likely causal variants, based on CAVIAR, have only a 12% overlap with *cis*-sQTL credible sets (fig. S17). Functional enrichment of overlapping and non-overlapping *cis-*eQTLs and *cis-*sQTLs, based on stringent LD filtering, showed that the patterns characteristic for each type — such as enrichment of *cis*-eQTL in enhancers and *cis*-sQTLs in splice sites — are even stronger for distinct loci (fig. S17).

**Figure 4.**
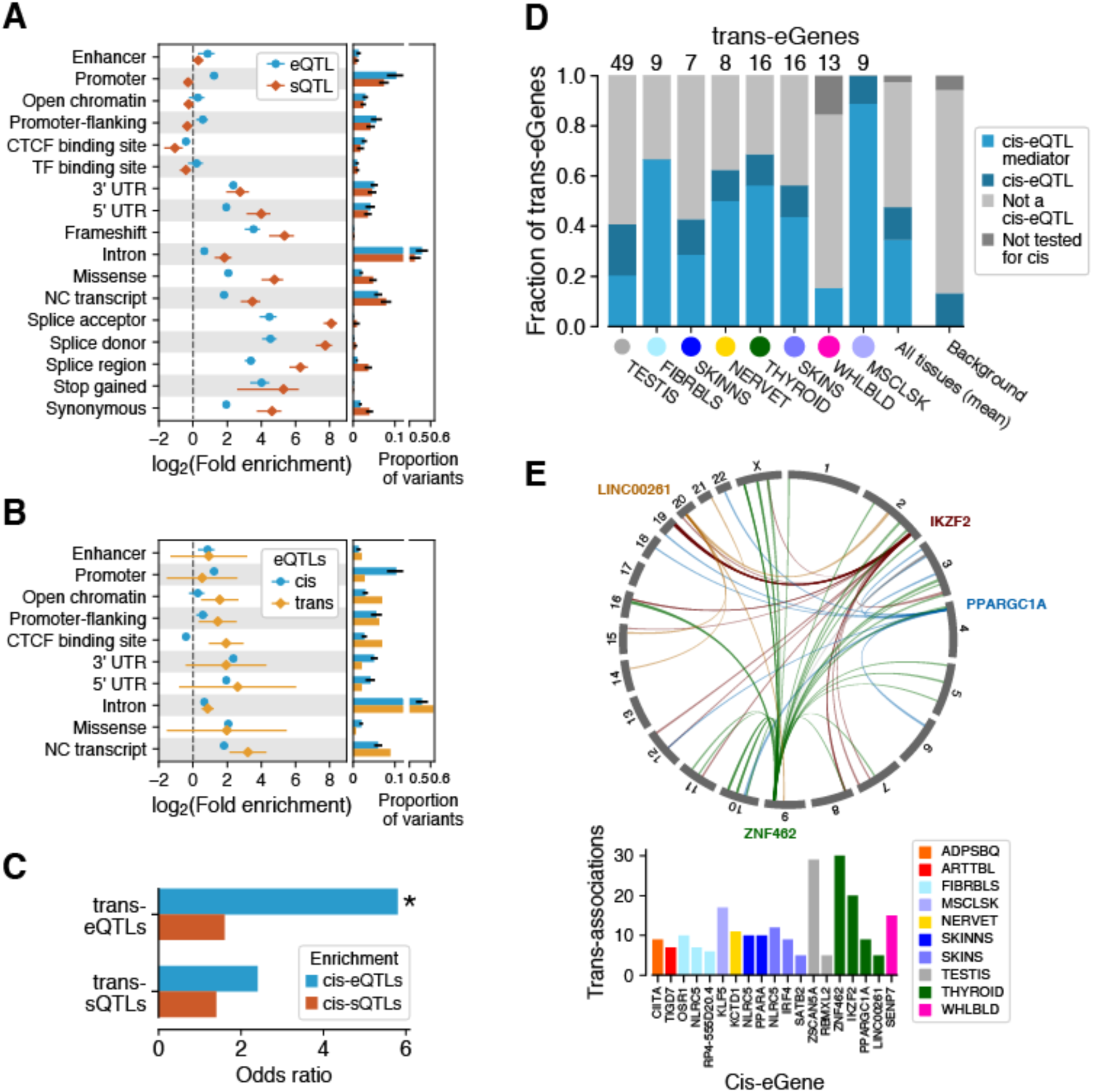
Functional mechanisms of genetic regulatory effects. QTL enrichment in functional annotations for (**A**) *cis*-eQTLs and *cis*-sQTLs and for (**B**) *trans*-eQTLs. *cis*-QTL enrichment is shown as mean ± s.d. across tissues; *trans*-eQTL enrichment as 95% C.I. **(C)** Enrichment of lead *trans*-e/sVariants tested in *cis* being also *cis*-e/sVariants in the same tissue. * denotes significant enrichment, p < 10^−21^. (**D**) Proportion of *trans*-eQTLs that are significant *cis*-eQTLs or mediated by *cis*-eQTLs. **(E)** Trans associations of cis-mediating genes identified through colocalization (PP4 > 0.8 and nominal association with discovery trans-eVariant p < 10^−5^). Top: associations for four Thyroid *cis*-eQTLs (indicated by gene names); bottom: *cis*-mediating genes with ≥5 colocalizing *trans*-eQTLs.

We hypothesized that eVariants and their target eGenes in *cis* are more likely to be in the same topologically associated domains (TADs) that allow chromatin interactions between more distant regulatory regions and target gene promoters (*36*). To test this, we analyzed TAD data from ENCODE (*1*) and *cis*-eQTLs from matching GTEx tissues (table S3). Compared to matching random variant-gene pairs and controlling for distance from the transcription start site, *cis*-eVariant-eGene pairs were significantly enriched for being in the same TAD (median log odds 1.52; all p<10^−12^) (fig.S18).

*Trans*-eQTLs are significantly enriched in regulatory annotations that suggest both pre- and post-transcriptional mechanisms (fig. 4b). Unlike *cis*-eQTLs, *trans*-eQTLs are strongly enriched in CTCF binding sites, suggesting that disruption of CTCF binding may underlie distal genetic regulatory effects, potentially via its effect on interchromosomal chromatin interactions (*36*). *trans*-eQTLs have also been shown to be partially driven by *cis*-eQTLs (*37, 38*). Indeed, we observed a significant enrichment of lead *trans*-eVariants tested in *cis* being also *cis*-eVariants in the same tissue (5.9x; two-sided Fisher’s exact test p = 5.03×10^−22^, fig. 4c). Lack of analogous strong enrichment suggests that *cis*-sQTLs are less important contributors to *trans*-eQTLs (p = 0.064), and *trans*-sVariants had no significant enrichment of either *cis*-eQTLs (p = 0.051) or *cis*-sQTLs (p = 0.53). A further demonstration of the important contribution of *cis*-eQTLs to *trans*-eQTLs is that, based on mediation analysis, 77% of lead *trans*-eVariants that are also *cis*-eVariants appear to act through the *cis*-eQTL (figs. 4d, S19). Colocalization of *ci*s-eQTLs and *trans*-eQTLs was widespread and often tissue-specific, with figure 4e showing *cis*-eQTLs with at least ten nominally significant colocalized *trans*-eQTLs each (PP4 > 0.8 and *trans*-eQTL p-value < 10^−5^), pinpointing how local effects on gene expression can potentially lead to downstream regulatory effects across the genome (fig. S20, table S15).

### Genetic regulatory effects mediate complex trait associations

In order to analyze the role of regulatory variants in genetic associations for human traits, we first asked whether variants in the GWAS catalog were enriched for significant QTLs, compared to all variants tested for QTLs (*11*). We observed a 1.46-fold enrichment for *cis*-eQTLs (63% vs 43%) and 1.86-fold enrichment for *cis*-sQTLs (37% vs 20%). The enrichment was even stronger, 6.97-fold (0.029% vs 0.0042%) for *trans*-eQTLs, consistent with other analyses (*39*) (figs. 5a, S21-22, tables S5-6).

**Figure 5.**
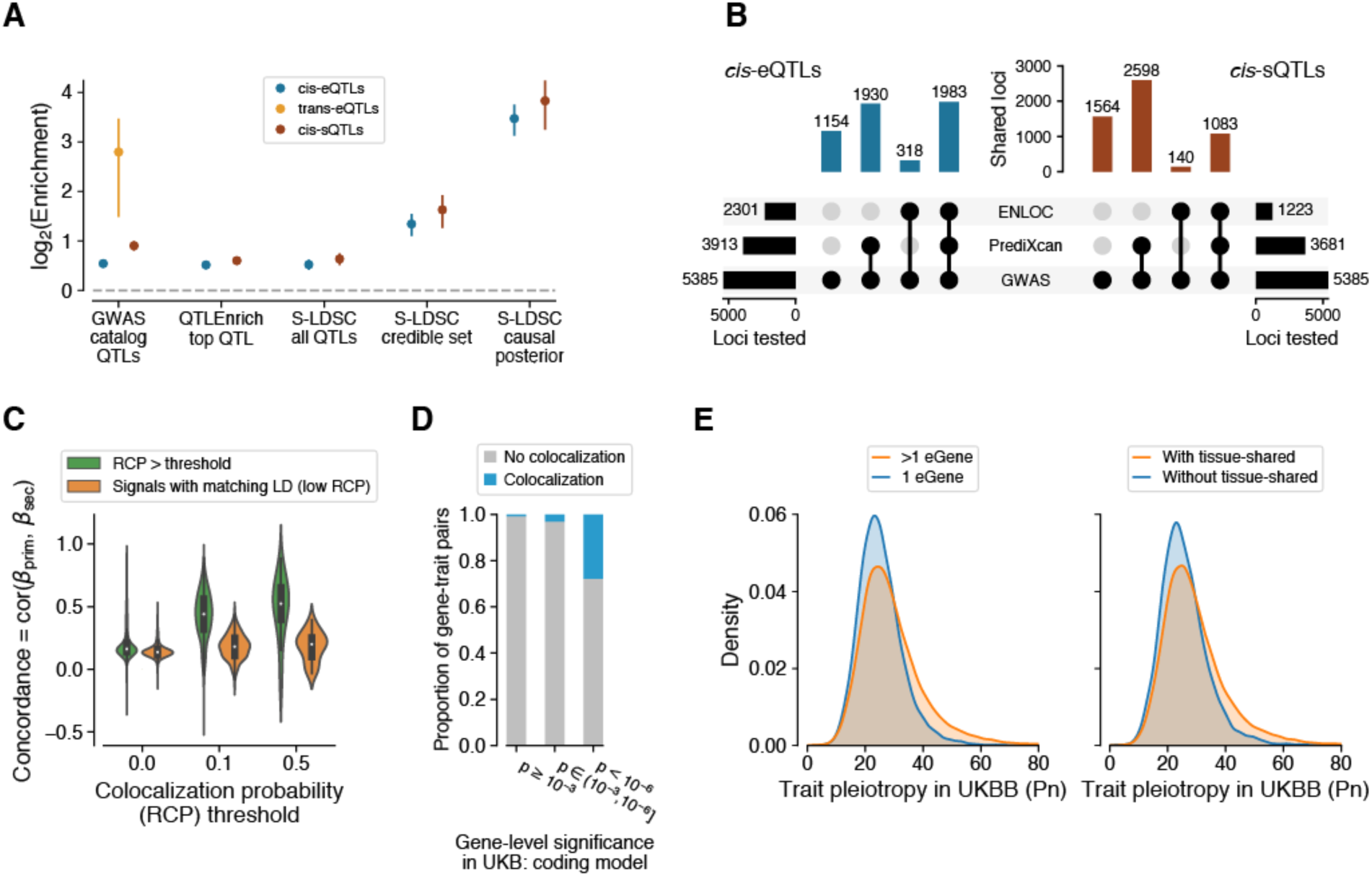
Regulatory mechanisms of GWAS loci. **(A)** GWAS enrichment of *cis*-eQTLs, *cis*-sQTLs, and *trans*-eQTLs measured with different approaches: enrichment based on GWAS summary statistics of the most significant *cis*-QTL per eGene/sGene with QTLEnrich and LD Score regression with all significant *cis*-QTLs (S-LDSC all *cis*-QTLs), simple QTL overlap enrichment with all GWAS catalog variants, and LD Score regression with fine-mapped *cis*-QTLs in the 95% credible set (S-LDSC credible set) and using posterior probability of causality as a continuous annotation (S-LDSC causal posterior). Enrichment is shown as mean and 95% CI. **(B)** Number of GWAS loci linked to e/sGenes through colocalization (ENLOC) and association (PrediXcan), aggregated across tissues. (**C)** Concordance of mediated effects among independent *cis*-eQTLs for the same gene is shown for different levels of colocalization probability, which is used as a proxy for the gene’s causality. As the null, we show the concordance for LD matched genes without colocalization. (**D)** Proportion of colocalized *cis*-eQTLs with a matching phenotype for genes with different level of rare variant trait association in the UK Biobank. (**E)** Horizontal GWAS trait pleiotropy score distribution for *cis*-eQTLs that regulate multiple vs. a single gene (left), and for *cis*-eQTLs that are tissue-shared vs specific.

This approach does not leverage the full power of genome-wide GWAS and QTL genetic association statistics, nor account for LD contamination, a situation wherein the causal variants for QTL and GWAS signals are distinct but LD between the two causal variants can suggest a false functional link (*40*). Hence, for subsequent analyses (below) we selected 87 Genome Wide Association Studies (GWAS) representing a broad array of binary and continuous complex traits that have summary results available in the public domain (*11, 41*) (tables S4, S11), and *cis*-QTL statistics calculated from the European subset of GTEx donors to match the ancestry of GWAS studies (fig. S24). Analyses described were performed for all pairwise combinations of 87 phenotypes and 49 tissues, and are summarized using an approach that accounts for similarity between tissues and variable standard errors of the QTL effect estimates, driven mainly by tissue sample size (fig. S22, (*11*)).

To analyze the mediating role of *cis*-regulation of gene expression on complex traits (*35, 42*), we used two complementary approaches, QTLEnrich (*43*) and Stratified LD score regression (S-LDSC) (*11*). To rule out the possibility that enrichment is driven by specific features of *cis*-QTLs such as allele frequency, distance to the transcription start site, or local level of LD (number of LD proxy variants; r^2^ ≥ 0.5), we used QTLEnrich. We found a 1.43-fold (SE=0.04) and 1.52-fold (SE=0.04) enrichment of trait associations among best *cis*-eQTLs and *cis*-sQTLs, respectively, adjusting for enrichment among matched null variants (fig. 5a, tables S7). The fact that these enrichment estimates differ little from those derived from the GWAS catalog overlap (above), even after accounting for the potential confounders, indicates how relatively robust these estimates are. Next, we used S-LDSC adjusting for functional annotations (*44*) to confirm the robustness of these results and to analyze how GWAS enrichment is affected by the causal e/sVariant being typically unknown (*11*). We computed the heritability enrichment of all *cis*-QTLs, fine-mapped *cis*-QTLs (in 95% credible set and posterior probability > 0.01 from dap-g), and fine-mapped *cis*-QTLs with maximum posterior inclusion probability as continuous annotation (MaxCPP) (*45*) (fig. 5a). The largest increase in GWAS enrichment was for likely causal *cis*-QTL variants (11.1-fold (SE=1.2) for *cis*-eQTLs and 14.2-fold (SE=2.4) for and *cis*-sQTLs, for the continuos annotation), which is strong evidence of shared causal effects of *cis*-QTLs and GWAS, and for the importance of fine-mapping.

Joint enrichment analysis of *cis*-eQTLs and *cis*-sQTLs shows an independent contribution to complex trait variation from both (fig. S23, (*11*)), consistent with their limited overlap (fig. S17). The relative GWAS enrichments of *cis*-sQTLs and *cis-*eQTLs were similar (fig. 5a; not significant for the robust QTLEnrich and LDSC analyses), but the larger number of *cis*-eQTLs discovered (fig. 2a) suggests a greater aggregated contribution of *cis*-eQTLs.

To provide functional interpretation of the 5,385 significant GWAS associations in 1,167 loci from approximately independent LD blocks (*46*) across the 87 complex traits, we performed colocalization with *enloc* (*16*) to quantify the probability that the *cis*-QTL and GWAS signals share the same causal variant. We also assessed the association between the genetically regulated component of expression or splicing and complex traits with PrediXcan (*11, 41, 47*). Both methods take multiple independent *cis*-QTLs into account, which is critical in large *cis*-eQTL studies such as GTEx with widespread allelic heterogeneity. Of the 5,385 GWAS loci, 43% and 23% were colocalized with a *cis*-eQTL and *cis*-sQTL, respectively (fig. 5b). A large proportion of colocalized genes coincide with significant PrediXcan trait associations with predicted expression or splicing (median of 86% and 88% across phenotypes respectively, figs. S25-S28, tables S8). Together, these results suggest target genes and their potential molecular changes for thousands of GWAS loci.

Having multiple independent *cis*-eQTLs for a large number of genes allowed us to test whether mediated effects of primary and secondary *cis*-eQTLs on phenotypes — the ratio of GWAS and *cis*-eQTL effect sizes — are concordant. To make sure that concordance is not driven by residual LD between primary and secondary signals, we used LD-matched *cis*-eGenes with low colocalization probability as controls (*11, 41*), and observed a significant increase in primary and secondary *cis*-eQTL concordance for colocalized genes (p-value < 10^−30^; fig. 5c). Additionally, colocalization of a *cis*-eQTL increased the colocalization of an independent *cis*-sQTL in the same locus (OR = 4.27, p < 10^−16^), and correspondingly colocalization of a *cis*-sQTL increased *cis*-eQTL colocalization (OR = 4.54 p < 10^−16^; fig. S29). This indicates that multiple regulatory effects for the same gene often mediate the same complex trait associations. Furthermore, genes with suggestive rare variant trait associations in the UK Biobank (*48*) have a substantially increased proportion of colocalized eQTLs for the same trait (fig. 5d), showing concordant trait effects from rare coding and common regulatory variants (*49*). These genes, as well as those with multiple colocalizing *cis*-QTLs, represent bona fide disease genes with multiple independent lines of evidence.

The growing number of genome and phenome studies has revealed extensive pleiotropy, where the same variant or locus associates with multiple organismal phenotypes (*50*). We sought to analyze how this phenomenon can be driven by gene regulatory effects. First, we calculated the number of *cis*-eGenes of each fine-mapped and LD-pruned *cis*-eVariant per tissue at local LFSR < 5%, with cross-tissue smoothing of effect sizes with *mashr* (*11, 51*). We observed that a median of 57% of variants were associated with more than one gene per tissue, typically co-occurring across tissues, indicating widespread regulatory pleiotropy. Using a binary classification of *cis*-eVariants with regulatory pleiotropy defined as those associated with more than one gene, we observed that they are more significantly associated with complex traits compared to matched *cis*-eVariants (fig. S30). This could be due to the fact that if a variant regulates multiple genes, there is a higher probability that at least one of them affects a GWAS phenotype. However, *cis*-eVariants with regulatory pleiotropy also have higher GWAS complex trait pleiotropy (*50*) than *cis*-eVariants with effects on a single gene (fig. 5e). This observation suggests a mechanism for complex trait pleiotropy of genetic effects where the expression of multiple genes in *cis*, rather than a single eGene effect, translates into diverse downstream physiological effects. Furthermore, GWAS pleiotropy is higher for tissue-shared (*41*) than tissue-specific *cis*-eQTLs, indicating that regulatory effects affecting multiple tissues are more likely to translate to diverse physiological traits (fig. 5e).

*Cis-* and *trans*-eQTLs can provide insights into potential mechanisms and effects of trait-associated variants. In one such example, rs1775555 on chr10p14 is a fibroblast-specific *cis*-eQTL for *GATA3* (p=7.4×10^−70^) and a lincRNA gene *GATA3-AS1* (p=1.8×10^−45^) and a *trans*-eQTL for *MSTN* on chromosome 2, which encodes a TGF-β ligand secreted protein (fig. S31) and has a role in muscle growth and also the immune system (*52*). *GATA3* is a transcription factor known to regulate a range of processes of immune system including T cell development, Th2 differentiation, and immune cell homeostasis and survival (*53*). The *cis-* (*GATA3*) and *trans*-eQTL (*MSTN*) associations colocalized (PP4 > 0.99) in fibroblasts, and mediation analysis supports that the effect of rs1775555 on *MSTN* is mediated through GATA3 (p=2.1×10^−22^, (*11*)). We also found that the *cis-* and *trans*-eQTL effect of rs1775555 colocalized with associations for multiple immune traits, including combined eosinophil and basophil counts, hayfever/eczema, and asthma (PP4 > 0.97 for all eQTL-trait combinations; fig. S31). *DTNA, C4orf26, GK5, HSD11B1, SLC44A1, ARHGAP25, MAN2A1* are additional genes that showed *trans* association with this variant (FDR 10%, corrected for number of cross-chromosomal genes tested for association with rs1775555). While the causal relationships are not obvious, this locus demonstrates broad impact on multiple phenotypes and both local and distal gene expression.

### Tissue-specificity of genetic regulatory effects

The GTEx data provide a unique opportunity to study patterns and mechanisms of tissue-specificity of the transcriptome and its genetic regulation. Pairwise similarity of GTEx tissues was quantified using gene expression and splicing, as well as allelic expression, eQTLs in *cis* and *trans*, and *cis*-sQTLs (figs. 6a, S34, (*11*)). These show highly consistent patterns of tissue relatedness, indicating that the same biological processes that drive transcriptome similarity also control tissue sharing of genetic effects (fig. 6b). As seen in earlier versions of the GTEx data (*9, 10*), the brain regions form a separate cluster, and testis, LCLs, whole blood, and sometimes liver tend to be outliers, while most other organs have a notably high degree of similarity between each other. This indicates that blood is far from an ideal proxy for most tissues, but that some other relatively accessible tissues, such as skin, may be better at capturing molecular effects in other tissues.

**Figure 6.**
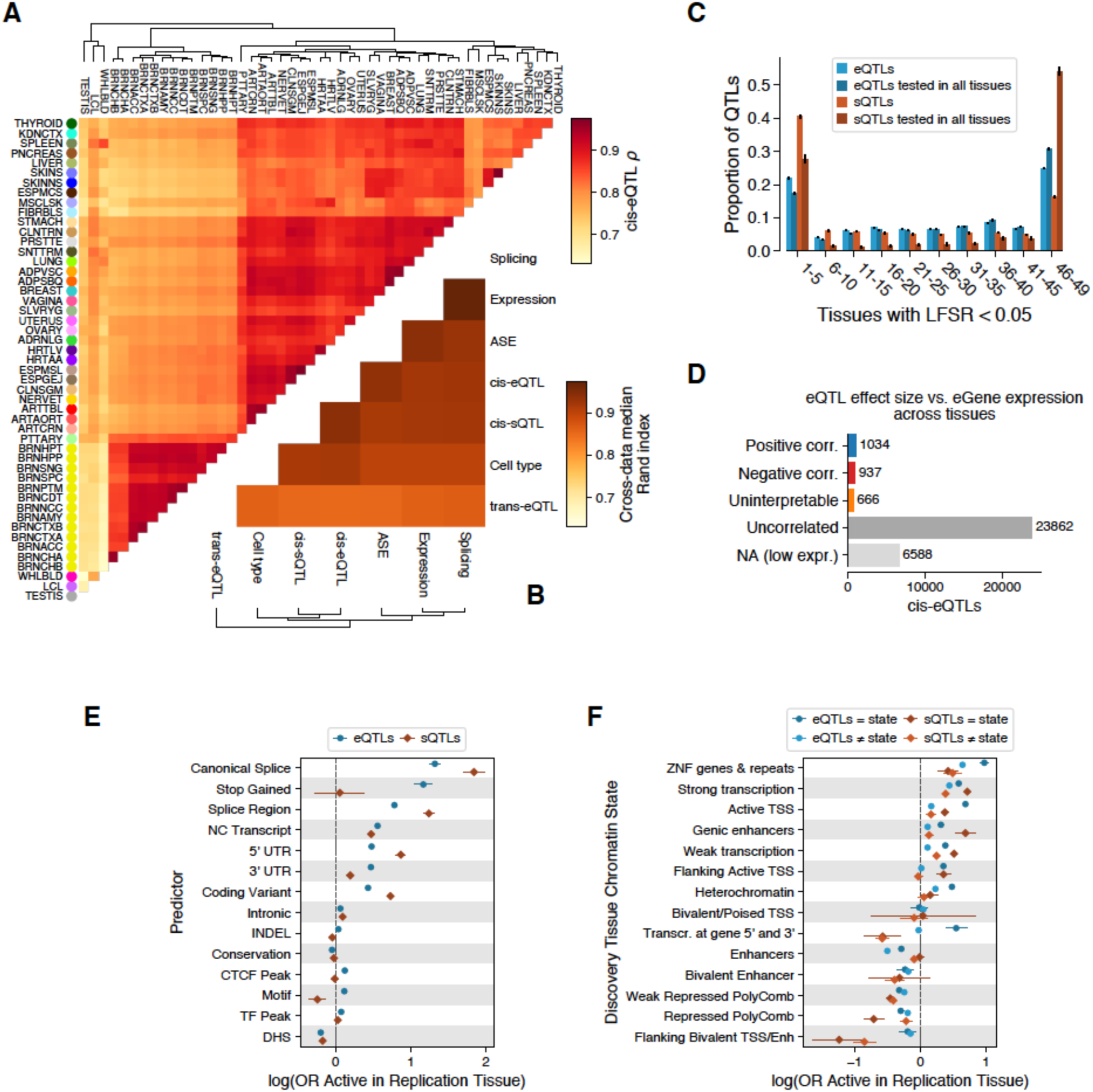
Tissue-specificity of *cis-*QTLs. (**A)** Tissue clustering based on pairwise Spearman correlation of *cis*-eQTL effect sizes. (**B)** Similarity of tissue clustering across core data types quantified using median pairwise Rand index calculated across tissues. (**C)** Tissue activity of *cis* expression and splicing QTLs, where an eQTL was considered active in a tissue if it had a *mashr* local false sign rate (LFSR, equivalent to FDR) of < 5%. This is shown for all *cis*-QTLs and only those that could be tested in all 49 tissues (red and blue). **(D)** Spearman correlation (corr.) between *cis*-eQTL effect size and eGene expression level across tissues. *cis*-eQTL counts are shown for those not tested due to low expression level, tested but without significant (FDR < 5%) correlation (uncorrelated), a significant correlation but effect sizes crossed zero which made the correlation direction unclear (uninterpretable), positively correlated, and negatively correlated. (**E-F)** The effect of genomic function on *cis-*QTL tissue sharing modeled using logistic regression, using functional annotations (**E**) and chromatin state (**F**). CTCF Peak, Motif, TF Peak, and DHS indicate if the *cis*-QTL lies in a region annotated as having one of these features in any of the Ensembl Regulatory Build tissues. For chromatin states, model coefficients are shown for the discovery and replication tissues that have the same or different chromatin states.

The overall tissue specificity of QTLs (*11*) follows a U-shaped curve recapitulating previous GTEx analyses (*9, 10*), where genetic regulatory effects tend to be either highly tissue-specific or highly shared (fig. 6c), with *trans*-eQTLs being more tissue-specific than *cis*-eQTLs (fig. S33). *Cis*-sQTLs appear to be significantly more tissue specific than *cis*-eQTLs when considering all mapped *cis*-QTLs, but this pattern is reversed when considering only those *cis*-QTLs where the gene or splicing event is quantified in all tissues (figs. 6c, S32). This indicates that splicing measures are more tissue-specific than gene expression, but genetic effects on splicing tend to be more shared, consistent with pairwise tissue sharing patterns (fig. S34). This is important for understanding effects that disease-causing splicing variants may have across tissues, and for validation of splicing effects in cell lines that rarely are an exact match to cells *in vivo*. Next, we analyzed the sharing of allelic expression (AE) across multiple tissues of an individual, which is a sensitive metric of sharing of any heterozygous regulatory variant effects in that individual and has been particularly useful for analysis of rare, potentially disease-causing variants (*54*). Using a clustering approach (*11*), we found that in 97.4% of the cases, AE across all tissues forms a single cluster. This suggests that in AE analysis, different tissues are often relatively good proxies for one another, provided that the gene of interest is expressed in the probed tissue (fig. S35).

We next computed the cross-tissue correlation of eQTL effect size and eGene expression level — often a proxy for gene functionality — and discovered that 1,971 *cis*-eQTLs (7.4%; FDR 5%) had a significant and robust correlation between eGene expression and *cis*-eQTL effect size across tissues (fig. 6d, S36). These correlated *cis*-eQTLs are split nearly evenly between negative (937) and positive (1,034) correlations. Thus, the tissues with the highest *cis*-eQTL effect sizes are equally likely to be among tissues with higher or lower expression levels for the gene. *Trans*-eQTLs show a different pattern, being typically observed in tissues with high expression of the *trans*-eGene relative to other tissues (fig. S36). These observations raise the question how to prioritize the relevant tissues for eQTLs in a disease context. To address this, we chose a subset of GWAS traits where previous studies provide a strong prior for the likely relevant tissue(s) (table S12). Analyzing colocalized *cis*-eQTLs for 1,778 GWAS loci (*11*), we discovered that the relevant tissues were modestly but significantly enriched in having high expression and effect sizes (p<1.5×10^−4^) (figs. S37-38, table S9). This indicates that both effect size and gene expression level are important in the interpretation of the tissue context where an eQTL may have downstream phenotypic effects.

The diverse patterns of QTL tissue-specificity raise the question of what molecular mechanisms underlie the ubiquitous regulatory effects of some genetic variants and the highly tissue-specific effects of others. To gain insight into this question, we modeled *cis*-eQTL and *cis*-sQTL tissue specificity using logistic regression as a function of the lead eVariant’s genomic and epigenomic context (*11*). *Cis*-QTLs where the top eVariant was in a transcribed region had overall higher sharing than those in classical transcriptional regulatory elements, indicating that genetic variants with post- or co-transcriptional expression or splicing effects have more ubiquitous effects (fig. 6e). Canonical splice and stop gained variant effects had the highest probability of being shared across tissues, which may benefit disease-focused studies relying on likely gene-disrupting variants. We also considered whether varying regulatory activity between tissues contributed to tissue-specificity of genetic effects, and found that shared chromatin state between the discovery and query tissues was associated with increased probability of *cis*-eQTL sharing and vice-versa (fig. 6f). *cis*-eQTLs and *cis*-sQTLs followed similar patterns. Since *cis*-sQTLs are more enriched in transcribed regions and likely post-transcriptional mechanisms (fig. 4a), this is likely to contribute to their higher overall degree of tissue-sharing (fig. 6c). In comparison to *cis*-eQTLs, *cis*-sQTLs are indeed more often located in regions where regulatory effects are shared. These data offer a possibility to predict if an *cis*-eQTL observed in a GTEx tissue is active in another tissue of interest, based on its annotation and properties in the discovery tissue (*11*). After incorporating additional features including *cis*-QTL effect size, distance to transcription start site, and eGene/sGene expression levels, we obtain reasonably good predictions of whether a *cis*-QTL is active in a query tissue (median AUC = 0.779 and 0.807, min = 0.703 and 0.721, max = 0.807 and 0.875 for *cis*-eQTLs and *cis*-sQTLs, respectively; fig. S39). This suggests that it is possible to extrapolate the GTEx *cis*-eQTL catalog to additional tissues or, for example, developmental stages where population-scale data for QTL analysis are particularly difficult to collect.

### From tissues to cell types

The GTEx tissue samples consist of heterogeneous mixtures of multiple cell types. Hence, the RNA extracted and QTLs mapped from these samples reflect a composite of effects that may vary across cell types and may mask cell type-specific mechanisms. To characterize the effect of cell type heterogeneity on analyses from bulk tissue, we used the xCell method (*55*) to estimate the enrichment of 64 reference cell types from the bulk expression profile of each sample (*11*). The resulting enrichment scores were generally biologically meaningful, with for example myocytes enriched in heart left ventricle and skeletal muscle, hepatocytes enriched in liver, and various blood cell types enriched in whole blood, spleen, and lung, which is known to harbor a large leukocyte population (fig. S40). As discussed in more detail in (*56*), these results need to be interpreted with caution given the scarcity of validation data and quality and quantity of cell type reference data sets. Nonetheless, the pairwise relatedness of GTEx tissues derived from their cell type composition is highly correlated with tissue-sharing of regulatory variants (figs. 5b, S41, S34), suggesting similarity of regulatory variant activity between tissue pairs may often be due to the presence of similar cell types, and not necessarily shared regulatory networks within cells. This highlights the key role that characterizing cell type diversity will have, not only for understanding tissue biology, but the underlying role of genetic variation as well.

Enrichment of many cell types shows inter-individual variation within a given tissue (*56*). In eQTL analysis, this variation can be leveraged to identify *cis*-eQTLs and *cis*-sQTLs with cell type specificity by extending the QTL model to include an interaction between genotype and cell type enrichment (*11, 57*). We applied this approach to seven tissue-cell type pairs that were chosen based on having robustly quantified cell types and the tissue where each cell type was most enriched (fig. 7a; an additional 36 pairs are described in (*56*)). Power to discover cell type interacting *cis*-eQTLs and *cis-*sQTLs (ieQTLs and isQTLs, respectively) varied as a function of tissue heterogeneity and complexity as well as sample size (*56*). We notably identified 1120 neutrophil ieQTLs in whole blood and 1087 epithelial cell ieQTLs in transverse colon (fig. 1a); of these, 76 and 229 respectively, involved an eGene for which no QTL was detected in bulk tissue. eQTLs from purified neutrophils of an external data set (*58*) had higher neutrophil ieQTL effect sizes than eQTLs from other blood cell types (fig. S42). For other cell types external replication data was lacking. Thus, we verified the robustness of the ieQTLs by the allelic expression validation approach that was used for sex- and population-biased *cis*-eQTL analyses: for ieQTL heterozygotes, we calculated the Spearman correlation of cell type enrichment and ieQTL effect size from AE data, and observed a high validation rate (*56*). It is important to note that ie/isQTLs should not be considered cell type-specific QTLs, because the enrichment of any cell type may be (anti-)correlated with other cell types (fig. S43). While full deconvolution of *cis*-eQTL effects driven by specific cell types remains a challenge for the future, ieQTLs and isQTLs can be interpreted as being enriched for cell type-specific effects. In most subsequent analyses to characterize the properties of ieQTLs and isQTLs, we focused on the neutrophil ieQTLs, which are numerous and supported by external replication data.

**Figure 7.**
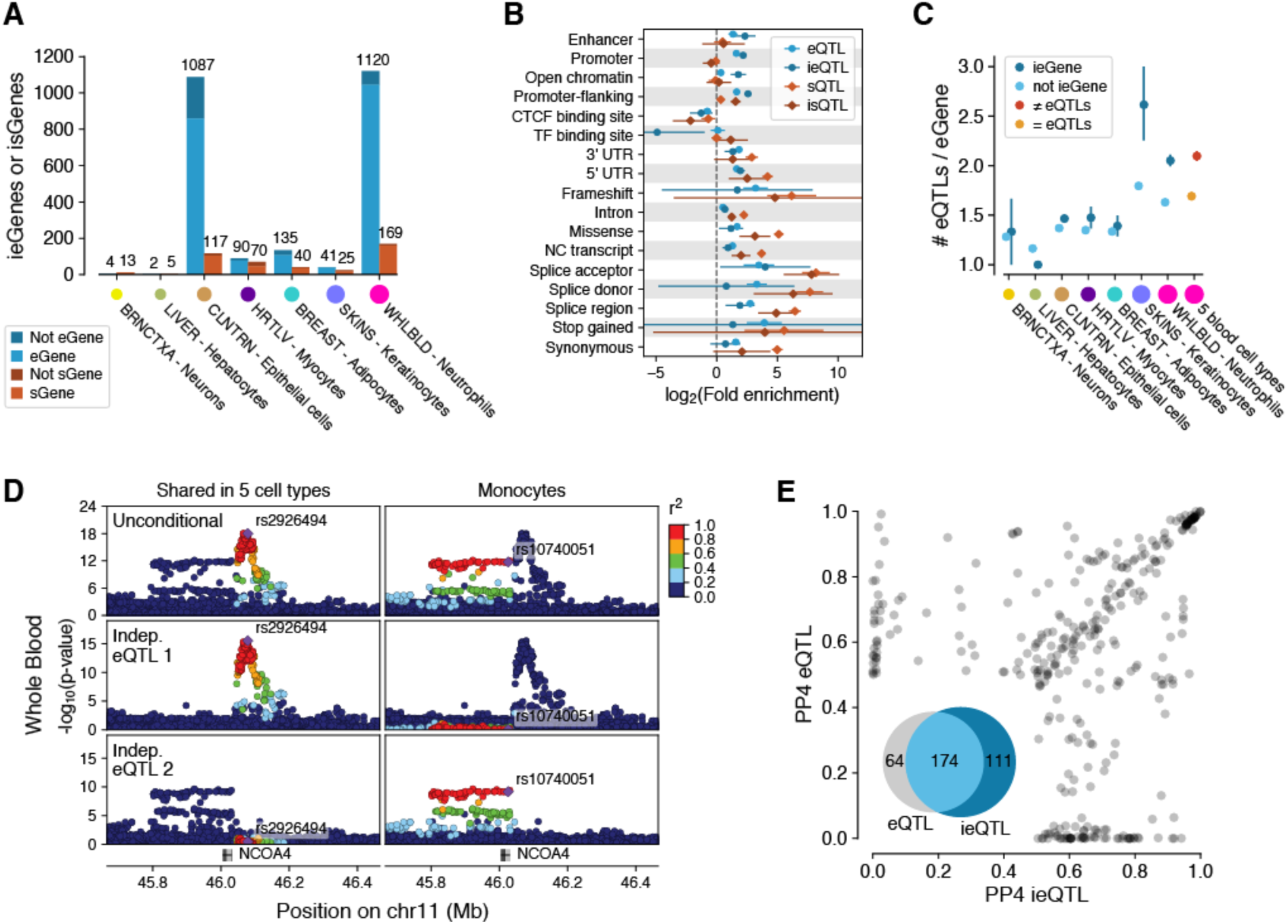
Cell type interacting *cis*-eQTLs and *cis*-sQTLs. (**A**) Number of cell type interacting *cis-*eQTLs and *cis-* sQTLs (ieQTLs and isQTLs, respectively) discovered in seven tissue-cell type pairs, with shading indicating whether the ieGene or isGene was discovered by *cis*-eQTL/*cis*-sQTL analysis in bulk tissue. Colored dots are proportional to sample size. (**B**) Functional enrichment of neutrophil ieQTLs and isQTLs compared to *cis*-eQTLs and *cis*-sQTLs from whole blood. (**C**) Proportion of conditionally independent *cis*-eQTLs per eGene, for eGenes that do or do not have ieQTLs in GTEx, and for eGenes that have shared (= eQTLs) or non-shared (≠ eQTLs) *cis-*eQTL across five sorted blood cell types. (**D**) Whole blood *cis-*eQTL p-value landscape for *NCOA4*, for the standard analysis (top row, Unconditional) and for two independent *cis-*eQTLs (bottom rows). In a data set of 5 sorted cell types (*58*), analyses of all cell types yielded a lead eVariant, rs2926494 (left), which is in high LD with the first independent *cis-*eQTL but not the second. The lead variant in monocyte *cis-*eQTL analysis, rs10740051, is in high LD with the second conditional *cis-*eQTL, indicating that this *cis-*eQTL is active specifically in monocytes. Thus, the full GTEx whole blood *cis-* eQTL pattern and allelic heterogeneity is composed of *cis-*eQTLs that are active in different cell types. **(E)** COLOC posterior probability (PP4) of GWAS colocalization with whole blood ieQTLs and eQTLs of the same eGene for 36 GWAS traits.

Analysis of functional enrichment of neutrophil ieQTLs and isQTLs shows that these largely follow the enrichment patterns observed for bulk tissue *cis*-QTLs (fig. 7b), with ieQTLs more strongly enriched in promoter flanking regions and enhancers, which are known to be major drivers of cell type specific regulatory effects (*2*). We observed similar patterns for epithelial cell ieQTLs (fig. S44).

We hypothesized that the widespread allelic heterogeneity observed in the bulk tissue *cis*-eQTL data is partially driven by an aggregate signal from *cis*-eQTLs that are each active in a different cell type present in the tissue. Indeed, the number of *cis*-eQTLs per gene is higher for ieGenes than for standard eGenes in several tissues (fig. 7c). While differences in power could contribute to this pattern, it is strongly corroborated by eGenes that have independent *cis*-eQTLs (LD < 0.05) in five purified blood cell types (*58*) also showing an increased amount of allelic heterogeneity in GTEx whole blood (fig. 7c,d). Thus, insights into cell type specificity provides new understanding of mechanisms of genetic architecture of gene expression, with promise of improved resolution into complex patterns of allelic heterogeneity when effects manifesting in different cell types can be distinguished from each other.

Next, we analyzed how cell type interacting *cis*-QTLs contribute to the interpretation of regulatory variants underlying complex disease risk. GWAS colocalization analysis of neutrophil ieQTLs (*11*) revealed multiple loci (111, ∼32%) that colocalize only with ieQTLs and not with whole blood *cis*-eQTLs (fig. 7e), even though 75% (42/56) of the corresponding eGenes have both *cis*-eQTLs and ieQTLs. Improved resolution into allelic heterogeneity appears to contribute to this, with fig. S45 showing an example of a locus where the absence of colocalization between a platelet count GWAS signal and bulk tissue *cis*-eQTL for *SPAG7* appears to be due to the whole blood signal being an aggregate of multiple independent signals. The neutrophil ieQTL analysis uncovers a specific signal that mirrors the GWAS association, suggesting that platelet counts are affected by *SPAG7* expression only in specific cell type(s). Thus, in addition to novel colocalizations pinpointing potential causal genes, ieQTL analysis has the potential to provide insights into cell type specific mechanisms of complex traits.

## Discussion

The GTEx v8 data release represents the deepest survey of both intra- and inter-individual transcriptome variation across a large number of tissues. With 838 donors and 15,253 samples, we have created a comprehensive catalog of genetic variants that influence gene expression and splicing in *cis*. The fine-mapping data of GTEx *cis*-eQTLs provides a catalog of thousands of likely causal functional variants – the largest resource of this type. While *trans*-QTL discovery, as well as characterization of sex-specific and population-specific genetic effects, are still limited by sample size, analyses of the V8 data provide important insights into each. Cell type interacting *cis*-eQTLs and *cis*-sQTLs, mapped using computational estimates of cell type enrichment, constitute an important addition to the GTEx resource. The strikingly similar tissue-sharing patterns across these data types suggests shared biology from cell type composition to transcriptome variation and genetic regulatory effects. Our results indicate that shared cell types between tissues may be a key factor behind tissue-sharing of genetic regulatory effects, which will constitute a key challenge to tackle in the future. Finally, GWAS colocalization with *cis*-eQTLs and *cis*-sQTLs provides rich opportunities for further functional follow-up and characterization of regulatory mechanisms of GWAS associations.

Given the very large number of *cis*-eQTLs, the extensive allelic heterogeneity – multiple independent regulatory variants affecting the same gene – is unsurprising. With well-powered *cis*-QTL mapping, it becomes possible and important to describe and disentangle these effects; the assumption of a single causal variant in a *cis*-eQTL locus no longer holds true for data sets of this scale. Similarly, we highlight *cis*-eQTL and *cis*-sQTL effects on the same gene, typically driven by distinct causal variants. The joint complex trait contribution of independent *cis*-eQTLs and *cis*-sQTLs, and *cis*-eQTLs and rare coding variants for the same gene highlights how different genetic variants and functional perturbations can converge at the gene level to similar physiological effects. This orthogonal evidence pinpoints gold-standard disease genes, and could be leveraged to build allelic series, a powerful tool for estimating dosage-risk relationship for the purposes of drug development (*59*). Finally, we provide mechanistic insights into the cellular causes of allelic heterogeneity, showing the separate contributions from *cis*-eQTLs active in different cell types to the combined signal seen in a bulk tissue sample. With evidence that this increased cellular resolution improves colocalization in some loci, cell type specific analyses appear particularly promising for finer dissection of genetic association data.

Integration of GTEx QTL data and functional annotation of the genome provides powerful insights into the molecular mechanisms of transcriptional and post-transcriptional regulation that affect gene expression levels and splicing. A large proportion of *cis*-eQTL effects are driven by genetic perturbations in classical regulatory elements of promoters and enhancers, with an enrichment of tissue-specific and cell-type interacting *cis*-eQTLs in enhancers and related elements that thus contribute to context-specific genetic effects. Furthermore, we demonstrate that regulatory elements and transcription factors with variable activity across tissues and cell types modify *cis*-QTL effect sizes. While *cis*-eQTLs are enriched for a wide range of functional regions, the vast majority of *cis*-sQTL are located in transcribed regions, with likely co-/post-transcriptional regulatory effects. Interestingly, these appear to be less tissue-specific, which likely contributes to the higher tissue-sharing of *cis*-sQTLs than *cis*-eQTLs.

Approximately half of the observed *trans*-eQTLs are mediated by *cis*-eQTLs, demonstrating how local genetic regulatory effects can translate to effects at the level of cellular pathways. All types of QTLs that were studied are strong mediators of genetic associations to complex traits, with a higher relative enrichment for *cis*-sQTLs than *cis*-eQTLs, with *trans*-eQTLs having the highest enrichment of all (*35*). With large GWAS/PheWAS studies having uncovered extensive pleiotropy of complex trait associations, the GTEx data provide important insights into its molecular underpinnings: variants that affect the expression of multiple genes and multiple tissues have a higher degree of complex trait pleiotropy, indicating that some of the pleiotropy arises at the proximal regulatory level. Dissecting this complexity, and pinpointing truly causal molecular effects that mediate specific phenotype associations will be a considerable challenge for the future.

This study of the GTEx v8 data set has provided essential insights into genetic regulatory architecture and functional mechanisms. The extensive catalog of QTLs and associated data sets of annotations, cell types enrichments, and GWAS summary statistics provides rich material that requires careful interpretation for insights into the biology of gene regulation and functional mechanisms of complex traits. We have demonstrated how QTL data can be used to inform on multiple layers of GWAS interpretation: mapping of likely causal variants, proximal regulatory mechanisms, target genes in *cis*, pathway effects in *trans*, in the context of multiple tissues and cell types. However, our understanding of genetic effects on cellular phenotypes is far from complete. We envision that further investigation into genetic regulatory effects in specific cell types, study of additional tissues and developmental time points not covered by GTEx, incorporation of a diverse set of molecular phenotypes, and continued investment in increasing sample sizes from diverse populations will continue to provide transformative scientific discoveries.

## Data availability

All GTEx protected data are available via dbGaP (accession phs000424.v8). Access to the raw sequence data is now provided through the AnVIL platform (https://gtexportal.org/home/protectedDataAccess). The GTEx V8 non-protected data are available on the GTEx Portal, with multiple data views and analysis results publicly available on the Portal (www.gtexportal.org), as well as in the UCSC and Ensembl browsers. All components of the single tissue cis-QTL pipeline are available at https://github.com/broadinstitute/gtex-pipeline, and analysis scripts are available at https://github.com/broadinstitute/gtex-v8. Residual GTEx biospecimens have been banked, and remain available as a resource for further studies (access can be requested on the GTEx Portal, https://www.gtexportal.org/home/samplesPage).

## Authors

### Lead Analysts*

François Aguet^1#^, Alvaro N Barbeira^2^, Rodrigo Bonazzola^2^, Andrew Brown^3,4^, Stephane E Castel^5,6^, Brian Jo^7,8^, Silva Kasela^5,6^, Sarah Kim-Hellmuth^5,6,9^, Yanyu Liang^2^, Meritxell Oliva^2,10^, Princy Parsana^11^

### Analysts*

Elise Flynn^5,6^, Laure Fresard^12^, Eric R Gamazon^13,14,15,16^, Andrew R Hamel^17,1^, Yuan He^18^, Farhad Hormozdiari^19,1^, Pejman Mohammadi^5,6,20,21^, Manuel Muñoz-Aguirre^22,23^, YoSon Park^24,25^, Ashis Saha^11^, Ayellet V Segrè^1,17^, Benjamin J Strober^18^, Xiaoquan Wen^26^, Valentin Wucher^22^

### Manuscript Working Group*

François Aguet^1^, Kristin G Ardlie^1^, Alvaro N Barbeira^2^, Alexis Battle^18,11^, Rodrigo Bonazzola^2^, Andrew Brown^3,4^, Christopher D Brown^24^, Stephane E Castel^5,6^, Nancy Cox^16^, Sayantan Das^26^, Emmanouil T Dermitzakis^3,27,28^, Barbara E Engelhardt^7,8^, Elise Flynn^5,6^, Laure Fresard^12^, Eric R Gamazon^13,14,15,16^, Diego Garrido-Martín^22^, Nicole R Gay^29^, Gad Getz^1,30^, Roderic Guigó^22,31^, Andrew R Hamel^17,1^, Robert E Handsaker^32,33,34^, Yuan He^18^, Paul J Hoffman^5^, Farhad Hormozdiari^19,1^, Hae Kyung Im^2^, Brian Jo^7,8^, Silva Kasela^5,6^, Seva Kashin^32,33,34^, Sarah Kim-Hellmuth^5,6,9^, Alan Kwong^26^, Tuuli Lappalainen^5,6^, Xiao Li^1^, Yanyu Liang^2^, Daniel G MacArthur^33,35^, Pejman Mohammadi^5,6,20,21^, Stephen B Montgomery^12,29^, Manuel Muñoz-Aguirre^22,23^, Meritxell Oliva^2,10^, YoSon Park^24,25^, Princy Parsana^11^, John M Rouhana^17,1^, Ashis Saha^11^, Ayellet V Segrè^1,17^, Matthew Stephens^36^, Barbara E Stranger^2,37^, Benjamin J Strober^18^, Ellen Todres^1^, Ana Viñuela^38,3,27,28^, Gao Wang^36^, Xiaoquan Wen^26^, Valentin Wucher^22^, Yuxin Zou^39^

### Analysis Team Leaders*

François Aguet^1^, Alexis Battle^18,11^, Andrew Brown^3,4^, Stephane E Castel^5,6^, Barbara E Engelhardt^7,8^, Farhad Hormozdiari^19,1^, Hae Kyung Im^2^, Sarah Kim-Hellmuth^5,6,9^, Meritxell Oliva^2,10^, Barbara E Stranger^2,37^, Xiaoquan Wen^26^

### Senior Leadership*

Kristin G Ardlie^1^, Alexis Battle^18,11^, Christopher D Brown^24^, Nancy Cox^16^, Emmanouil T Dermitzakis^3,27,28^, Barbara E Engelhardt^7,8^, Gad Getz^1,30^, Roderic Guigó^22,31^, Hae Kyung Im^2^, Tuuli Lappalainen^5,6^, Stephen B Montgomery^12,29^, Barbara E Stranger^2,37^

### Manuscript Writing Group

François Aguet^1^, Hae Kyung Im^2^, Alexis Battle^18,11^, Kristin G Ardlie^1^, Tuuli Lappalainen^5,6^

### Corresponding Authors

François Aguet^1^, Kristin G Ardlie^1^, Tuuli Lappalainen^5,6^

## Acknowledgements

We thank the donors and their families for their generous gifts of organ donation for transplantation, and tissue donations for the GTEx research project; the Genomics Platform at the Broad Institute for data generation; Jeffrey Struewing for his support and leadership of the GTEx project; Mariya Khan and Christopher Stolte for the illustrations in Figure 1; and Ron Do, Daniel Jordan, and Marie Verbanck for providing GWAS pleiotropy scores. This work was funded by GTEx program grants: HHSN268201000029C (F.A., K.G.A., A.V.S., X.Li., E.T., S.G., A.G., S.A., K.H.H., D.Y.N., K.H., S.R.M., J.L.N.), 5U41HG009494 (F.A., K.G.A.), 10XS170 (Subcontract to Leidos Biomedical) (W.F.L., J.A.T., G.K., A.M., S.S., R.H., G.Wa., M.J., M.Wa., L.E.B., C.J., J.W., B.R., M.Hu., K.M., L.A.S., H.M.G., M.Mo., L.K.B.), 10XS171 (Subcontract to Leidos Biomedical) (B.A.F., M.T.M., E.K., B.M.G., K.D.R., J.B.), 10ST1035 (Subcontract to Leidos Biomedical) (S.D.J., D.C.R., D.R.V.), R01DA006227-17 (D.C.M., D.A.D.), Supplement to University of Miami grant DA006227. (D.C.M., D.A.D.), HHSN261200800001E (A.M.S., D.E.T., N.V.R., J.A.M., L.S., M.E.B., L.Q., T.K., D.B., K.R., A.U.), R01MH101814 (M.M-A., V.W., S.B.M., R.G., E.T.D., D.G-M., A.V.), U01HG007593 (S.B.M.), R01MH101822 (C.D.B.), U01HG007598 (M.O., B.E.S.), U01MH104393 (A.P.F.), as well as other funding sources: R01MH106842 (T.L., P.M., E.F., P.J.H.), R01HL142028 (T.L., Si.Ka., P.J.H.), R01GM122924 (T.L., S.E.C.), R01MH107666 (H.K.I.), P30DK020595 (H.K.I.), UM1HG008901 (T.L.), R01GM124486 (T.L.), R01HG010067 (Y.Pa.), R01HG002585 (G.Wa., M.St.), Gordon and Betty Moore Foundation GBMF 4559 (G.Wa., M.St.), 1K99HG009916-01 (S.E.C.), R01HG006855 (Se.Ka., R.E.H.), BIO2015-70777-P, Ministerio de Economia y Competitividad and FEDER funds (M.M-A., V.W., R.G., D.G-M.), NIH CTSA grant UL1TR002550-01 (P.M.), Marie-Skłodowska Curie fellowship H2020 Grant 706636 (S.K-H.), R35HG010718 (E.R.G.), FPU15/03635, Ministerio de Educación, Cultura y Deporte (M.M-A.), R01MH109905, 1R01HG010480 (A.Ba.), Searle Scholar Program (A.Ba.), R01HG008150 (S.B.M.), 5T32HG000044-22, NHGRI Institutional Training Grant in Genome Science (N.R.G.), EU IMI program (UE7-DIRECT-115317-1) (E.T.D., A.V.), FNS funded project RNA1 (31003A_149984) (E.T.D., A.V.), DK110919 (F.H.), F32HG009987 (F.H.)

