## Supplementary material for "The GTEx Consortium atlas of genetic regulatory effects across human tissues"

### Supplementary Materials

The GTEx Consortium

October 2019

**7 Sex-biased *cis*-eQTL mapping** **21**

**8 Population-biased *cis*-eQTL mapping** **22**

**9 Genomic annotation data** **24**

**10 Fine mapping of *cis*-eQTLs** **25**

**15 Supplementary Table legends** **59**

**16 Author Contributions** **60**

### 1 Biospecimen Collection and Processing

#### 1.1 Donor characteristics

All human donors were deceased, with informed consent obtained via next-of-kin consent for the collection and banking of de-identified tissue samples for scientific research [1].

Both sexes were enrolled as GTEx donors (**fig. S1**) but males represent approximately two thirds of the final cohort. All eligible donor age groups (20-70 years) are represented, but most enrolled donors were older individuals. While most of the donors are white, the V8 release includes 103 black or African American individuals. Diverse causes of death are represented, with common causes of mortality in the general population being typically common also among GTEx donors. The GTEx eligibility requirements are described in [1] and excluded individuals with metastatic cancer and individuals who had received chemotherapy for cancer within the prior two years. From each donor, a median of 19 tissues had RNA-seq data available after quality control.

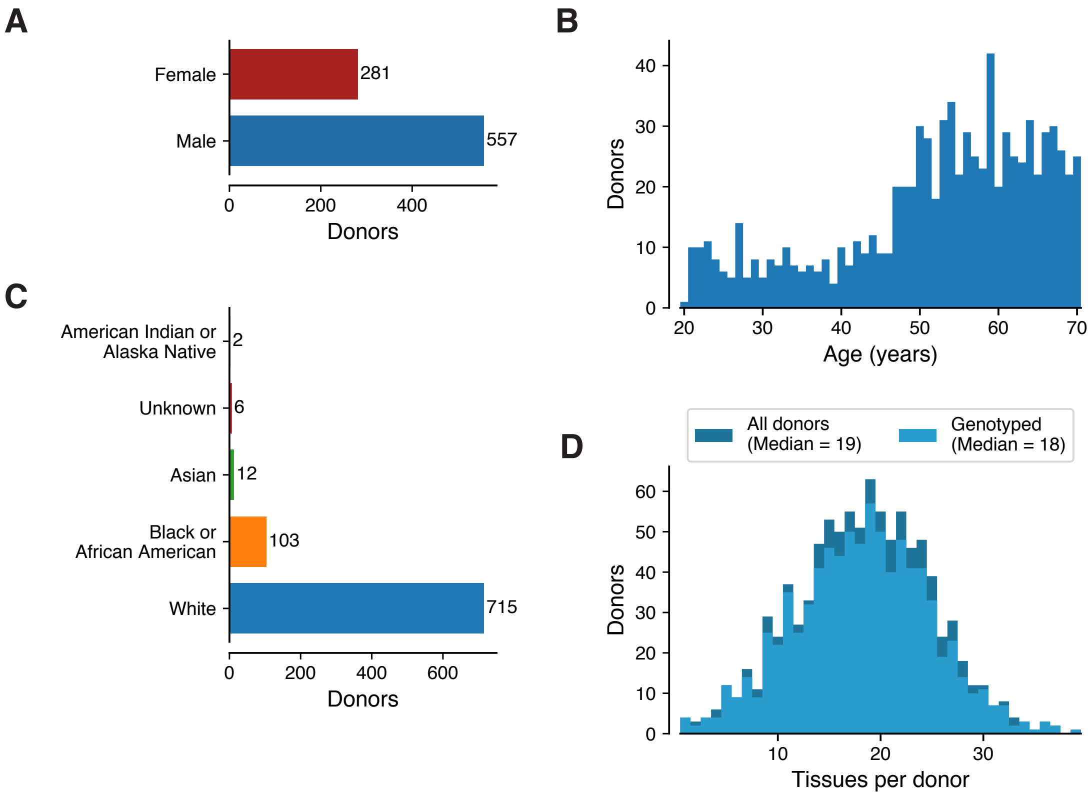

**Fig. S1. Donor characteristics**. Distribution of A) sex, B) age, C) self-identified ancestry, and D) the number of collected tissues.

#### 1.2 Biospecimen collection

The biospecimen collection is described in detail in [2], and a complete description of the donor enrollment and consent process, as well as biospecimen procurement methods, sample fixation, and histopathological review procedures are described in [1]. In brief, whole blood and skin samples were collected from each donor, and shipped overnight to the GTEx Laboratory Data Analysis and Coordination Center (LDACC) at the Broad Institute. These samples were used for DNA genotyping (primarily from whole blood), RNA expression analysis, and culturing and transformation of fibroblast and lymphoblastoid cell lines, respectively. In addition to these samples, two adjacent aliquots were prepared from all other sampled tissues and preserved in PAXgene tissue kits, with ischemic time varying between the different tissue sites (**fig. S2**). One of each paired samples was embedded in paraffin (PFPE) for histopathological review and the second was shipped to the LDACC for processing and molecular analysis. Brains were collected from approximately one-third of the donors, and were shipped on ice to the brain bank at the University of Miami, where eleven brain sub-regions were sampled and flash-frozen. These samples were then shipped to the LDACC for processing and analysis.

A robust quality management program was established and implemented for data management, Standard Operating Procedure (SOP) development, and auditing of collections. Document control software was used to ensure all biospecimen collection sites used current versions of SOPs, and training was conducted prior to implementation of all new procedures. Supporting quality documents were developed to provide consistency and clarity to the program, and many of those documents, such as the SOPs used and workflows for the project, are available to the public (http://biospecimens.cancer.gov/resources/sops/default.asp).

#### 1.3 Molecular analyte extraction and QC

Detailed protocols for the extraction of DNA and RNA from blood, cell pellets, and PAXgene-fixed and frozen tissues were described in [2]. The same protocols were used for all GTEx samples to avoid introduction of batch effects among samples, which were processed continually throughout the project. To control for variable RNA quality [2], RNA sequencing was only performed for samples with a RIN score of 5.5 or higher and with at least 500 ng of total RNA.

The 49 tissues with $\geq$70 genotyped samples that were included in the QTL and other downstream analyses vary in their sample size (n=73 to 706), ischemic time, and RNA quality (RIN). Additionally, the donor age range varies by tissue; notably the brain samples were collected primarily from older individuals (**fig. S2**).

| Tissue Site Detail | Abbreviation | Color | Samples from genotyped donors |
|---|---|---|---|
| Adipose - Subcutaneous | ADPSBQ | #FF6600 | 581 |
| Adipose - Visceral (Omentum) | ADPVSC | #FFAA00 | 469 |
| Adrenal Gland | ADRNLG | #33DD33 | 233 |
| Artery - Aorta | ARTAORT | #FF5555 | 387 |
| Artery - Coronary | ARTCRN | #FFAA99 | 213 |
| Artery - Tibial | ARTTBL | #FF0000 | 584 |
| Brain - Amygdala | BRNAMY | #EEEE00 | 129 |
| Brain - Anterior cingulate cortex (BA24) | BRNACC | #EEEE00 | 147 |
| Brain - Caudate (basal ganglia) | BRNCDT | #EEEE00 | 194 |
| Brain - Cerebellar Hemisphere [Frozen] | BRNCHB | #EEEE00 | 175 |
| Brain - Cerebellum [PAXgene] | BRNCHA | #EEEE00 | 209 |
| Brain - Cortex [PAXgene] | BRNCTXA | #EEEE00 | 205 |
| Brain - Frontal Cortex (BA9) [Frozen] | BRNCTXB | #EEEE00 | 175 |
| Brain - Hippocampus | BRNHPP | #EEEE00 | 165 |
| Brain - Hypothalamus | BRNHPT | #EEEE00 | 170 |
| Brain - Nucleus accumbens (basal ganglia) | BRNNCC | #EEEE00 | 202 |
| Brain - Putamen (basal ganglia) | BRNPTM | #EEEE00 | 170 |
| Brain - Spinal cord (cervical c-1) | BRNSPC | #EEEE00 | 126 |
| Brain - Substantia nigra | BRNSNG | #EEEE00 | 114 |
| Breast - Mammary Tissue | BREAST | #33CCCC | 396 |
| Cells - EBV-transformed lymphocytes | LCL | #CC66FF | 147 |
| Cells - Cultured fibroblasts | FIBRBLS | #AAEEFF | 483 |
| Colon - Sigmoid | CLNSGM | #EEBB77 | 318 |
| Colon - Transverse | CLNTRN | #CC9955 | 368 |
| Esophagus - Gastroesophageal Junction | ESPGEJ | #8B7355 | 330 |
| Esophagus - Mucosa | ESPMCS | #552200 | 497 |
| Esophagus - Muscularis | ESPMSL | #BB9988 | 465 |
| Heart - Atrial Appendage | HRTAA | #9900FF | 372 |
| Heart - Left Ventricle | HRTLV | #660099 | 386 |
| Kidney - Cortex | KDNCTX | #22FFDD | 73 |
| Liver | LIVER | #AABB66 | 208 |
| Lung | LUNG | #99FF00 | 515 |
| Minor Salivary Gland | SLVRYG | #99BB88 | 144 |
| Muscle - Skeletal | MSCLSK | #AAAAFF | 706 |
| Nerve - Tibial | NERVET | #FFD700 | 532 |
| Ovary | OVARY | #FFAAFF | 167 |
| Pancreas | PNCREAS | #995522 | 305 |
| Pituitary | PTTARY | #AAFF99 | 237 |
| Prostate | PRSTTE | #DDDDDD | 221 |
| Skin - Not Sun Exposed (Suprapubic) | SKINNS | #0000FF | 517 |
| Skin - Sun Exposed (Lower leg) | SKINS | #7777FF | 605 |
| Small Intestine - Terminal Ileum | SNTTRM | #555522 | 174 |
| Spleen | SPLEEN | #778855 | 227 |
| Stomach | STMACH | #FFDD99 | 324 |
| Testis | TESTIS | #AAAAAA | 322 |
| Thyroid | THYROID | #006600 | 574 |
| Uterus | UTERUS | #FF66FF | 129 |
| Vagina | VAGINA | #FF5599 | 141 |
| Whole Blood | WHLBLD | #FF00BB | 670 |

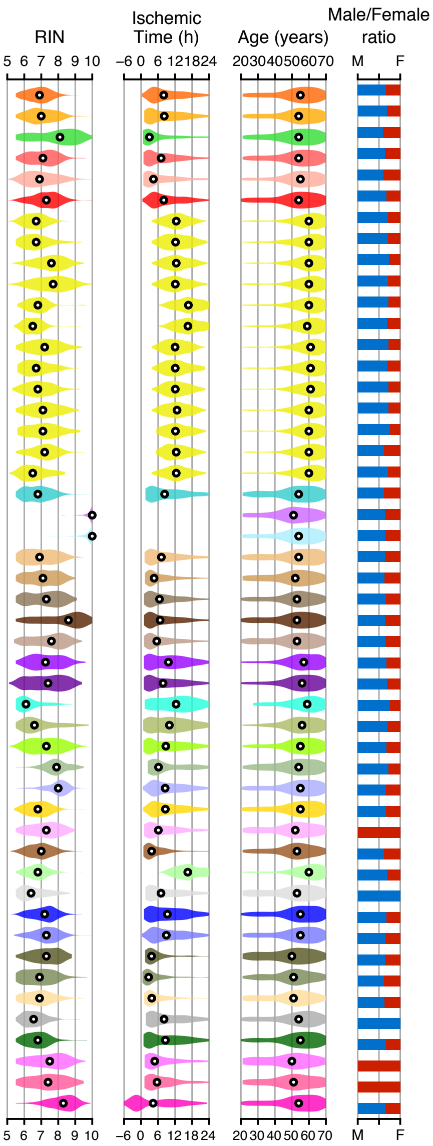

**Fig. S2. Summary of the tissues and samples of the QTL analysis**. Frontal Cortex and Cerebellar Hemisphere were sampled in duplicate: each was sampled on site during initial tissue collection (BRNCHA and BRNCTXA), and again after the brain was received by the brain bank (BRNCHB and BRNCTXB). Two cell lines were generated: an EBV-transformed lymphoblastoid cell line from blood (LCL) and cultured primary fibroblasts from fresh skin (FIBRBLS). RIN (RNA integrity number), ischemic time, and donor age distributions for each tissue are shown as density plots, with the median indicated in black; donor sex distributions are shown as stacked bar plots.

### 2 Whole Genome Sequencing

#### 2.1 Whole genome sequencing

Whole genome sequencing (WGS) was performed for 899 samples from 869 unique GTEx donors, to a median depth of 32×. Sequencing methods and protocols were improved and updated several times over the course of the GTEx project, and hence samples were occasionally resequenced using newer protocols to enable comparisons with older-sequenced samples, resulting in sample duplicates. Samples and general protocols are as follows: Libraries of whole genome DNA from 79 GTEx donors were sequenced on an Illumina HiSeq 2000 at the Broad institute, using a PCR-based protocol, and as 101-bp paired-end reads for 67 samples and 151-bp paired-end reads for 12 samples. Libraries of whole genome DNA from 820 samples from 801 donors (including 11 that were also sequenced on HiSeq 2000) were sequenced on an Illumina HiSeq X at the Broad Institute as 151-bp paired-end reads. 571 of the samples were sequenced using a PCR-based protocol, and the remaining 249 using a PCR-free protocol (17 samples were sequenced using both protocols, and two samples were sequenced in duplicate using the PCR-free protocol). Library construction was performed as described in [3], with minor modifications including replacing the Illumina paired-end adapters with palindromic forked adapters with unique 8-base index sequences embedded within the adapter. Sequencing was performed following the manufacturer's protocol. All sample information tracking was performed by automated LIMS messaging.

Of the 899 samples, 30 were lower-quality replicates, and samples from 31 donors were excluded from further analyses for the following reasons: large chromosomal abnormalities were observed for 22, including two with chr21 trisomy confirmed to have Down's syndrome, one with a chr17p mosaic trisomy [4], four Klinefelter individuals (three confirmed by histological examination of testis tissue and one identified based only on gene expression where no testis tissue was available), one XXX female, and 14 had large (>1Mb) duplication and/or deletion events; three donors had documented sepsis; one had cerebral palsy; one sex-mismatch who was genetically XY based on exome and Xist expression analysis but underwent sex reassignment surgery to a female shortly after birth; one was related to another donor; and samples from 3 donors were aligned with a different pipeline at the time variant calling was performed and therefore excluded from further analyses. The final analysis freeze set contained variant calls from 838 donors. DNA for WGS was derived primarily from whole blood (779/838 samples). However, for some donors, a blood sample was not collected, or the DNA extracted from the whole blood was poor in quality, in which case a tissue sample was used as a DNA source (thyroid tissue was used for 14 donors and lung for 11, with the remainder scattered across tissue types).

The quality control steps that led to the identification of flagged samples and donors, as well as the variant calling and quality control pipeline are described in the sections below.

#### 2.2 WGS data processing

Output from Illumina software was processed with a pipeline based on Picard (http://broadinstitute.github.io/picard/) and reads were aligned with BWA-MEM (http://bio-bwa.sourceforge.net), using base quality score recalibration and local realignment at known insertions and deletions (indels), to yield BAM files aligned to the human reference genome build GRCh38 (including all ALT, HLA, and decoy sequences). Reference files used to generate the variant call set, including the human reference genome, the whole genome calling interval list, known indels used for local realignment, and known variants for Variant Quality Score Recalibration (VQSR) are available at https://console.cloud.google.com/storage/browser/genomics-public-data/resources/broad/hg38/v0/.

#### 2.3 Variant calling and quality control

Variants (SNPs and indels) were jointly called across an initial, pre-quality control set of 927 GTEx WGS samples combined with 6 non-GTEx WGS samples for quality control purposes, using GATK HaplotypeCaller v3.5. Only autosomes and chromosome X were used in variant calling. The non-GTEx samples were subsequently removed from the VCF, as well as 61 GTEx samples that failed sample QC (based on BAM- and VCF-derived QC metrics), yielding a total of 866 donors with WGS variant calls. Multi-allelic sites were split into biallelic sites using Hail v0.1 (http://hail.is). Compound HET calls (calls with two different ALT alleles, e.g., ALT1/ALT2) were encoded as 0/1, with '1' referring to the ALT allele of the split biallelic site. All VCFs and tables described in the sections below are available as a part of the dbGaP release.

##### 2.3.1 WGS sample-level quality control

BAM- and VCF-based statistics were computed to detect technical outliers among the WGS BAMs: (i) BAM-level summary statistics and outlier cutoffs included mean sequence coverage (<25×), percent of chimeric reads (>0.05), and median and standard deviation of insert size, computed with Picard, and contamination rate (>0.05) estimated with VerifyBamID (https://genome.sph.umich.edu/wiki/VerifyBamID). Samples were also tested for RNA contamination by assessing levels of split reads aligning to exon-exon junctions in the transcriptome using HISAT [5], and for bacterial contamination based on the fraction (>0.05) of read pairs with short insert sizes (<30bp); (ii) VCF-based sample-level QC metrics used for outlier detection and computed with GATK included: call rate, number of SNPs, number of deletions, number of insertions, insertion to deletion ratio, transition to transversion ratio, heterozygous to homozygous ratio. All samples had a call rate above 99%. Samples that were 4 median absolute deviations above or below the median of any of the above QC metrics were manually inspected. Extreme outliers with 4 median absolute deviations from the median for several QC metrics were excluded from further analyses. Samples were evaluated within ancestry, sequencing technology, and PCR+/- sample subsets.

Principal component analysis (PCA) was performed pre-QC using Hail v0.1 (0.1_ff26e57, https://hail.is) and a pruned set of SNPs ($r^2 < 0.01$) to determine the samples' genetic ancestry. This was needed for both proper outlier evaluation of the VCF-based QC metrics within each subset of samples, and to check the self-reported ancestry. The ancestry of donors was inferred using the first 3 genotype PCs and K-nearest neighbors (k = 3), using the QCed ancestry of WGS donors from release v7 as the training set. The inferred ancestry was checked against self-reported ones. In three samples, the self-reported and inferred ancestry were remarkably different, with samples positioning in the middle of a different cluster, and for these three samples the subjects attributes file was updated to the inferred ancestry.

Genetic relatedness was evaluated by computing identity by descent (IBD) in Hail, and one sample among a pair of individuals with IBD of ∼0.25 (corresponding to first degree cousins) was excluded from downstream analyses.

###### 2.3.2 Long copy number variation analyses

To identify samples with large chromosomal abnormalities (both to detect samples with quality issues and biological outliers for exclusion from the analysis freeze), we ran the Genome STRiP [6] module that detects Long Copy Number Variation (>1Mb) on the 899 WGS BAMs that passed sample QC. 37 samples were found to have at least one duplication or deletion of >1Mb (list provided to dbGaP). Of these samples, 14 were flagged for including potentially pathogenic large CNVs based on literature support (reasons listed in `GTEx_Analysis_2017-06-05_v8_WholeGenomeSeq_flagged_donors.xlsx` in the WGS VCF archive on dbGaP), as well as full chromosomal duplications, which included two individuals with Down Syndrome and an individual with chr17 trisomy. An additional 3 WGS samples were flagged for large CNVs detected by Genome STRiP in release V7. We also used the GENOME STRiP ChrX and ChrY dosage estimates for to check for sex discrepancies. In addition to the detection of the 3 Klinefelter individuals and the XXX female, we detected one XY self-identified female who underwent sex reassignment surgery to a female shortly after birth.

###### 2.3.3 Sample set of the WGS analysis freeze

In addition to the 22 biological outliers detected with Genome STRiP, 4 samples with outlier clinical phenotypes (sepsis or cerebral palsy), an additional Klinefelter individual detected only by RNA-seq sex check, and 1 sample from a pair of related donors (first degree cousins, IBD of ∼0.25) were excluded from downstream analyses, yielding a total of 838 donors for the WGS analysis freeze in the GTEx V8. A summary of all excluded samples is available on dbGaP. Outlier samples were removed from the VCF using SelectVariants from GATK v3.7-0-gcfedb67.

###### 2.3.4 WGS variant quality control

To increase the quality of genotype calls in the VCF with 838 analysis freeze samples, genotype quality scores (GQ) and genotype posteriors (PP) were recomputed with genotype likelihoods and allele frequencies and counts from 1000 Genomes Project Phase 3 (lifted over from GRCh37 to GRCh38) as a reference panel (ftp://ftp.1000genomes.ebi.ac.uk/vol1/ftp/release/20130502/supporting/GRCh38_positions/), using the CalculateGenotypePosteriors module in GATK v3.5. Multi-allelic sites were split into biallelic sites using Hail. Compound HET calls (e.g., ALT1/ALT2) were set to missing. To obtain high confidence variant calls for downstream analyses, extensive variant QC was applied to all variant sites (see **Table S1**). A variant site was removed based on any the following criteria: (i) failed VQSR at a sensitivity level below 99.8% for SNPs or 99.95% for InDels; (ii) lay in a Low Complexity Region (LCR); (iii) had a low Inbreeding Coefficient (<-0.3); (iv) became monomorphic after assigning the following genotype calls to missing: compound HET sites (ALT1/ALT2) after splitting multi-allelic sites to bi-allelic sites, low genotype quality (GQ<20), calls with allelic imbalance (AB>0.8 or AB<0.2), or heterozygous calls in chrX nonPAR regions in male samples; (v) had a missingness rate >= 15%; (vi) failed Hardy-Weinberg Equilibrium testing ($P<10^{-8}$) in European or African American subsets for autosomes or in European females for the X chromosome; (vii) showed significant association ($P<10^{-8}$) with the sequencing technology, library construction batch, or PCR+ versus PCR- library preparation; or (viii) showed significant non-random missingness of reference alleles with MAF>1% on the autosomes. All variant QC metrics were computed at the site level except for GQ and AB, and compound HET filters that were at the genotype call level. The number of variants removed sequentially at each QC step is summarized in **Table S1**. All QC-failed variants were removed from the VCF using Hail, yielding a post-sample and variant QCed WGS VCF for release v8. All computed QC metric statistics were added back to the post-sample, pre-variant QC'd VCF with 838 individuals, to enable custom QC for other projects. In this VCF, the 0/1 encoding of compound HET calls in split biallelic sites was kept, so that multi-allelic sites can be reconstructed from all their biallelic sites. Variant QC was performed using GATK v3.5, Hail v0.1 (0.1_ff26e57), and PLINK 1.9 (https://www.cog-genomics.org/plink2). All sample- and variant-level QC steps were performed using a custom pipeline developed in R, Python, and Hail v0.1.

| **Filtering criterion** | **Total sites** | **Bi-allelic sites split from InDels** | **Multi-allelic sites** | **MAF $\geq$ 1%** |
|---|---|---|---|---|
| Initial variant calls | 66,463,168 | 54,152,863 | 11,279,240 | 17,900,327 |
| VQSR not PASS* | 58,492,632 | 46,391,982 | 9,637,709 | 16,340,678 |
| In LCRs | 49,108,668 | 44,438,858 | 2,279,757 | 11,692,570 |
| InbreedingCoeff $<$ -0.3 | 48,821,365 | 44,176,119 | 2,229,674 | 11,436,058 |
| Monomorphic after assigning compound HET sites (ALT1/ALT2) to missing after splitting multi-allelic sites to bi-allelic sites | 48,772,911 | 44,149,214 | 2,181,220 | 11,457,236 |
| Monomorphic after assigning genotype calls with AB $>$ 0.8 or AB $<$ 0.2 to missing | 48,138,994 | 43,792,153 | 2,067,813 | 11,290,157 |
| Monomorphic after assigning genotype calls with GQ $<$ 20 (GQ calculated from PPs) to missing** | 47,911,767 | 43,609,447 | 2,043,813 | 11,382,683 |
| Monomorphic after assigning heterozygous genotype calls on chrX nonPAR males to missing | 47,887,958 | 43,597,559 | 2,039,377 | 11,379,387 |
| Missingness $\geq$ 15% | 46,629,217 | 43,114,532 | 1,225,732 | 10,784,573 |
| HWE test ($P < 10^{-8}$) | 46,610,074 | 43,098,576 | 1,223,708 | 10,765,916 |
| PCR+/- batch association ($P < 10^{-8}$) | 46,609,670 | 43,098,368 | 1,223,564 | 10,765,524 |
| HiSeq batch association ($P < 10^{-8}$) | 46,608,349 | 43,097,627 | 1,223,180 | 10,764,630 |
| LCSET batch association ($P < 10^{-8}$) | 46,607,502 | 43,096,968 | 1,223,039 | 10,763,912 |
| Non-random missingness for autosomal sites with haplotype MAF $>$ 1% ($P < 10^{-8}$)*** | 46,569,704 | 43,066,451 | 1,223,039 | 10,726,114 |

**Table S1. Genetic variant QC filtering**. Summary of sites removed after each variant QC step.
*VQSR cutoffs were set to 99.8% for SNPs and 99.95% for InDels.
**Genotype quality scores (GQ) were recomputed from genotype posterior probabilities (PPs) estimated with GATK's CalculateGenotypePosteriors, using 1000 Genomes Project Phase 3 as the reference panel.
*** This QC step was performed using PLINK 1.9. All other steps were implemented in Hail v0.1 (0.1_ff26e57).
All variant QC steps were performed sequentially, in the order listed, and variants that failed any criterion were not tested in subsequent steps.

##### 2.3.5 Genotype principal component analysis

We computed genotype principal components (PCs) based on the post-sample and variant QC WGS VCF, using EIGENSTRAT (`smartpca.perl -i <geno> -a <snp> -b <ind> -k 20 -m 0 -o out -e out.eval -p out.plot -l out.log`; https://github.com/argriffing/eigensoft/blob/master/EIGENSTRAT/). PCA was performed on a set of LD-independent variants with a call rate $\geq$ 99% and MAF $\geq$ 0.05. LD pruning was performed using Plink (`plink --bfile input --indep-pairwise 200 100 0.1 --out`).

The 838 post-QC GTEx samples clustered into 3 main subpopulations: European, African, and Asian (**fig. S3A**). We used the top 5 genotype PCs to correct for population stratification, as the top 3 PCs proportionately captured the most variance among subpopulations (**fig. S3B**), and the top 5 PCs significantly correlated with 11 subpopulations inferred for all GTEx WGS samples, using 1000 Genomes Project samples and k-nearest neighbors clustering (**fig. S3C**, $P < 3 \times 10^{-8}$ from F test; adjusted $R^2 = 0.053 - 0.98$)

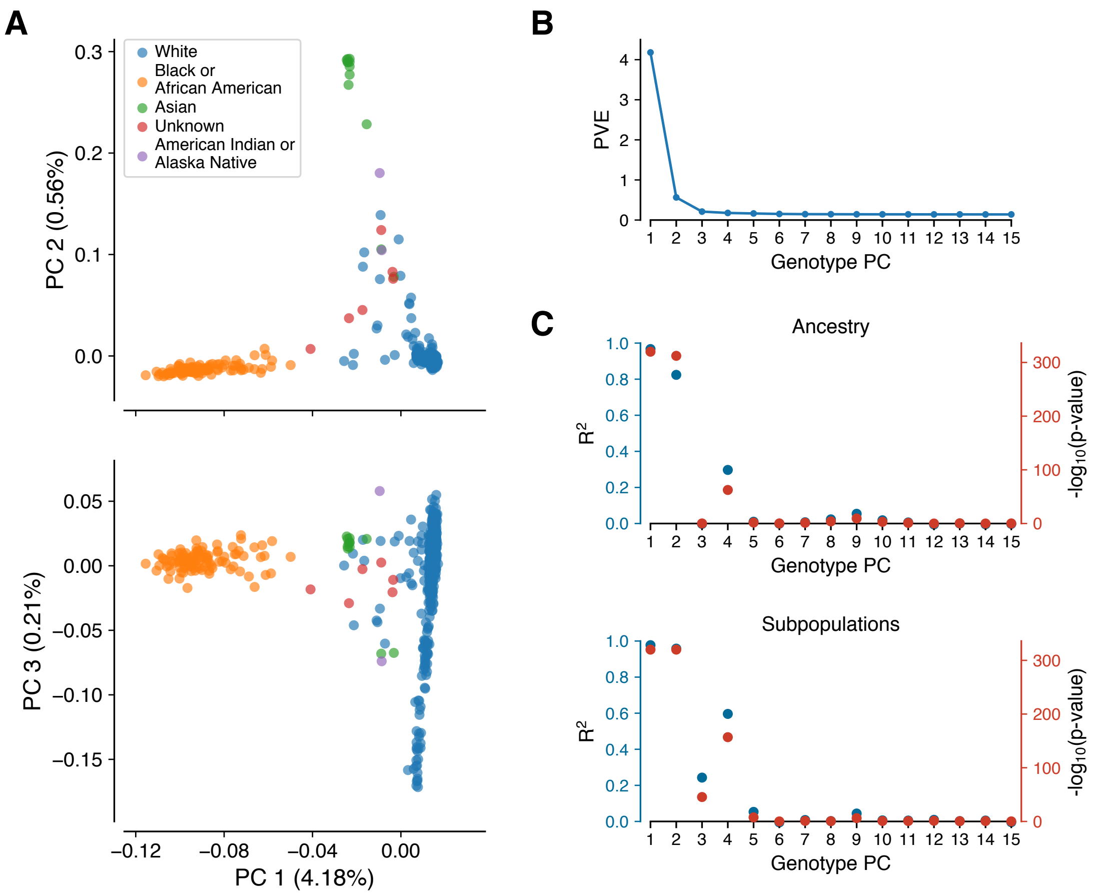

**Fig. S3. Genotype principal components**. (A) Top three genotype principal components (PCs), color coded by race as reported in the subject phenotype annotation file. (B) Percent variance explained (PVE) for the top 15 genotype PCs. (C) Correlation of genotype PCs with race as reported in the subject phenotype annotation file (top) and with 11 subpopulations inferred for all GTEx WGS samples, using 1000 Genomes Project samples and k-nearest neighbors clustering (bottom).

##### 2.3.6 Variant identifiers

Variant IDs were constructed by concatenating chr, position, REF and ALT alleles, and a suffix indicating the genome build (b38), separated by an underscore. For insertions and deletions, the preceding base is added. RS IDs from dbSNP 150 GRCh38p7 (`ftp://ftp.ncbi.nlm.nih.gov/snp/organisms/human_9606_b150_GRCh38p7/VCF`) were added to a variant lookup table available through the GTEx Portal.

#### 2.4 Read-aware phasing

To enable better functional interpretation of regulatory effects and to impute missing calls, read-aware phasing was performed on the sample- and variant-QCed VCF of 838 samples, using SHAPEIT v2 (r837) with extractPIRs (r68) [7]. The phasing procedure in SHAPEIT2 imputes missing calls and was performed in the following order:

1. extractPIRs was run to extract phase informative reads (PIRs) from the WGS BAMs from all individuals with the command: `extractPIRs --bam ${bam_list} --vcf ${vcf} --out ${pir}`. Defaults were used for filtering on minimal read mapping quality ($\geq$10) and base quality ($\geq$13).

2. SHAPEIT 2 was run with the command:
`shapeit -assemble --input-vcf ${vcf} --input-pir ${pir} -O ${haplotypes}`.
For chromosome X, PAR regions and non-PAR regions were phased separately, with PAR regions phased like autosomes and the non-PAR region phased using the `--chrX` and `--input-sex` flags.

3. The following post-processing step were applied: (i) calls with dosage differences before and after phasing were set to missing (1,185,212 sites had $\geq$1 sample set to missing); (ii) calls in split biallelic sites that were compound HETs (e.g.,

ALT1/ALT2) or alleles absent in that sample (e.g., a REF/ALT2 call for a REF/ALT1 bi-allelic site) that were coded as missing prior to phasing were reset to missing (43,380 sites had a different dosage after phasing in $\geq$1 sample); (iii) sites that became monomorphic after phasing or as a result of post-processing steps were removed; (iv) sites with missingness $\geq$15% after the above changes were removed.

The final phased VCF, which contains 46,526,292 sites, is the Analysis Freeze WGS VCF v8 used for downstream analyses. See also Section 5 for description of further RNA-seq read based phasing.

### 3 RNA Expression

#### 3.1 RNA library preparation and sequencing

RNA sequencing was performed at the Broad Institute using the Illumina TruSeq$^{TM}$ RNA sample preparation protocol, which was based on polyA+ selection of mRNA and was not strand-specific. This protocol was used continuously throughout the project to reduce the introduction of batch effects.

Briefly, total RNA was quantified using the Quant-iT$^{TM}$ RiboGreen®RNA Assay Kit and normalized to 5 ng per µL. An aliquot of 200 ng for each sample was transferred into library preparation, which was an automated variant of the Illumina TruSeq$^{TM}$ RNA sample preparation protocol (Revision A, 2010; (http://www.illumina.com/documents/products/datasheets/datasheet_truseq_sample_prep_kits.pdf). This method used oligo dT beads to select mRNA from the total RNA sample followed by heat fragmentation and cDNA synthesis from the RNA template. The resultant cDNA then went through library preparation (end repair, base 'A' addition, adapter ligation, and enrichment) using Broad Institute-designed indexed adapters substituted in for multiplexing. After enrichment, the libraries were quantified with qPCR using the KAPA Library Quantification Kit for Illumina Sequencing Platforms and then pooled equimolarly. The entire process was performed in 96-well plates and all pipetting was performed by either Agilent Bravo or Hamilton Starlet liquid handlers with electronic tracking throughout the process in real-time, including reagent lot numbers, specific automation used, time stamps for each process step, and automatic registration.

Pooled libraries were normalized to 2 nM and denatured using 0.1 N NaOH prior to sequencing. Flow cell cluster amplification and sequencing were performed according to the manufacturer's protocols using either the HiSeq 2000 or HiSeq 2500. Sequencing generated 76bp paired-end reads and an eight-base index barcode read and was run with a coverage goal of 50M reads (the median achieved was ~83M total reads). Raw sequence data were processed using the Broad Institute's Picard pipeline, which includes de-multiplexing and data aggregation steps.

#### 3.2 RNA-seq alignment and quality control

RNA-seq data were aligned to the human reference genome GRCh38/hg38 (excluding ALT, HLA, and decoy contigs) with STAR v2.5.3a [8]. To avoid potential artifacts from allelic mapping bias in allelic expression and sQTL analyses, the RNA-seq data were re-aligned with STAR v2.6.0c using the `--waspOutputMode` option [9] and a VCF containing the corresponding genotypes. These data were used in splicing quantification and allelic expression analysis.

Quality control of the samples was performed as described in [2]. Briefly, low-quality samples were identified and removed based on the following alignment metrics: $<$ 10 million mapped reads; read mapping rate $<$ 0.2; intergenic mapping rate $>$ 0.3; base mismatch rate (mismatched bases divided by total aligned bases) $>$ 0.01 for read mate 1 or $>$ 0.02 for read mate 2; rRNA mapping rate $>$ 0.3. Additionally, outlier samples were identified based on expression profile using a correlation-based statistic and sex incompatibility checks, following methods described in [10]. Among technical replicates (same aliquot sequenced multiple times for QC purposes), the sample with the highest number of reads was retained for inclusion in the analysis freeze set. Finally, samples from donors with cytogenetic anomalies (see Section 2) were excluded from analyses.

#### 3.3 Analysis freeze of tissues and samples for eQTL analyses

After QC, the v8 release contained 17,382 RNA-seq samples. Among these, samples were selected based on donor genotype availability and a threshold of at least 70 samples per tissue, resulting in a set of 15,201 samples from 49 tissues across 838 donors used for QTL analyses. The tissues and samples are summarized in **fig. S2**, which also contains the abbreviations and color scheme used throughout the paper.

#### 3.4 Quantification of gene expression and splicing

##### 3.4.1 Gene annotation

The quantification was based on the GENCODE Release 26 annotation (`https://www.gencodegenes.org/releases/26.html`), collapsed to a single transcript model for each gene, using a custom isoform collapsing procedure, comprising the following steps: 1) exons associated with transcripts annotated as "retained_intron" and "read_through" were excluded; 2) exon intervals overlapping within a gene were merged; 3) the intersections of exon intervals overlapping between genes were excluded; 4) the remaining exon intervals were mapped to their respective gene identifier and stored in GTF format. This annotation is available on the GTEx Portal (`gencode.v26.GRCh38.genes.gtf`).

All gene biotypes were used in QTL mapping in order to create a comprehensive *cis*-QTL data set for all quantified genes. However, since poly-A RNA-sequencing data is not expected to capture many noncoding gene types and their annotation and functional interpretation is often unclear, we used only protein-coding and lincRNA in downstream analyses. In *trans*-QTL mapping only protein-coding and lincRNA genes were used to avoid genes enriched for mapping artefacts.

##### 3.4.2 Gene expression quantification

Gene-level expression quantification was performed using RNA-SeQC [11]. Gene-level read counts and TPM values were produced using the following read-level filters: 1) reads were uniquely mapped (corresponding to a mapping quality of 255 for STAR BAMs); 2) reads were aligned in proper pairs; 3) the read alignment distance was $\leq 6$; 4) reads were fully contained within exon boundaries. Reads overlapping introns were not counted. These filters were applied using the "-strictMode" flag in RNA-SeQC.

Gene expression values for all samples from a given tissue were normalized for eQTL analyses using the following procedure: 1) read counts were normalized between samples using TMM [12]; 2) genes were selected based on expression thresholds of $\geq$0.1 TPM in $\geq$20% of samples and $\geq$6 reads (unnormalized) in $\geq$20% of samples; 3) expression values for each gene were inverse normal transformed across samples.

##### 3.4.3 Splicing quantification

We quantified splicing based on the intron excision phenotypes computed by LeafCutter [13] with the following steps and filters: 1) Intron usage was quantified using the `bam2junc.sh` script provided by the LeafCutter software, with an additional step to filter out reads that did not pass WASP filtering (specifically, reads with the tag `vW:i:[2-7]`; see Section 3.2); 2) Intron clusters were generated using the `leafcutter_cluster.py` script from LeafCutter, with the following options: `--min_clu_reads 30 --min_clu_ratio 0.001 --max_intron_len 500000` and mapped to genes using the `map_clusters_to_genes.R` script with exon coordinates derived from the collapsed gene model described in Section 3.4.1; 3) Introns with few counts or low complexity (diversity of counts across samples) were filtered out as follows to avoid numerical issues with the calculation of Beta-approximated empirical in splicing QTL p-values in FastQTL: introns without any read counts in >50% of samples, or with fewer than $\max(10, 0.1n)$ unique values, where $n$ is the sample size, were filtered out. Additionally, introns with insufficient variability across samples were removed based on thresholds applied to a z-score, $z$, of cluster read fractions across individuals (`*_perind.counts.gz` files): $(\sum_i(|z_i| < 0.25) \geq n-3) \wedge (\sum_i(|z| > 6) \leq 3)$. The latter step only removed a small number of outlier introns across tissues, with a maximum of 31 in Kidney - Cortex; 4) The filtered counts were normalized using the `prepare_phenotype_table.py` script from LeafCutter and the resulting per-chromosome files merged and converted to BED format with the start/end position corresponding to the TSS of the gene to which each intron was mapped in step 2.

#### 3.5 Latent factor analysis of expression and splicing variation

##### 3.5.1 PEER analysis of gene expression variation

To account for hidden batch effects and other technical and biological sources of transcriptome-wide variance in the gene expression data, we used the Probabilistic Estimation of Expression Residuals (PEER) method to estimate a set of latent covariates for gene expression levels for each tissue type [14]. The number of PEER factors was selected to maximize *cis*-eGene discovery, across four sample size bins: tissues with fewer than 150 samples, tissues with $\geq 150$ and $< 250$ samples, tissues with $\geq 250$ and $< 350$ samples, and tissues with $\geq 350$ samples. The optimization was performed as described in [15], and resulted in the selection of 15, 30, 45 and 60 PEER factors, respectively, for the four sample size bins.

The gene expression variance captured by PEER factors from each tissue were correlated with known technical and biological covariates recorded for each sample and donor (**fig. S4**). The covariates that were most consistently associated with PEER factors include factors related to parameters of donor death, ischemic time, sequencing quality control metrics, and nucleic acid isolation and library construction batches.

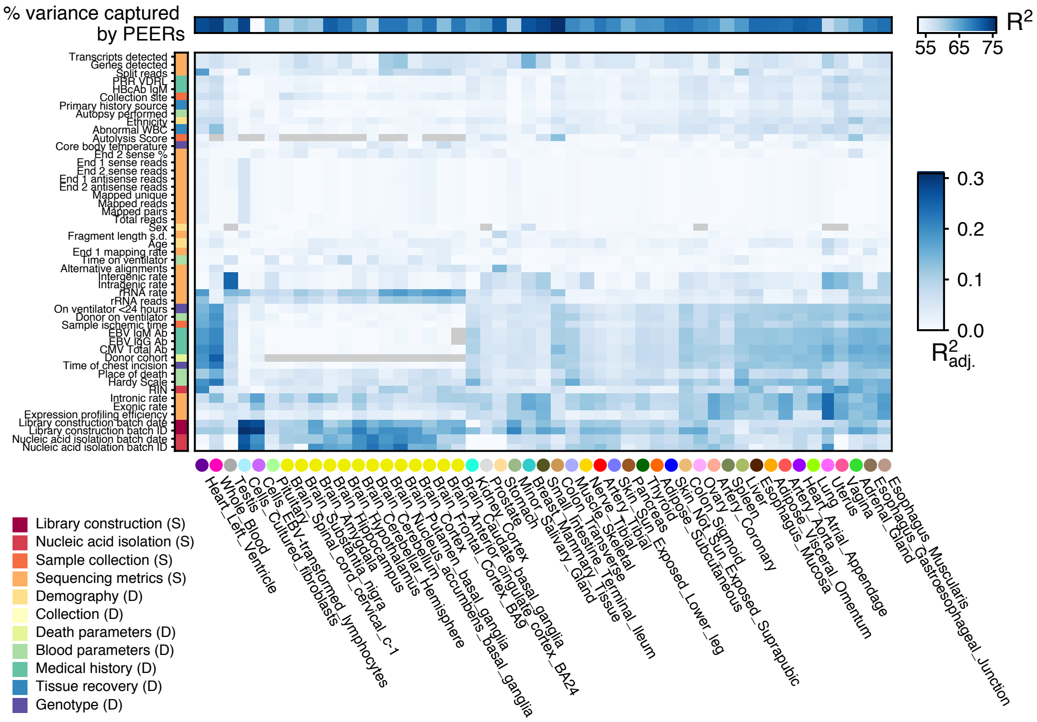

**Fig. S4. PEER factors for eQTL mapping**. Proportion of expression variance captured by the PEER factors computed for each tissue ($R^2$, top bar), and proportion of variance (adjusted $R^2$) removed by the PEER factors explained by known sample and donor covariates. Each cell shows the total proportion of variance removed by all PEER factors. Only covariates with $\geq$0.05 $R^2_{adj.}$ in any tissue are shown. Tissues and covariates are ordered based on hierarchical clustering with average Euclidean distance. Gray cells indicate unavailable data.

##### 3.5.2 PEER analysis of splicing variation

For splicing quantifications, PEER factors were computed based on the normalized counts matrices described in Section 3.4.3. Unlike *cis*-eQTL discovery, the number of *cis*-sGenes increased only marginally with increasing numbers of PEER factors, and varied less across tissues (**fig. S5**). As a result, 15 PEER factors were uniformly computed for each tissue.

The splicing variance captured by PEER factors most consistently correlated with nucleic acid extraction and library preparation batches, and also parameters of donor death and ischemic time (**fig. S5**).

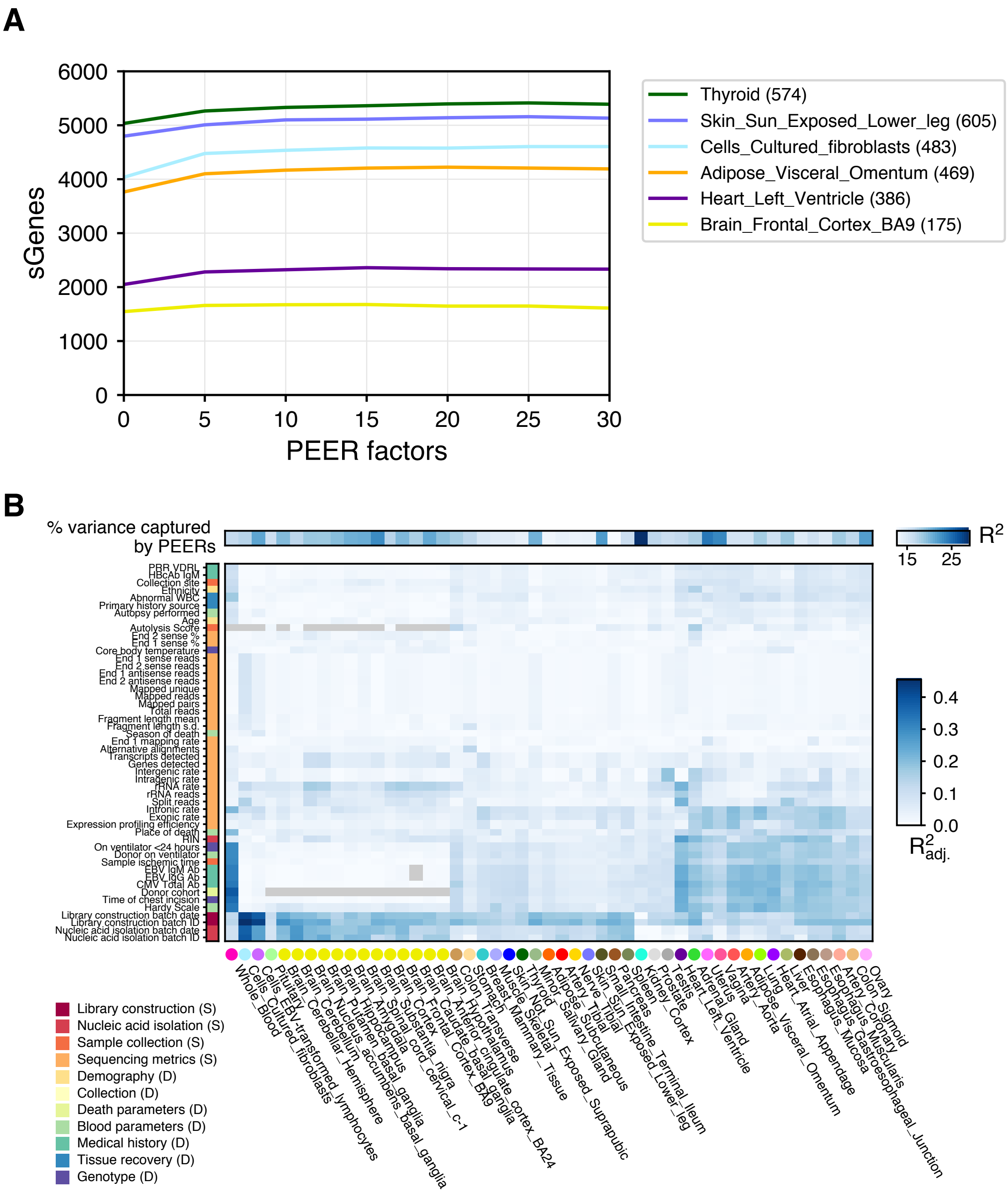

**Fig. S5. PEER factors for sQTL mapping**. A) *cis*-sGene discovery (0.05 FDR) as a function of PEER factors computed in increments of 5 factors. B) Proportion of intron inclusion variance captured by the 15 PEER factors computed for each tissue ($R^2$, top bar), and proportion of variance (adjusted $R^2$) removed by the PEER factors explained by known sample and donor covariates. Each cell shows the total proportion of variance removed by all PEER factors. Only covariates with $\geq$0.05 $R^2_{adj.}$ in any tissue are shown. Tissues and covariates are ordered based on hierarchical clustering with average Euclidean distance. Gray cells indicate unavailable data.

### 4 QTL mapping

#### 4.1 Covariates for QTL analysis

To control for population effects on the discovery of QTLs, we included the first five genotype PCs as covariates, which capture the major population structure among GTEx donors (**fig. S3**, and Section 2). Additionally, WGS sequencing platform (HiSeq 2000 or HiSeq X), WGS library construction protocol (PCR-based or PCR-free) and donor sex were included in the set of covariates used in the association analyses. We consider these to be the minimal set of covariates to use in most QTL mapping with GTEx data.

For eQTL mapping in *cis* and *trans* we used PEER factors optimized by sample size as described above in Section 3.5.1. For sQTL mapping in *cis* and *trans* we used 15 PEER factors as described above in Section 3.5.2.

#### 4.2 *cis*-eQTL mapping

*cis*-eQTL mapping was performed using FastQTL [16]. The mapping window was defined as 1 Mb up- and down-stream of the transcription start site (TSS), and the adaptive permutation mode was used with the setting `--permute 1000 10000`. The phased VCF described in Section 2.4 was used, and all variants with minor allele frequency $\geq$ 0.01 across the 838 donors were included. The same set of variants was tested in all tissues. The beta distribution-extrapolated empirical *P*-values from FastQTL were used to calculate gene-level q-values [17] with a fixed *P*-value interval for the estimation of $\pi_0$ (the 'lambda' parameter was set to 0.85). A false discovery rate (FDR) threshold of $\leq$ 0.05 was applied to identify genes with at least one significant *cis*-eQTL ("eGenes").

To identify the list of all significant variant-gene pairs associated with *cis*-eGenes, a genome-wide empirical *P*-value threshold, $p_t$, was defined as the empirical *P*-value of the gene closest to the 0.05 FDR threshold. $p_t$ was then used to calculate a nominal *P*-value threshold for each gene based on the beta distribution parameters (from FastQTL) of the minimum *P*-value distribution $f(p_{\text{min}})$ obtained from the permutations for the gene. Specifically, the nominal threshold was calculated as $F^{-1}(p_t)$, where $F^{-1}$ is the inverse cumulative distribution. For each gene, variants with a nominal *P*-value below the gene-level threshold were considered significant and included in the final list of variant-gene pairs.

As expected, *cis*-eQTL discovery was strongly correlated with sample size in each tissue, as well as the number of expressed genes per tissue with especially testis expressing many more genes than other tissues (**Fig. 2, fig. S6, table S2**). The comparison of *cis*-eQTL discovery across the GTEx pilot (v3), mid-stage (v6) and the current v8 release shows a substantial increase as a result of the improved power (**fig. S6**).

There are a small number of genes for which we do not observe a *cis*-eQTL in any tissue. We used the PANTHER over-representation test (release 20171205) against 21,042 human genes as background to test for enrichment in GO biological processes of different sets of non-eGenes using the GO database release 2018-04-04. Significant GO IDs (Bonferroni-adjusted P-value $<$ 0.05) were selected for analysis with REVIGO to group similar ontological terms. Non-eGenes genes are enriched for processes such as detection of (chemical) stimuli that include olfactory genes that are not well captured in GTEx gene expression data (**fig. S7**).

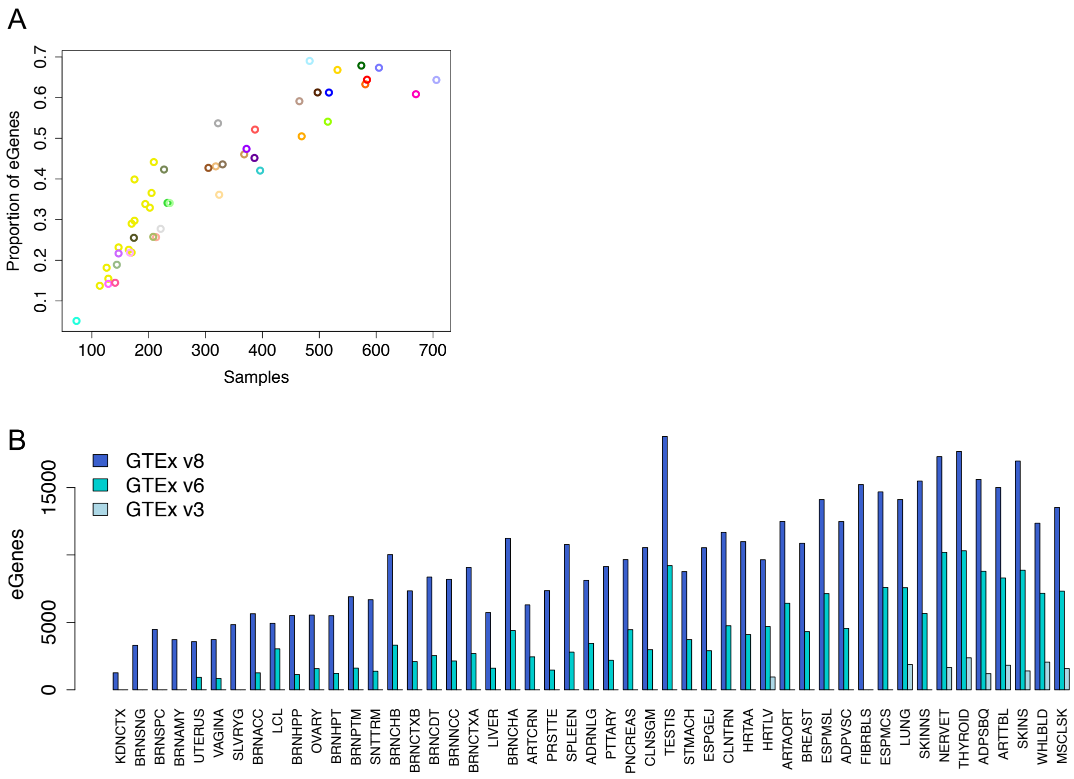

**Fig. S6. Statistics of *cis*-eQTLs**. A) Proportion of eGenes / detected genes across tissues; B) Number of eGenes in the GTEx v3 data release (pilot [2]), v6 data release (mid-stage [15]) and v8 data release.

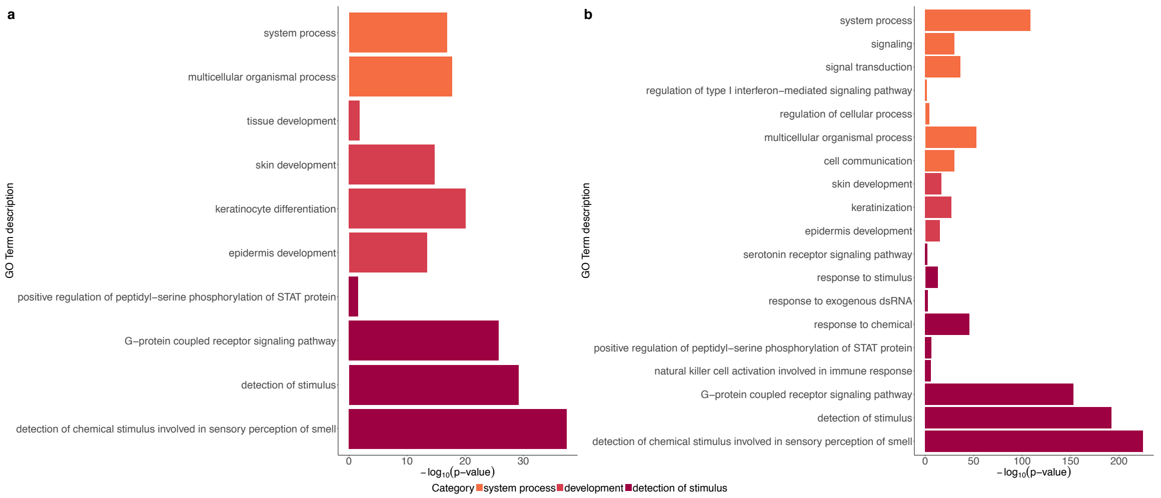

**Fig. S7. Properties of genes that do not have *cis*-eQTLs.** a) Gene Ontology (GO) analysis of tested protein-coding non-eGenes (408 genes), which yielded 11 over-represented GO IDs. b) GO analysis of a more stringent set of protein-coding non-eGenes. Selected genes included those not tested in GTEx (449 genes) or those with a minimum nominal P value across tissues greater than 0.1 (311 genes). Of these stringent 760 non-eGenes, 639 were mapped in the GO analysis. 19 over-represented GO IDs were identified. For both a and b, the x-axis represents the $-\log_{10}$(P-value) resulting from GO analysis. GO IDs are colored by the broader enrichment category to which each corresponds.

#### 4.3 *cis*-sQTL mapping

The overall mapping approach for *cis*-sQTLs was based on and largely similar to the *cis*-eQTL mapping approach described in Section 4.2. *cis*-sQTL mapping was performed with FastQTL, testing for associations with variants within ±1Mb of each gene's TSS, and using the covariates described in Section 4.1 similarly to Section 4.2 as well as 15 PEER factors calculated from the splicing quantifications. We used grouped permutations (`--grp` option) to jointly compute an empirical p-value over all intron clusters of a gene. The top nominal *cis*-sQTL for a gene was defined as the top association among all of its assigned clusters and introns. Empirical p-values were obtained by computing this top association for 1,000-10,000 permutations of the sample labels (`--permute 1000 10000` option). To identify *cis*-sGenes, FDR was computed in the same manner as for *cis*-eQTLs (Section 4.2), including the computation of a sGene-level nominal p-value threshold used to identify all significant variant-intron pairs.

#### 4.4 Independent *cis*-QTL mapping

Multiple independent signals for a given expression phenotype were identified by forward stepwise regression followed by a backwards selection step, with the same approach applied to both *cis*-eQTLs and *cis*-sQTLs. The gene-level significance threshold was set to be the maximum beta-adjusted *P*-value (correcting for multiple-testing across the variants) over all eGenes/sGenes in a given tissue. At each iteration, we performed a scan for *cis*-QTLs using FastQTL, correcting for all previously discovered variants and all covariates used in regular *cis*-QTL mapping. If the beta adjusted *P*-value for the lead variant was not significant at the gene-level threshold, the forward stage was complete and the procedure moved on to the backward stage. If this *P*-value was significant, the lead variant was added to the list of discovered *cis*-QTLs as an independent signal and the forward step moves on to the next iteration. The backwards stage consisted of testing each variant separately, controlling for all other discovered variants. To do this, for each e/sVariant, we scanned for *cis*-QTLs controlling for standard covariates and all other e/sVariants. If no variant was significant at the gene-level threshold the variant in question was dropped, otherwise the lead variant from this scan, which controls for all other signals found in the forward stage, was chosen as the variant that represents the signal best in the full model.

In order to validate the conditional mapping of independent *cis*-eQTLs, we used the orthogonal dap-g approach [18]. It implements a Bayesian variable selection model to identify independent *cis*-eQTLs for each gene. It uses a signal-level posterior inclusion probability (SPIP) to quantify the strength of each association signal, which is represented by SNPs in LD. When the SPIP $>$ 0.90 (i.e., a local false discovery rate $<$ 0.10), a 90% Bayesian credible set of representative SNPs is constructed for the corresponding eQTL. Here, we count the number of 90% credible sets of independent *cis*-eQTLs across tissues. The two methods produced very similar results (**fig. S8**).

We discovered a large number of independent *cis*-eQTLs, especially in tissues with larger sample sizes with better statistical power, and the number of *cis*-eQTLs increases linearly with sample size. The number of independent *cis*-sQTLs is slightly lower, but this might also be a technical artefact of less statistical power (**Fig. 2, fig. S8, table S2**).

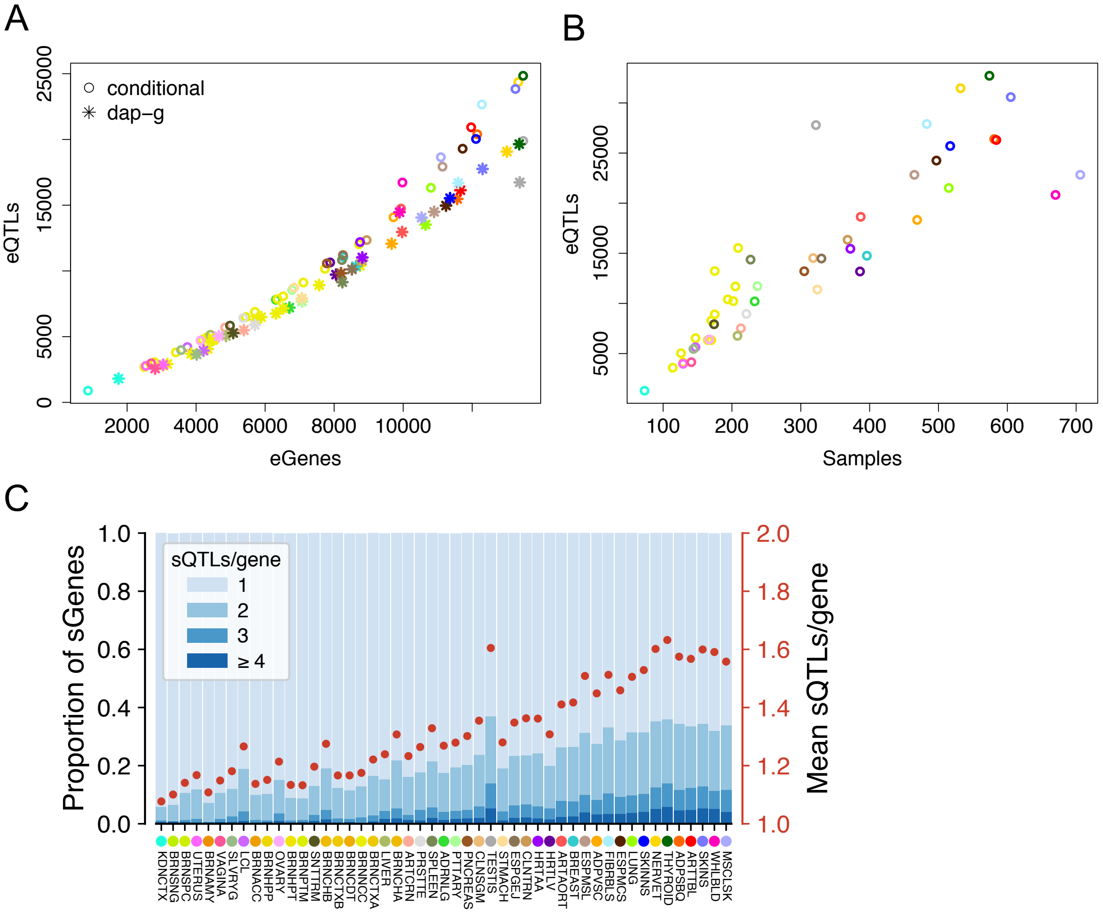

**Fig. S8. Allelic heterogeneity**. A) Comparison of the two methods for estimating allelic heterogeneity; B) number of total *cis*-eQTLs per tissue as a function of sample size; C) allelic heterogeneity of *cis*-sQTLs.

#### 4.5 *trans*-eQTL mapping

For *trans*-eQTL mapping, we used the same covariates as for *cis*-eQTL analysis (Section 4.1). The genotype, expression and covariates were used to map *trans*-eQTLs for all gene-variant pairs not in the same autosomal chromosome, using the `linear_regression()` function in Hail v0.2. We excluded any variant with mappability $< 1$, based on k-mer length $75$, from association testing. Since the number of tests exceed $5 * 10^{11}$ in most tissues, we only saved the pairs that passed a p-value threshold of $10^{-5}$. Candidate *trans*-eGenes were restricted to protein-coding and lincRNA genes, as annotated in GENCODE v26. Finally, we applied the hg38 cross-mapping filter as described in [19] with settings of k-mer length $75$ for exons and $36$ for UTRs, applying this to the filtered set of variant-gene pairs with p-values below $10^{-5}$ to exclude any gene with mappability $< 0.8$ and any variant-gene pair where the target eGene cross-maps with any gene within 1Mb of the variant.

Gene-level FDR was calculated by taking, per tissue, the most extreme P-value per gene across all tested SNPs, multiplying that P-value by $10^6$, in concordance with the effective number of tests on average assumed in genome-wide association studies, and applying Benjamini-Hochberg on those adjusted extreme P-values across genes to control FDR.

#### 4.6 *trans*-sQTL mapping

For *trans*-sQTL mapping, we limited our search to protein coding and lincRNA genes as annotated in GENCODE v26 that had an average 36-mer mappability $\geq$0.8 (we used a shorter k-mer length compared to *trans*-eQTLs due to the higher risk of mapping artifacts with the split reads used to quantify intron excision ratios). To further minimize potential mapping artifacts, we also filtered the analysis freeze VCF used for *cis*-QTLs based on the following exclusion criteria: all variants with missingness in the phased VCF (due to phasing errors or multi-allelic sites; 270,106 and 11,021, respectively); variants with missingness $>$ 0.02 in the unphased VCF (704,228); variants with 75-mer SNP mappability $<$0.9 (1,624,562) [19]; and variants that failed HWE in

| Tissue | Samples | Expr genes | *cis*-eGenes | eGenes $> 1$ eQTLs | Spliced genes | *cis*-sGenes | eGenes $> 1$ sQTLs |
|---|---|---|---|---|---|---|---|
| Adipose_Subcutaneous | 581 | 18329 | 12146 | 5409 | 11536 | 4662 | 1604 |
| Adipose_Visceral_Omentum | 469 | 18358 | 9719 | 3129 | 11818 | 3790 | 1040 |
| Adrenal_Gland | 233 | 18000 | 6311 | 1220 | 11487 | 2070 | 364 |
| Artery_Aorta | 387 | 17999 | 9937 | 3442 | 11377 | 3356 | 882 |
| Artery_Coronary | 213 | 18358 | 4840 | 714 | 11877 | 1876 | 303 |
| Artery_Tibial | 584 | 17716 | 11970 | 5628 | 11048 | 4377 | 1465 |
| Brain_Amygdala | 129 | 18649 | 2794 | 244 | 11601 | 768 | 55 |
| Brain_Anterior_cingulate_cortex_BA24 | 147 | 18739 | 4312 | 562 | 11951 | 1049 | 104 |
| Brain_Caudate_basal_ganglia | 194 | 18825 | 6523 | 1304 | 12020 | 1550 | 179 |
| Brain_Cerebellar_Hemisphere | 175 | 18520 | 7734 | 1960 | 11637 | 2101 | 399 |
| Brain_Cerebellum | 209 | 18656 | 8720 | 2519 | 11576 | 2447 | 533 |
| Brain_Cortex | 205 | 18915 | 7108 | 1639 | 12010 | 1774 | 293 |
| Brain_Frontal_Cortex_BA9 | 175 | 18865 | 5704 | 1010 | 12076 | 1437 | 178 |
| Brain_Hippocampus | 165 | 18796 | 4292 | 530 | 11707 | 1000 | 102 |
| Brain_Hypothalamus | 170 | 19140 | 4189 | 526 | 12242 | 1204 | 108 |
| Brain_Nucleus_accumbens_basal_ganglia | 202 | 18855 | 6359 | 1256 | 12113 | 1599 | 207 |
| Brain_Putamen_basal_ganglia | 170 | 18433 | 5440 | 914 | 11485 | 1169 | 103 |
| Brain_Spinal_cord_cervical_c-1 | 126 | 18849 | 3414 | 348 | 11753 | 965 | 102 |
| Brain_Substantia_nigra | 114 | 18642 | 2500 | 180 | 11496 | 686 | 45 |
| Breast_Mammary_Tissue | 396 | 18923 | 8270 | 2097 | 12277 | 3705 | 981 |
| Cells_Cultured_fibroblasts | 483 | 16763 | 12280 | 6335 | 10631 | 4204 | 1394 |
| Cells_EBV-transformed_lymphocytes | 147 | 16585 | 3747 | 408 | 10595 | 2113 | 398 |
| Colon_Sigmoid | 318 | 18449 | 8252 | 2272 | 11680 | 2909 | 688 |
| Colon_Transverse | 368 | 18787 | 8943 | 2548 | 12265 | 3103 | 730 |
| Esophagus_Gastroesophageal_Junction | 330 | 18206 | 8256 | 2237 | 11456 | 2933 | 683 |
| Esophagus_Mucosa | 497 | 18038 | 11723 | 5057 | 11271 | 3577 | 1027 |
| Esophagus_Muscularis | 465 | 18083 | 11140 | 4536 | 11257 | 3669 | 1145 |
| Heart_Atrial_Appendage | 372 | 17645 | 8751 | 2574 | 11397 | 2725 | 659 |
| Heart_Left_Ventricle | 386 | 16675 | 7885 | 2102 | 10535 | 2135 | 426 |
| Kidney_Cortex | 73 | 18649 | 868 | 23 | 11252 | 468 | 27 |
| Liver | 208 | 17243 | 4415 | 630 | 10330 | 1284 | 197 |
| Lung | 515 | 18977 | 10804 | 3854 | 12390 | 4326 | 1360 |
| Minor_Salivary_Gland | 144 | 18799 | 3564 | 373 | 12354 | 1432 | 171 |
| Muscle_Skeletal | 706 | 16584 | 11092 | 4937 | 9928 | 3759 | 1270 |
| Nerve_Tibial | 532 | 18853 | 13337 | 6635 | 11795 | 4793 | 1688 |
| Ovary | 167 | 18490 | 4126 | 518 | 11705 | 1721 | 261 |
| Pancreas | 305 | 17471 | 7788 | 2117 | 10586 | 2011 | 407 |
| Pituitary | 237 | 19549 | 6781 | 1408 | 12489 | 2525 | 492 |
| Prostate | 221 | 19366 | 5366 | 870 | 12358 | 2132 | 378 |
| Skin_Not_Sun_Exposed_Suprapubic | 517 | 18714 | 12110 | 5234 | 11807 | 4265 | 1345 |
| Skin_Sun_Exposed_Lower_leg | 605 | 18738 | 13254 | 6525 | 11760 | 4717 | 1621 |
| Small_Intestine_Terminal_Ileum | 174 | 19110 | 4982 | 730 | 12621 | 1802 | 235 |
| Spleen | 227 | 18546 | 8225 | 2051 | 11699 | 2492 | 534 |
| Stomach | 324 | 18389 | 6838 | 1499 | 11809 | 2350 | 447 |
| Testis | 322 | 23628 | 13480 | 4640 | 15990 | 7107 | 2621 |
| Thyroid | 574 | 18923 | 13477 | 6727 | 11817 | 4818 | 1729 |
| Uterus | 129 | 18529 | 2545 | 213 | 11766 | 1320 | 156 |
| Vagina | 141 | 18898 | 2701 | 255 | 12320 | 1270 | 134 |
| Whole_Blood | 670 | 15771 | 9979 | 4342 | 9005 | 2735 | 876 |

**Table S2. Summary of *cis*-QTLs per tissue**

the phased VCF separately in EUR and AFR individuals (inferred ancestries) on autosomes/PAR region and EUR females in the NONPAR region (p-value $<$10-8; 2855 total), resulting in a VCF with 8,502,701 variants (out of 10,770,860).

We mapped *trans*-sQTLs using tensorQTL [20], and restricted mapping to the set of variants with MAF $\geq$ 0.05 in each tissue. To enable FDR control based on a single set of genome-wide permutations, we inverse normal transformed the LeafCutter phenotypes. We generated summary statistics for all associations with p-values $< 10^{-5}$, and applied the same cross-mapping filter described for *trans*-sQTLs [19] to exclude variants within ±1Mb of genes that cross-map with candidate *trans*-sGenes. For gene-level FDR control, we applied the following procedure: for each gene, we selected the variant-phenotype pair with the smallest nominal p-value, and computed the beta-approximated empirical p-value based on 100,000 permutations of a standard normal distribution (to avoid inclusion of *cis* effects, an empirical distribution was calculated for each chromosome using variants on all other chromosomes; these permutations were computed once for each tissue). Since multiple intron excision phenotypes were tested for each gene, we then used the beta-approximated empirical CDF $F(x)$ to compute the distribution for the minimum p-value across $k$ phenotypes, which is given by $1-(1-F(x))^k$. Lastly, we applied the Benjamini-Hochberg correction to these p-values across genes, and used a 0.05 FDR threshold per tissue to define the set of *trans*-sQTLs. We further inspected these results for potential artifacts by checking for RNA-seq coverage differences that reflect alternative splicing (**fig. S9**) and by identifying putative *cis*-regulating genes through mediation and colocalization analyses (Sections 11.4.2 & 11.4.3).

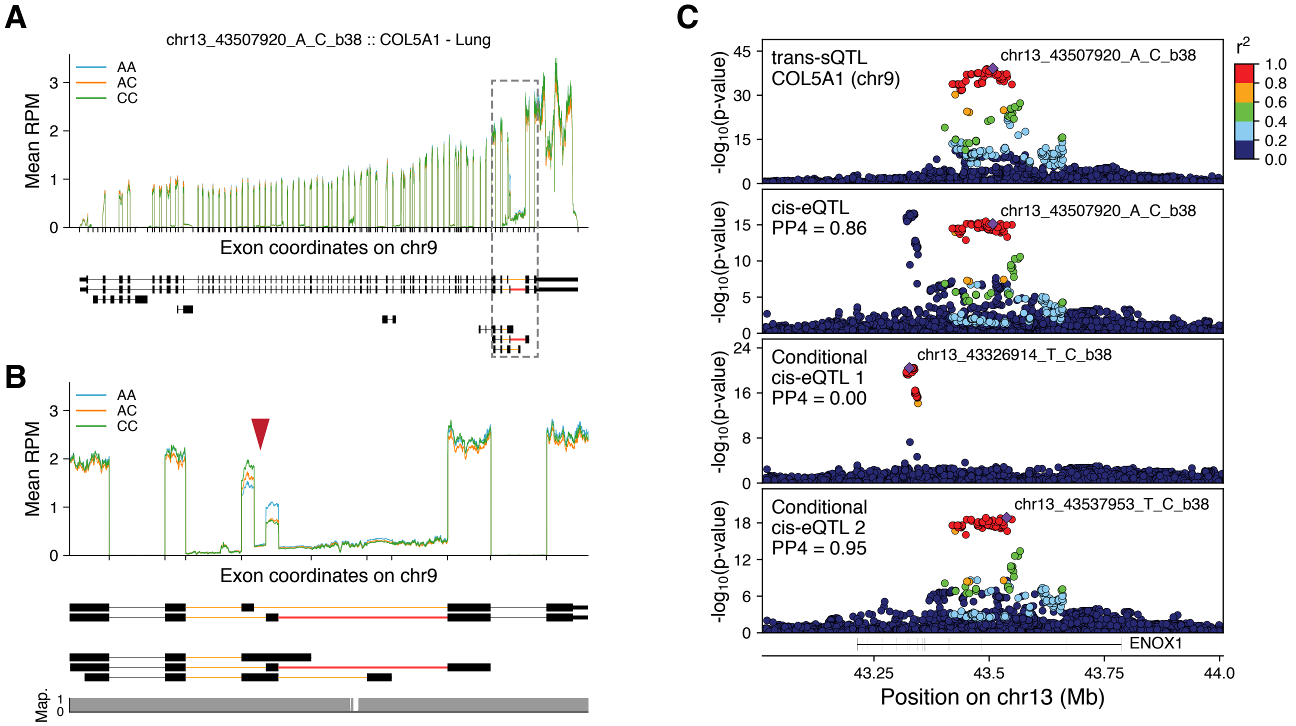

**Fig. S9.** ***Trans*-sQTL example**. A) RNA-seq coverage for *COL5A1* in lung, grouped by genotype for variant rs10047763 (chr13_43507920_A_C_b38). B) Enlarged view of the dashed region from (A), showing the difference in coverage between the two highlighted exons (red arrowhead) across genotypes. The orange and red introns at the bottom indicate the LeafCutter phenotype group tested, with the red intron (shared between two annotated isoforms) excision ratio producing the strongest association. Map.: 36-mer mappability. C) P-value landscapes for the COL5A1 *trans*-eQTL (top panel), the putative *cis*-mediating eQTL for *ENOX1* (second panel), and the two conditionally independent *cis*-eQTLs for *ENOX1* (bottom panels), indicating that only the second independent *cis*-eQTL mediates the *trans* effect. PP4: posterior probability of colocalization from COLOC [21].

### 5 Allele-specific expression

Allelic expression data was produced used the standard alignments, as well as alignments with the WASP filtering strategy to remove reads with allelic mapping bias (Section 3.2). The latter data was used in downstream analyses.

SNP-level ASE data was generated using the GATK ASEReadCounter tool (v3.8-0-ge9d806836) with the following settings: `--U ALLOW_N_CIGAR_READS -minDepth 1 --allow_potentially_misencoded_quality_scores --minMappingQuality 255 --minBaseQuality 10`. Raw SNP level data, consisting of the GATK tool output, were aggregated per subject across all tissues and used to produce analysis files, including only sites with $\geq$8 reads, assigning SNPs to genes, calculating the expected null ratio for each combination of ref/alt allele [22], calculating a binomial p-value by comparing to the expected null ratio, and calculating a multiple hypothesis corrected p-value per sample (subject-tissue) using Benjamini–Hochberg. Furthermore, we flagged sites that lie in low-mappability regions (75-mer mappability $< 1$, with $\leq$2 mismatches allowed for the 75-mer alignment), showed mapping bias in simulation [23], or had no more reads supporting two alleles than would be expecting from sequencing noise alone, indicating a potential genotyping error (FDR $<$ 1%, see [22] for description of test). Note that the genotype warning test cannot distinguish between strong allelic imbalance and a true genotyping error, so this flag should not be used when studying phenomena with mono-allelic expression (e.g., imprinting).

Haplotype-level data was generated using phASER v1.0.1 [24]. First, we complemented the WGS-based phasing (see Section 2.4) by RNA-seq read phasing with phASER, run using all available RNA-seq libraries per subject. These phased genotype data are provided in dbGap. Next, haplotypic expression was calculated using phASER Gene AE 1.2.0 and gene annotations with `--min_haplo_maf 0.05`. Haplotypic expression matrices containing all samples were generated using the `phaser_expr_matrix.py` script. This consists of a single string per sample per gene with the format `HAP_A_COUNT|HAP_B_COUNT`. One matrix was generated using only haplotypes that could be genome-wide phased such that the haplotypes are consistent across genes, according to available phasing data (note that some switch errors are expected over longer distances). Additionally, we provide another data matrix without trying to maintain genome-wide phasing across genes; compared to the previous data this in some cases includes data from (typically rare) variants with high coverage but unreliable genome-wide phasing.

### 6 QTL effect sizes

#### 6.1 *cis* and *trans*-eQTL effect size

*cis*-eQTL effect size was defined as allelic fold change (aFC), the ratio between the expression of the haplotype carrying the alternative eVariant allele to the one carrying the reference allele in $\log_2$ scale [25]. This was calculated for each top eVariant (based on p-value) per gene per tissue using the aFC v0.3 tool [25]. We calculated aFC for *cis*-eQTLs from two data sources: 1) raw gene count data that was that was normalized with DESeq size factors [26] and $\log_2$ transformed, with the aFC arguments `--min_samps 2 and --min_alleles 1` and including the same covariates that were used in the *cis*-eQTL mapping; and 2) haplotype-level ASE data, using the phASER add-on `phaser_cis_var.py` [27] for eQTLs with at least 10 individuals with ASE data and a minimum of 8 reads per individual, and a pseudo-count of 1 added to each the reference and alternative eQTL haplotype counts.

*trans*-eQTL effect size was calculated as allelic fold change, similarly to the expression-based calculation of *cis*-eQTLs. However, the interpretation of allelic fold change for *trans*-eQTLs comes with certain caveats. The allelic fold chance approach assumes a linear (i.e. codominant) effect of the genotype, which is the appropriate biological mechanism in *cis* but not necessarily in *trans*. Since *trans*-eQTL mapping uses a linear model, this assumption is probably not strongly violated, but if it is, the effect size estimates are not necessarily accurate.

The effect size distribution depends on the properties of the discovered eQTLs. In tissues with larger sample sizes, better power allows discovery of eQTLs of smaller effect than in smaller tissues, which is reflected as a larger proportion of eQTLs having more than two-fold effect on expression ($|\text{aFC}| \geq 1$ in $\log_2$ scale) in smaller tissues (**fig. S10**).

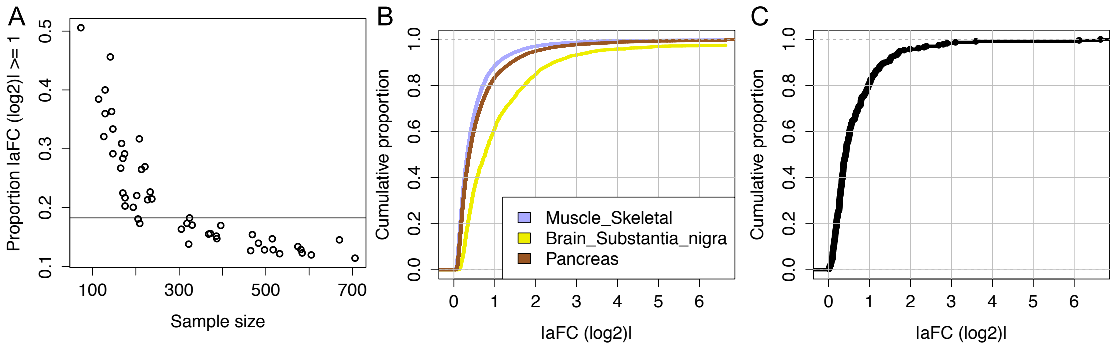

**Fig. S10. Effect size distribution of eQTLs.** A) Proportion of *cis*-eQTLs per tissue with over two-fold effect on gene expression, as a function of sample size. B) Examples of the absolute allelic fold change distribution for *cis*-eQTLs in three tissues of different sample sizes: Muscle Skeletal N = 706, Brain Substantial Nigra N = 114, Pancreas N = 305). C) The absolute allelic fold change distribution for *trans*-eQTLs.

#### 6.2 *cis*-sQTL effect size

The allelic fold change approach is not easily transferable to splicing quantifications. Thus, for *cis*-sQTL effect sizes we simply used the linear regression beta that lacks the biological interpretability of a fold change. Comparison of this to aFC calculated from ASE data of *cis*-QTL heterozygotes indicates the extent to which ASE data reflects different genetic regulatory effects in *cis*. The *cis*-eQTL beta is highly correlated with ASE AFC, indicating that *cis*-eQTLs are strong drivers of allelic imbalance [15]. However, *cis*-sQTLs could cause ASE only in the parts of the transcript that are affected by the splicing change, and this does not manifest in overall ASE data (**fig. S11**).

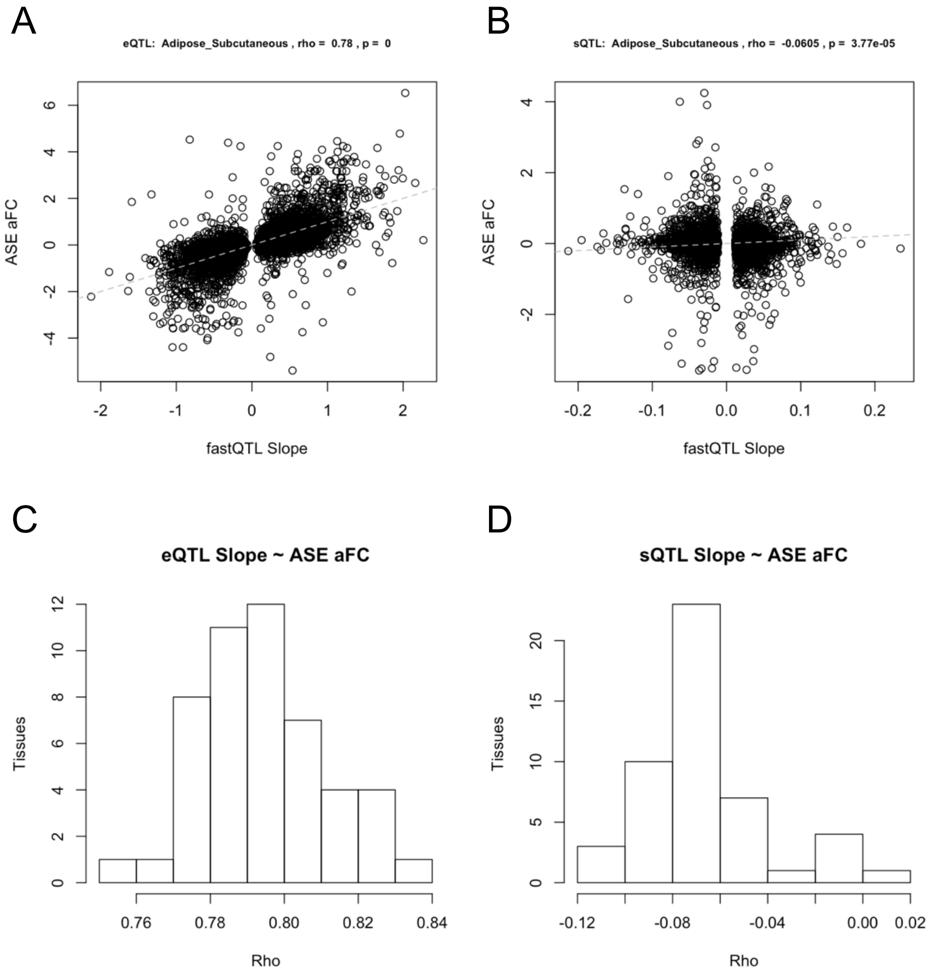

**Fig. S11. Correlation of allelic expression and *cis*-QTL effects.** A-B) Correlation of adipose subcutaneous allelic fold change calculated from allelic expression data. A) *cis*-eQTL allelic fold change calculated from expression data, and B) *cis*-sQTL linear regression beta. The small but significant negative correlation in *cis*-sQTLs is likely due to the direction of the splicing change being difficult to define, since LeafCutter captures complex features of inclusion/exclusion of different parts of the gene. C-D) Distribution of correlation coefficients from different tissues.

##### 6.3 ASE replication of interaction eQTLs

Since ASE data captures *cis*-eQTL effects, they can be used for an orthogonal internal replication analysis. In addition to calculating *cis*-eQTL aFC as a median of allelic imbalance in individuals heterozygous for the eQTL (Section 6.1), it is also possible to analyze this per individual. This is particularly useful for replication of interaction eQTLs (Sections 7, 8, 14), where correlation between the eQTL interaction factor and individual's aFC provides additional validation of the eQTL interaction. For this analysis, we used the phASER haplotype-based ASE data for genes with $\geq$10 eQTL heterozygote individuals with $\geq$8 reads of ASE data per gene. It should be noted that since these data are often sparse, the ASE replication p-values can be poor due to the small number of individuals or due to very noisy individual aFC estimates if read counts are low.

#### 7 Sex-biased *cis*-eQTL mapping

To identify *cis*-eQTLs with a potential sex bias, we tested the set of conditionally independent *cis*-eQTLs for an interaction between sex and genotype. We excluded sex-specific tissues and restricted our analysis to the 44 tissues shared between males and females. We considered variants with MAF$\geq$0.05 in the corresponding tissue, and in total tested 491,693 eQTLs corresponding to 30,121 genes and 273,041 variants across the 44 tissues. Specifically, we fitted the model

$$y_i = \beta_0 + \beta_1 \text{sex} + \beta_2 \text{genotype} + \beta_3 \text{sex} \times \text{genotype} + \lambda \mathbf{C} + \epsilon$$

where $y_i$ is the expression of gene $i$ and the matrix $\mathbf{C}$ contained the same set of covariates and PEER factors used to map eQTLs. To identify sex-biased *cis*-eGenes, we selected the minimum $\beta_3$ p-value for each gene, and applied Bonferroni correction by multiplying this p-value by the number of independent *cis*-eQTLs discovered for the gene. We then computed q-values [17] for gene-level FDR control based on these Bonferroni-corrected minimum p-values per gene, and used a FDR $\leq$ 25% threshold to identify genes with at least one significant sb-eQTL, in each tissue. We used ASE data to validate sb-eQTLs (Section 6.3) by calculating aFC from the ASE data (ASE-aFC) for individuals of each sex, and used the Wilcoxon rank sum test to identify differences in ASE-aFC between males and females (sex $\Delta$aFC). We used the $\pi$1 statistic to quantify validation (**fig. S12** ).

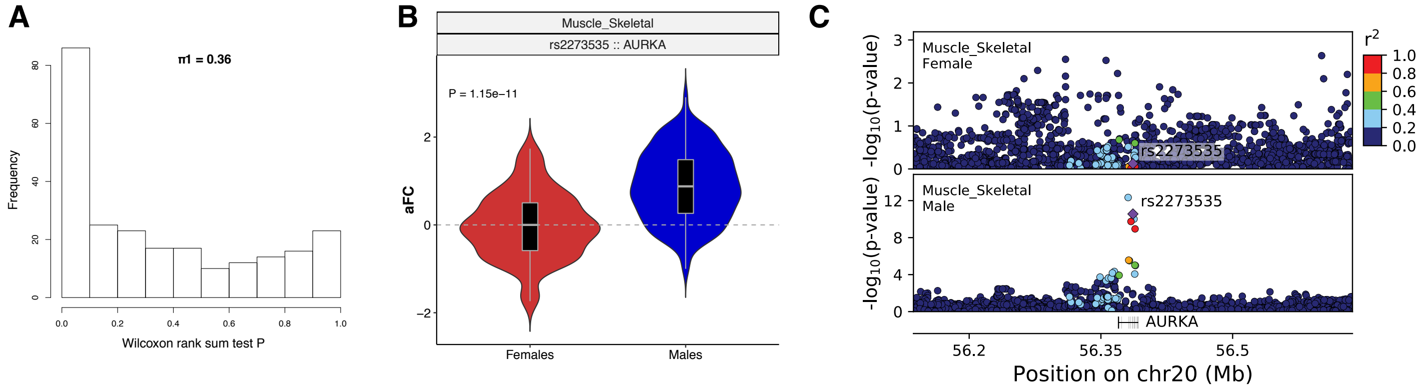

**Fig. S12. Sex-biased eQTLs (sb-eQTLs)**. A) Distribution of sex ΔaFC Wilcoxon rank sum test P-values for significant (FDR $\leq$ 25%) sb-eQTLs and corresponding $\pi$1 statistic. B) Comparison of rs2273535-*AURKA* aFC values between female and male muscle samples (Wilcoxon rank sum test). C) Regional plot of *AURKA* region. The significance of the muscle eQTL signal for genetic variants within 250 kilobases of the TSS of the *AURKA* gene is plotted for females and males. Variants are color-coded according to linkage disequilibrium ($r^2$) with the lead AURKA sb-eQTL variant rs2273535.

### 8 Population-biased *cis*-eQTL mapping

We conducted a systematic approach to characterize population differences in effect sizes of *cis*-eQTLs in individuals of European and African ancestry. We defined European Americans (EA, n = 588) and African Americans (AA, n = 86) as the subset of self-reported White and Black or African American individuals that grouped together tightly according to genotype principal components 1 and 2. We restricted our analysis to 31 tissues with more than 20 individuals from both groups.

We aimed to find significant protein-coding or lincRNA eGenes in GTEx where the *cis*-eQTL effect size is different between EA and AA, which we call population-biased eQTLs (pb-eQTLs). For each tissue, we analyzed the lead *cis*-eVariants for all significant eGenes with an additional filter of MAF $>$ 10% in both groups. We measured *cis*-eQTL effect size in EA and AA using the allelic fold change (aFC), which is robust to differences in allele frequency and expression level. We applied it with and without covariates. Of note, to obtain separate covariates for EA and AA samples we processed gene expression data separately for EA and AA individuals, as described in Section 3.4 and 3.5.1, using 5 PEERs for $<$ 40 samples and 10 PEERs for $\geq$ 40 and $<$ 70 samples. We excluded gene-variant pairs if aFC using covariates was not within the confidence interval for aFC based on without covariates estimates or vice versa, and constrained aFC to $\pm\log_2(100)$. To test for the significance of $\Delta$aFC (difference of aFC estimates between EA and AA), we permuted the ancestry group labels 100,000 times. We calculated the permutation p-value as the proportion of permuted $\Delta$aFC as extreme as or more extreme than the true $\Delta$aFC, followed by Benjamini-Hochberg correction to account for multiple testing, using FDR < 25%.

We formed a conservative high confidence set of pb-eQTLs by using stringent filters to remove differences potentially explained by LD or other artefacts summarized in **fig. S13**. The set included 1) pb-eQTLs where the eQTL effect in EA and AA was consistent but of different magnitude, or 2) pb-eQTLs where there was no strong eQTL effect in one of the populations. Firstly, eQTL mapping was performed separately in EA and AA groups on separately processed gene expression data and covariates to find the eVariants with lowest P-values in each group, denoted as $\mathrm{eVariant_{EA}}$ and $\mathrm{eVariant_{AA}}$, respectively, as well as in the standard eQTL mapping of the entire sample ($\mathrm{eVariant_{all}}$). For 1) all the following criteria needed to be meet: a) $\mathrm{eVariant_{EA}}$ and $\mathrm{eVariant_{AA}}$ are in strong LD ($r^2 > 0.6$) with the $\mathrm{eVariant_{all}}$, b) difference in effect size replicates using $\mathrm{eVariant_{EA}}$ and $\mathrm{eVariant_{AA}}$ (permutation P-value of $\Delta$aFC is $<$ 0.05 for both), and c) $\Delta$aFC is similar for $\mathrm{eVariant_{all}}$, $\mathrm{eVariant_{EA}}$ and $\mathrm{eVariant_{AA}}$ (maximum difference of 1, or even stronger effect in the right direction for $\Delta$aFC for $\mathrm{eVariant_{EA}}$ or $\mathrm{eVariant_{AA}}$). For 2) the following criteria needed to be meet: a) very low LD ($r^2 < 0.1$) between $\mathrm{eVariant_{EA}}$ and $\mathrm{eVariant_{AA}}$ and $\mathrm{eVariant_{all}}$, and b) no strong eQTL signal in either EA or AA (nominal P-value of $\mathrm{eVariant_{EA}}$ and $\mathrm{eVariant_{AA}}$ $>$ 0.001).

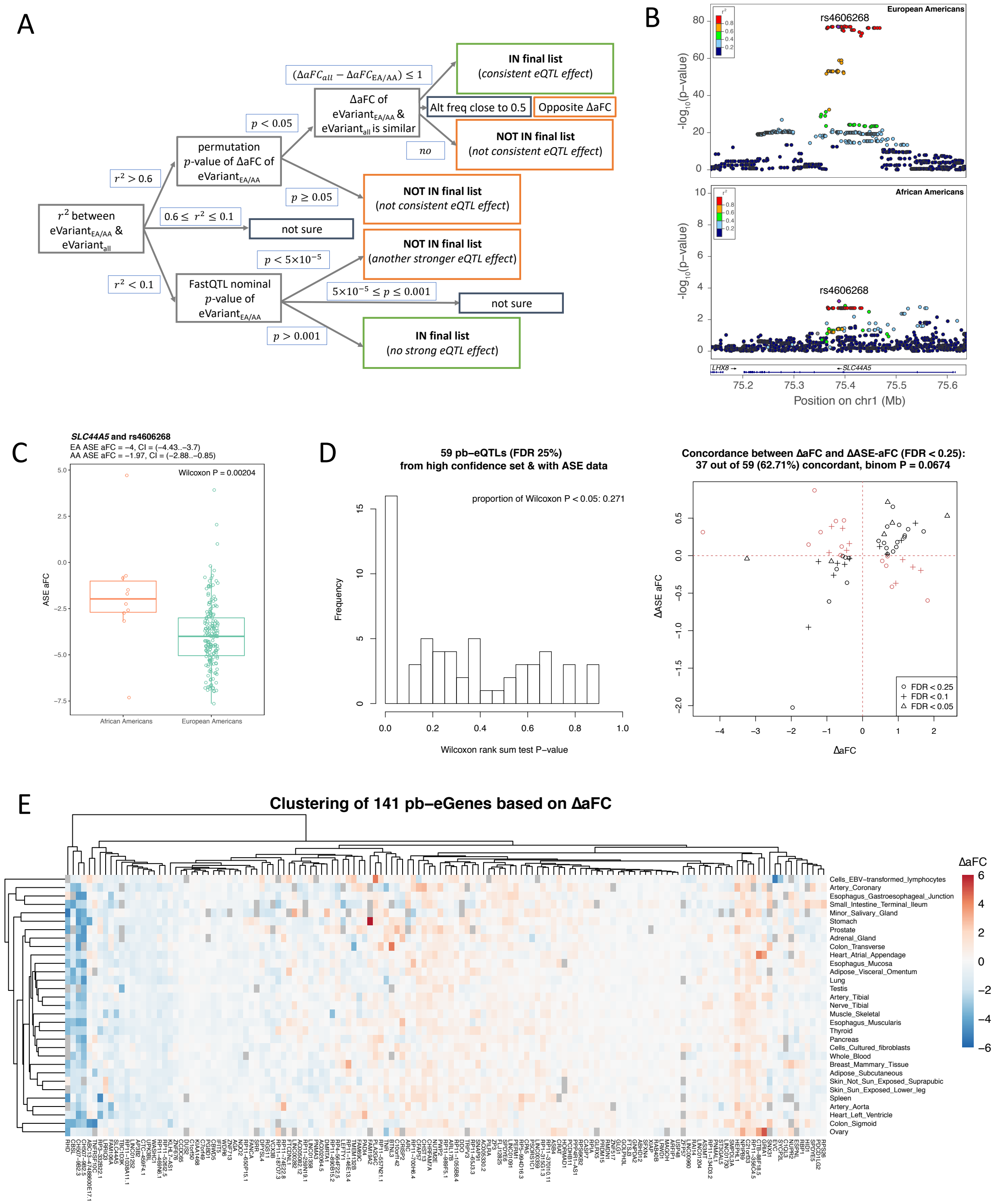

**Fig. S13. Population-biased *cis*-eQTLs (pb-eQTLs).** A) Filtering steps to compose the high confidence set of pb-eQTLs that are not artefacts of LD differences between European (EA) and African Americans (AA) (see text for description). B) Regional plot of a pb-eQTL association in esophagus mucosa for *SLC44A5*. Genetic variants within 250 kilobases of the TSS of the *SLC44A5* gene are plotted for EA and AA, with LD is calculated using the respective population group. rs460628 is the eVariant with lowest P-value in EA and AA. C) Validation of pb-eQTL for *SLC44A5* in esophagus mucosa using ASE data. The box plots show the ASE aFC (x-axis) distribution in EA and AA. D) Validation of pb-eQTLs using ASE aFC. Histogram of the Wilcoxon P-values testing the difference in ASE aFC between EA and AA (on the left) and scatter plot of ΔaFC and ΔASE-aFC (on the right). E) Tissue sharing of pb-eQTL signals. ΔaFC was calculated for every high confidence pb-eGene across 31 tissues (choosing one variant per eGene). Tissue sharing patterns of ΔaFC are illustrated on a heatmap. Euclidean distance was used as the distance measure, with a complete-linking clustering method. Grey tiles indicate cases where it was not possible to calculate aFC for a given pb-eGene and pb-eVariant pair in a specific tissue due to a low number of samples or if the aFC estimate hit the cap value ($\pm \log_2(100)$).

We used ASE data to validate pb-eQTLs, as described in Section 6.3. We used the Wilcoxon rank sum test to test for the difference in ASE aFC between EA and AA. To quantify the validation rate, we calculated the proportion of pb-eQTLs with Wilcoxon p-value $< 0.05$, and estimated the concordance between $\Delta$aFC and $\Delta$ASE-aFC (**fig. S13**).

### 9 Genomic annotation data

The standard functional annotation was performed on all variants in the pre-QC WGS vcf using Ensembl's Variant Effect Predictor (VEP) and Loss-Of-Function Transcript Effect Estimator (LOFTEE) (VEP v85, GENCODE v26) implemented in Hail v0.1. The annotations were added to a sites-only VCF with all 866 samples pre-sample QC, and a total of 69,763,935 sites (including monomorphic sites).

For regulatory annotation of the variants, we used primarily the Ensembl Regulatory Build [28] that has compiled and re-analyzed data from ENCODE, Epigenomics Roadmap, and other projects. This includes 1) annotation of regulatory features (enhancer, promoter, etc) and their activity in each analyzed tissue and cell type; 2) peaks of transcription factor binding and DNAseI hypersensitivity; and 3) transcription factor motif overlap.

In analyses of tissue-sharing of regulatory activity, the Epigenomics Roadmap data and chromatin state annotation provided a substantially better overlap with GTEx tissues than chromatin states available in the Regulatory Build. Thus, in some analyses, as indicated below, we used the ROADMAP core 15-state model (https://egg2.wustl.edu/roadmap/web_portal/chr_state_learning.html).

We matched the ENCODE and Epigenomics Roadmap tissues and cell types to GTEx tissues as described in table S3.

| GTEx tissue | Epigenomics roadmap biospecimen | ENCODE biospecimen |
|---|---|---|
| Adipose_Subcutaneous | Adipose Nuclei (E063) | NA |
| Adipose_Visceral_Omentum | Adipose Nuclei (E063) | NA |
| Adrenal_Gland | Fetal Adrendal Gland (E080) | NA |
| Artery_Aorta | Aorta (E065) | NA |
| Brain_Anterior_cingulate_cortex_BA24 | Brain Cingulate Gyrus (E069) | NA |
| Brain_Caudate_basal_ganglia | Brain Anterior Caudate (E068) | NA |
| Brain_Cerebellum | NA | astrocyte of the cerebellum |
| Brain_Cortex | Brain Angular Gyrus (E067),<br>Brain Inferior Temporal Lobe (E072),<br>Brain Dorsolateral Prefrontal Cortex (E073) | SK-N-MC |
| Brain_Frontal_Cortex_BA9 | Brain Inferior Temoral Lobe (E072),<br>Brain - Dorsolateral Prefrontal Cortex (E073) | NA |
| Brain_Hippocampus | Brain Hippocampus Middle (E071) | NA |
| Brain_Spinal_cord_cervical_c-1 | NA | astrocyte of the spinal cord |
| Brain_Substantia_nigra | Brain Substantia Nigra (E074) | NA |
| Breast_Mammary_Tissue | Breast Myoepithelial Primary Cells (E027) | T47D |
| Cells_EBV-transformed_lymphocytes | Lymphoblastoid Cells (E116) | NA |
| Colon_Sigmoid | Sigmoid Colon (E106) | NA |
| Colon_Transverse | Colonic Mucosa (E075),<br>Colon Smooth Muscle (E076) | DLD1 |
| Esophagus_Gastroesophageal_Junction | Esophagus (E079) | NA |
| Esophagus_Mucosa | Esophagus (E079) | NA |
| Esophagus_Muscularis | Esophagus (E079) | NA |
| Heart_Atrial_Appendage | Right Atrium (E104) | NA |
| Heart_Left_Ventricle | Left Ventricle (E095) | NA |
| Kidney_Cortex | Fetal Kidney (E086) | Caki2 |
| Kidney_Medulla | Fetal Kidney (E086) | NA |
| Liver | Liver (E066) | endothelial cell of<br>hepatic sinusoid |
| Lung | Lung (E096) | NCI-H460 |
| Muscle_Skeletal | Skeletal Muscle Male (E107),<br>Skeletal Muscle Female (E108) | SJCRH30 |
| Ovary | Ovary (E097) | NA |
| Pancreas | Pancreas (E098) | Panc1 |
| Prostate | NA | LNCaP clone FGC |
| Skin_Not_Sun_Exposed_Suprapubic | NHDF-Ad Adult Dermal Fibroblast Primary Cells (E126) | NA |
| Skin_Sun_Exposed_Lower_leg | NA | RPMI-7951 |
| Small_Intestine_Terminal_Ileum | Small Intestine (E109) | NA |
| Spleen | Spleen (E113) | NA |
| Stomach | Stomach Mucose (E110), Stomach Smooth Muscle (E111) | NA |
| Uterus | NA | endometrial microvascular<br>endothelial cells |
| Whole_Blood | Primary mononuclear cells from peripheral blood (E062) | NA |

**Table S3. Pairing of GTEx, ENCODE and Epigenomics Roadmap tissues and cell lines**.

### 10 Fine mapping of *cis*-eQTLs

#### 10.1 Fine mapping methods

Three fine-mapping methods were applied to the *cis*-eQTL data to produce estimates of the causal SNPs: CaVEMaN, CAVIAR and dap-g. The methodologies behind these three methods have been documented elsewhere [18, 29, 30], but in brief: CaVEMaN is based on a non-parametric resampling approach where the properties of causal variants are estimated from simulation studies; the CAVIAR method estimates causal posterior probabilities for specific variants using eQTL summary statistics and the linkage structure in the GTEx genotype data; while dap-g is a computational algorithm designed for Bayesian multi-SNP genetic association analysis which employs a spike-and-slab prior model to select potential multiple independent *cis*-eQTLs in eQTL mapping, exploring highly plausible association models in an efficient and fully automatic way.

The CaVEMaN and CAVIAR approaches used the results of the *cis*-eQTL mapping to identify a set of genes in each tissue with a *cis*-eQTL, and the methods were only run on these sets of genes. CaVEMaN further used the conditional analysis to identify the number of independent *cis*-eQTLs for each gene (as in Section 4.4), and used the methodology previously designed to create "single signal" phenotypes. In this way for each independent *cis*-eQTL identified by the conditional analysis, CaVEMaN reports the most likely causal variant along with the causal probability. CAVIAR maps independent *cis*-eQTLs as part of the algorithm, for each gene it reports the expected number of *cis*-eQTLs and for each gene-variant pair in the *cis*-window it reports the probability that variant has a causal effect on the expression of that gene. Dap-g in contrast identifies all *cis*-eQTLs as part of the algorithm and thus was applied to all genes. For each gene within each tissue, clusters of variants are identified, corresponding to independent *cis*-eQTL signals (see Section 4.4). The causal probability of each cluster is calculated by summing probabilities of individuals variants. Then, a set of significant clusters, controlling the local false discovery rate at 5%, was called by taking the largest set of clusters for which the mean causal probability is greater than 95%.

**Fig. S14** shows the size of the 90% credible set for dap-g: the number of variants necessary to have a greater than 90% probability that the causal variant is among them. The distribution is highly long tailed, in some eQTLs including thousands of SNPs. However, for most eQTLs the credible set is small, and the median size is 6 variants.

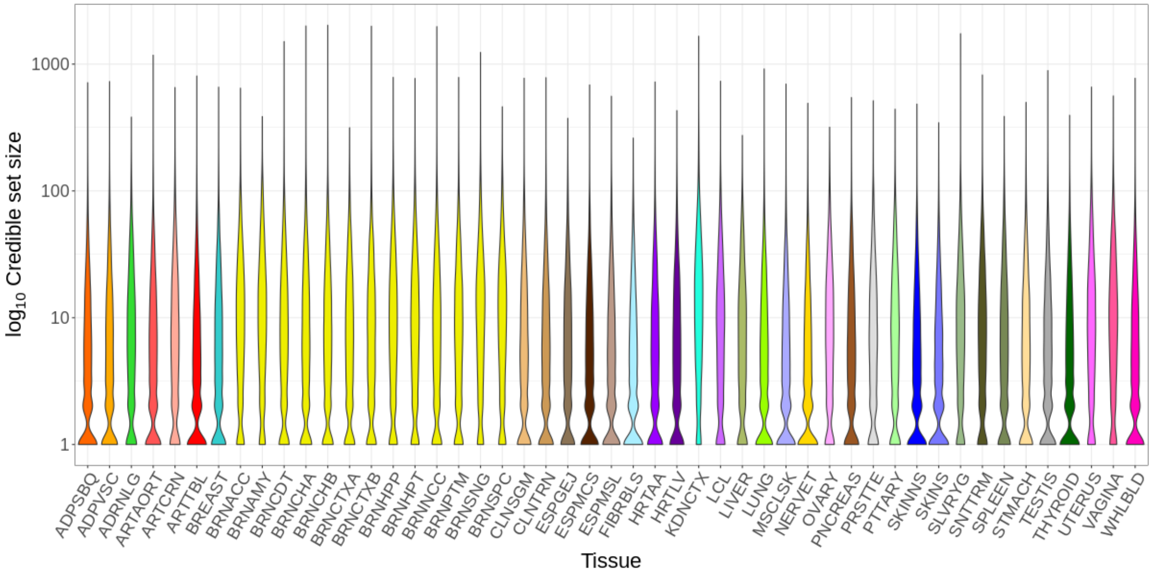

**Fig. S14. Fine-mapping credible sets**. The size of the 90% credible set for *cis*-eQTLs of each tissue based on dap-g.

The consensus set of high confidence causal variants was created by extracting the most likely causal variant for each of the independent eQTLs for both CaVEMaN and dap-g, and the most likely causal variant for CAVIAR. The consensus set consisted of the set of gene-tissue-variant trios where the causal probability was greater than 0.8 in all three methods (**fig. S15**). We identified 24,740 tissue-gene-variant trios that have high causal probability in all three methods.

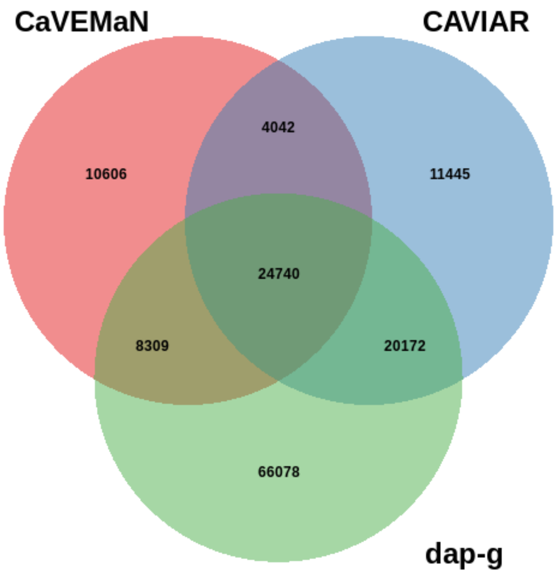

**Fig. S15. Fine-mapping consensus set**. The numbers and overlap of causal gene-tissue-variant trios with causal probability greater than 0.8 according to the different methods.

#### 10.2 Experimental validation

For experimental validation of the *cis*-eQTL fine-mapping results, we used two large-scale screens of regulatory variants, MPRA data set from lymphoblastoid cell lines [31] and SuRE data set from K562 and HepG2 cell lines [32]. For single SNP analysis, we annotated the top associated SNP in an eGene as its sole causal eQTL. Both experimental validation datasets (MPRA and SuRE) utilized p-values to characterize the success of validation for individual SNPs. We employed an EM algorithm (implemented in the software package torus [33]) to infer the latent validation status based on observed p-values and explicitly incorporate the causal eQTL annotations from various computational approaches. This analysis yielded an estimate of log-odds ratio and a corresponding 95% confidence interval for each computational annotation, which quantifies the increased likelihood of an annotated SNP being validated in each experiment in contrast to an unannotated SNP.

#### 10.3 Fine-mapped *cis*-eQTL for *CBX8*

eQTL variants are enriched in *cis*-regulatory elements and can co-localize with GWAS loci, and in such loci fine-mapping of the *cis*-eQTL can help in pinpointing the causal genetic variant(s) and the functional mechanism(s). With fine-mapping approaches narrowing our focus to a set of variants which are likely causing the *cis*-eQTL effects, functional annotations can help us suggest mechanisms for those variants.

First, we annotated fine-mapped variants to find loci where at least one variant overlaps a TF binding site and disrupts the TF motif. To this end, transcription factor binding motif information was downloaded from HOCOMOCO v11 and was filtered only for motifs with an A or B quality score (265 TF/motif pairs). Motifs and reverse complement motifs were translated into regex expressions, and lowercase (lower certainty) motif characters were allowed to be any base. We searched the region within 50 bases of each fine-mapped SNP for each motif and reverse complement motif using grep, as well as the same region with the alternative instead of reference allele.

We decided to focus on a potential transcription factor (TF) mechanism of an *cis*-eQTL for the *CBX8* (Chromobox 8, ENSG00000141570) gene. It has significant *cis*-eQTLs in multiple tissues, and the *cis*-eQTL with the largest effect size is found in lung. The lung *cis*-eQTL signal for *CBX8* co-localized with two GWAS traits: UKB birth weight and BCAC overall breast cancer (EUR) (ENLOC rcp=0.674, 0.678, respectively). These GWAS signals had p values of 2.7e-5 and 4.1e-6, respectively, and another study has found a genome-wide significant breast cancer association in this locus [34].

Caviar detected three SNPs in the 90% confidence set for the *CBX8* Lung *cis*-eQTL: rs9905914, rs1105820, and rs9896202 (**Fig. 3**). ChIP-seq peak and transcription factor (TF) overlap is depicted for each SNP and its 50-base-flanking region (**fig. S16**). rs9896202 is the only SNP that overlaps both a motif and a ChIP-seq peak for the same TF: *EGR1* (Early growth response 1). In order to further analyze the potential role of *EGR1* in the activity of this *cis*-eQTL, we investigated the relationship between eQTL effect size and *EGR1* expression across tissues. If differential binding of *EGR1* to the reference and alternative alleles is causing the *cis*-eQTL effect on *CBX8* expression, it is reasonable to expect that there will be a relationship between *cis*-eQTL effect size and *EGR1* activity or its proxy expression level across tissues. For rs1105820 we calculated effect sizes in each tissue using the allelic fold change (aFC) statistic. We calculated median *EGR1* expression (TPM) across individuals in each tissue, and then the Spearman correlation of median *EGR1* expression with the *cis*-eQTL aFC across tissues. The two measures were significantly correlated (**Fig. 3**; Spearman rho=-0.69, p=1.3e-7). Thus, fine-mapping and functional information suggest that the *CBX8* eQTL may be regulated by *EGR1*. The *CBX8 cis*-eQTL colocalizes with a breast cancer GWAS signal, and *CBX8* has been implicated in various cancers, including breast cancer [35, 36, 37]. Taken together, these data offer the hypothesis that -transregulation by *EGR1* could be regulating the genetic effect of the *CBX8* locus on breast cancer.

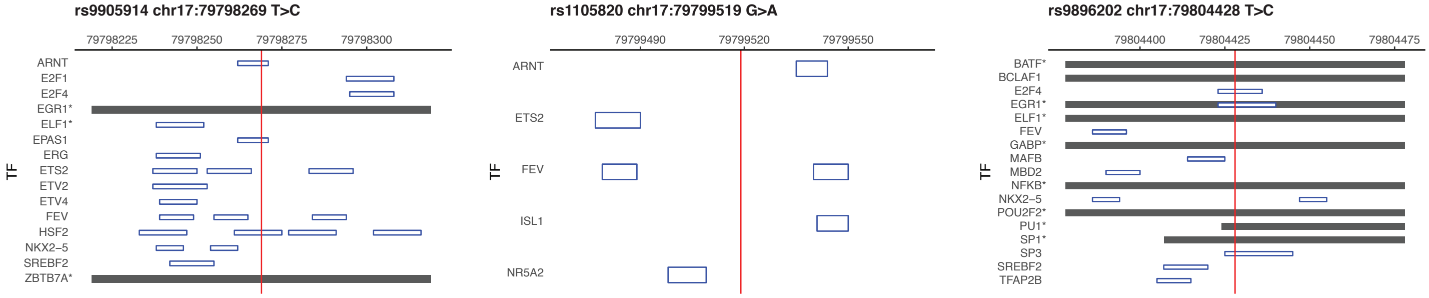

**Fig. S16. Fine-mapping and functional analysis of *CBX8 cis*-eQTL**. Transcription factor ChIP-seq peak (dark gray bar) and motif (blue box) overlap is depicted for three fine-mapped SNPs (red line) for the *CBX8* lung *cis*-eQTL, with 50 bases of flanking region on either side of the SNP. ChIP-seq bars are collapsed across experiments for the same transcription factor. *denotes a transcription factor for which ChIP-seq peak and motif information were both available. rs9896202 (right) is the only SNP that overlaps both a motif and a ChIP-seq peak for the same TF, *EGR1* (Early growth response 1).

### 11 Functional mechanisms

#### 11.1 Enrichment in genomic annotations

Functional enrichment analyses were performed using torus [33], with the annotations described in Section 9 and the command `torus -d ${qtl_statistics} -annot ${annotation_file} -est --fastqtl`. For enrichment of *trans*-eQTL variants, we combined the 162 intra-chromosomal variant-gene pairs discovered at 0.05 FDR across tissues, selected a random set of 5 million total background variants matched to the MAF of each variant (binning variants in the VCF into 50 bins between 0.01 and 0.5 MAF), excluding variants on the same chromosome as the gene in each pair, and computed the p-values and effect sizes as described in Section 4.5 in the corresponding tissue. For *trans*-eQTL enrichment, the option `--no_dtss` was added to ignore distance to the TSS.

#### 11.2 *cis*-eQTL-sQTL overlap

We investigated the overlap of *cis*-eQTLs and *cis*-sQTLs by comparing the overlap of the 90% credible sets of likely causal variants based on CAVIAR. For all the genes that had both an eQTL and and sQTL in *cis*, we calculated the proportion (# eVariants in eQTL & sQTL credible sets)/ (# eVariants in the eQTL credible set).

The cross-tissue median of mean overlap was 0.12. We observed that across the tissues, a median of 46.2% (range 0.394..0.549) of eQTL credible sets had zero overlap with likely causal sQTL variants. This proportion was inversely correlated with effect size (rho = -0.72, p = 1.9e-09) probably due to better power of defining the credible sets in larger tissues. **Fig. S17** shows the cumulative distribution of sharing proportions for three tissues of different sizes. These results show that variants affecting gene expression and splicing are mostly distinct.

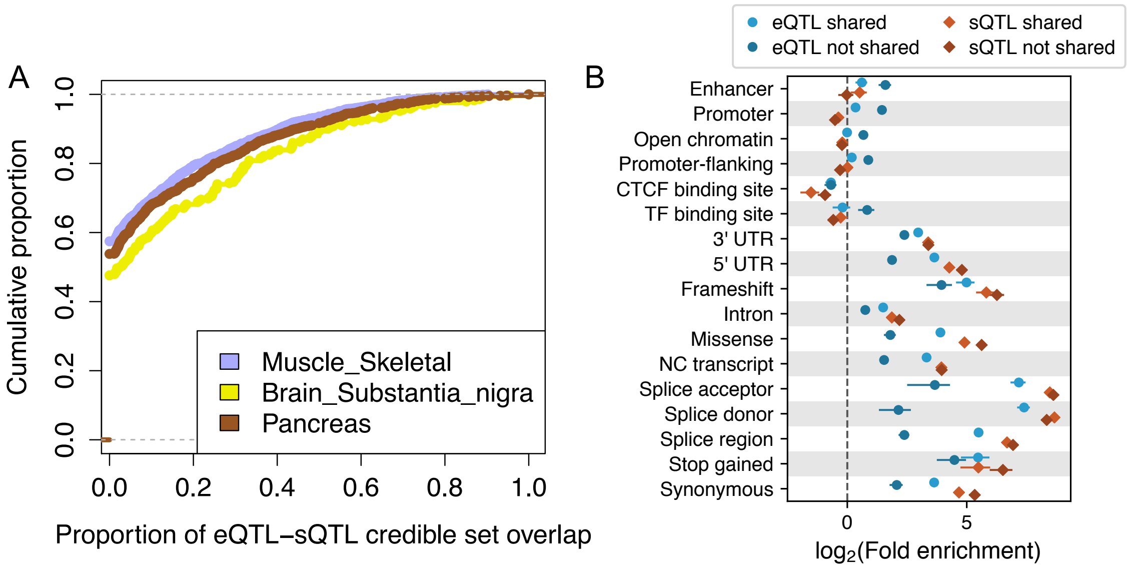

**Fig. S17. Overlap and functional enrichment of *cis*-eQTLs and *cis*-sQTLs.** A) The cumulative distribution of (# eVariants in eQTL & sQTL credible sets)/ (# eVariants in the eQTL credible set) for genes that have both an eQTL and an sQTL. Three tissues of different sample sizes are shown (Muscle Skeletal N = 706, Brain Substantial Nigra N = 114, Pancreas N = 305). B) *cis*-QTL enrichment in functional annotations for potentially shared (lead variant r2 > 0.8) and distinct (r2 < 0.2) *cis*-eQTLs and *cis*-sQTLs.

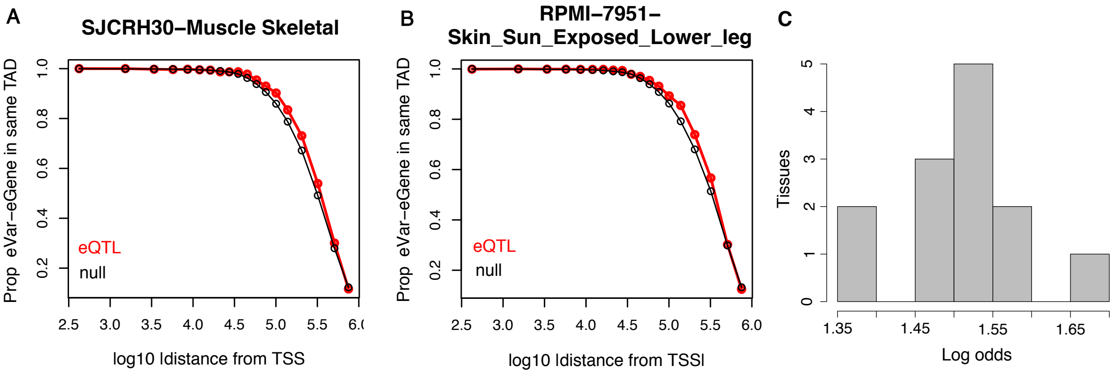

**Fig. S18. *cis*-eQTL enrichment in topology-associated domains (TADs).** A-B) Examples of the proportion of *cis*-eQTLs and null variants that occur in the same TAD as their target gene, as a function of distance from TSS. The title indicates the ENCODE cell line and the GTEx tissue name. C) Distribution of log odds of *cis*-eQTL enrichment in the same TAD, compared to null variants and including distance from TSS in the model, and calculated for each tissue with a matching ENCODE cell type.

#### 11.3 TAD enrichment

We analyzed the enrichment of independent *cis*-eVariant-eGene pairs being in the same topology associated domain (TAD). TAD data based on Hi-C was downloaded from ENCODE using all data released prior to May 2016, with non-overlapping single-scale TADs called by ENCODE. The coordinates were lifted over from hg19 to hg38 by the UCSC liftOver tool. We matched 13 pairs of ENCODE cell types to GTEx tissues (**table S3**). Within each tissue, we compared all the top variants of each independent *cis*-eQTL to a null of a random selection of 1M variants per tissue, selected from the 1Mb window for the genes that were tested for *cis*-eQTLs in that tissue. **Fig. S18** shows examples of the proportion of variants that are within the same TAD for real eQTLs and for the null, demonstrating that due to the large size of TADs, only at distances >100kb are a reasonable proportion of variant-gene pairs not within the same TAD. We analyzed the enrichment of eQTLs being in the same TAD with logistic regression: SameTAD eQTL + |TSSdistance| + eQTL*|TSSdistance|, where SameTAD is an indicator for whether a variant-gene pair is in the same TAD, eQTL is an indicator whether the variant-gene pair is an eQTL or a null, and |TSSdistance| is the absolute distance between the variant and the gene's TSS.

$$SameTAD = \beta_0 + \beta_1 eQTL + \beta_2|TSSdist| + \beta_3 eQTL \times |TSSdist| + \epsilon$$

where $SameTAD$ and $eQTL$ are indicator variables for whether a variant-gene pair is in the same TAD or if it's a true eVariant-eGene pair. $|TSSdist|$ is the absolute distance between the variant and the gene's TSS.

#### 11.4 *cis*-QTL contribution to *trans*-QTLs

##### 11.4.1 *cis*-QTL enrichment among *trans*-QTLs

To test the enrichment of *trans*-e/sGenes that have *cis*-QTLs, we took the *trans*-e/sGenes at FDR $< 0.05$, and analyzed their lead e/sVariant. A set of SNPs were randomly sampled to match for MAF of the *trans*-e/sVariants. We checked whether each of these SNPs was a) tested in *cis* analysis in the matched tissue and b) identified as a *cis*-QTL with any gene in the matched tissue.

##### 11.4.2 *trans*-QTL mediation analysis

We tested mediation for *trans*-QTLs that also had a *cis*-QTL for the corresponding lead *trans*-eVariant in the matched tissue using two-stage least squares (TSLS). We first performed ridge regression ($\alpha = 100$) to estimate the effect of genetic variation on the *cis*-eGene or sGene. Next, we used the learned regression coefficients to predict the expression or intron excision ratio of the *cis*-e/sGene for each individual, followed by a second regression to calculate the causal effect size of the *cis*-e/sGene's predicted expression on the *trans*-e/sGene's measured expression or intron excision ratio. These steps were computed as follows:

$$\begin{aligned}
\mathbf{x} &= {\beta_{cis}}^T Z + \epsilon_1 \\
\hat{\mathbf{x}} &= {\beta_{cis}}^T Z \\
\mathbf{y} &= \beta_{TSLS}\hat{\mathbf{x}} + \epsilon_2
\end{aligned}$$

where $Z$ is the matrix of all variants within 1Mb of *cis*-e/sGene TSS, $\mathbf{x}$ is *cis*-e/sGene expression level or intron excision ratio, $\mathbf{y}$ is *trans*-e/sGene expression level, and $\hat{\mathbf{x}}$ is the predicted expression of the *cis*-e/sGene. Note that the gene expression values $\mathbf{x}, \mathbf{y}$ were orthogonalized with respect to the covariates used in association mapping. A matched set of $\beta_{TSLS}$ statistics were generated with permuted *trans*-e/sGene levels, using $100$ permutations per real trio, and FDR was assessed based on these empirical p-values using q-values [17]. For splicing QTLs, the $\beta_{TSLS}$ p-values were adjusted by the number of phenotypes available for the sGene. The proportion of *trans*-eQTLs that are significant *cis*-eQTLs or mediated by *cis*-eQTLs is shown in **Fig. 4D**, the proportion of *trans*-eQTLs that are significant *cis*-sQTLs or mediated by *cis*-sQTLs is shown in **fig. S19A**, and the proportion of *trans*-sQTLs that are significant *cis*-eQTLs or mediated by *cis*-eQTLs is shown in **fig. S19B**.

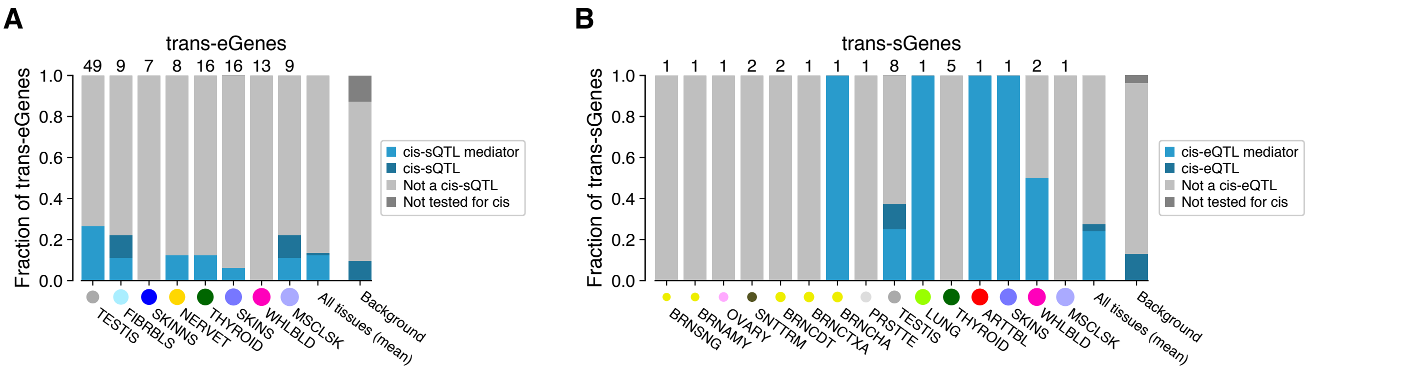

**Fig. S19. *cis*-QTL mediation of *trans*-QTL signals.** A) Proportion of *trans*-eQTLs that are significant *cis*-sQTLs or mediated by them. B) Proportion of *trans*-sQTLs that are significant *cis*-eQTLs or mediated by them.

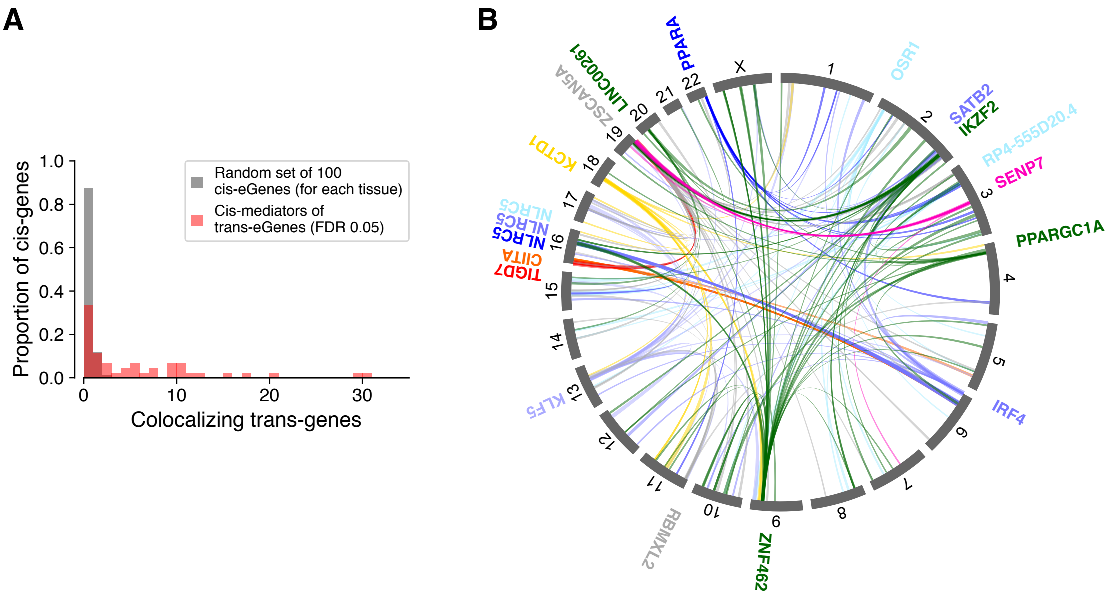

**Fig. S20. Genome-wide colocalizations of *cis*-eQTL mediating *trans*-eQTLs.** A) Number of colocalizations (PP4 $>$ 0.8 and nominal association with eVariant $p < 10^{-5}$) for *cis*-mediators of *trans*-eGenes and for randomly selected *cis*-eQTLs. B) Colocalized *cis*-eQTLs and *trans*-eQTLs (PP4 $>$ 0.8) shown for all *cis*-eQTLs (indicated by gene names) with $\geq$5 colocalizing *trans*-eQTLs (nominal association with discovery *trans*-eVariant $p < 10^{-5}$). The color coding indicates tissues.

##### 11.4.3 Colocalization of *cis*- and *trans*-QTLs

To identify additional *trans*-associations (that did not pass the genome-wide significance threshold for discovery) for the *cis*-regulating genes identified through mediation analysis (Section 11.4.2), we computed colocalization with *coloc* (using default priors). We used the 40 *cis*-eGenes that passed 0.05 FDR in mediation analyses, and computed colocalizations for all inter-chromosomal combinations (for genes with average mappability $>$ 0.8, based on 75-mer mappability with at most two mismatches), using a ±1Mb window around the corresponding eVariant. Candidate *trans*-associations were retained if the posterior probability of colocalization (PP4) was greater than 0.8, and if the nominal association p-value for the discovery *trans*-eVariant was $< 10^{-5}$. To determine the extent to which this approach may produce false positives, we repeated this analyses for a background set of 100 randomly chosen *cis*-eGenes in each tissue. We observed no associations for a majority of background *cis*-eGenes, with at most two associations that passed these thresholds (**fig. S20A**). *Cis*-mediating genes that were identified through mediation analysis but did not colocalize (PP4 $<$ 0.8) with the corresponding *trans*-eGene were excluded. In total, this analysis yielded 248 associations across 15 tissues; the *cis*-eGenes for which $\geq$5 associations were found are summarized in **Fig. 4E** and visualized in (**fig. S20B**).

#### 12 Complex trait associations

##### 12.1 GWAS summary results

To investigate the downstream effects of QTL loci using resources from the GTEx consortium, we first obtained the list of GWAS-significant SNPs from the GWAS catalog [38] (downloaded on 9/7/2018) containing 80,727 entries. Additionally, we selected GWAS of 87 phenotypes covering a broad array of categories including anthropometric, cardio-metabolic, immune, blood, psychiatric, and neurologic traits (**table S4**), and downloaded full sets of summary results from each GWAS consortium or study group; URLs and additional information is available in **table S11**.

We harmonized, lifted genomic coordinates over to hg38, and imputed $z$ scores for all missing variants using our own implementation of BLUP [39, 40] (Best Linear Unbiased Prediction). Imputation was performed within approximately independent LD regions [41], using correlation matrices calculated with GTEx genotype data from European individuals (to reflect the LD structure in the GWAS, predominantly conducted on European populations). This imputation ensured that all common variants in GTEx (i.e. with MAF > 0.01) were represented in the GWAS results. For a detailed description of the summary statistics processing, see [42].

| Category | Phenotype | Abbreviation | Sample Size |
|---|---|---|---|
| Psychiatric-neurologic | CNCR Insomnia all | INSOMN | 113006 |
| Psychiatric-neurologic | IGAP Alzheimer | AD | 54162 |
| Psychiatric-neurologic | Jones et al 2016 Chronotype | CHRONO | 128266 |
| Psychiatric-neurologic | Jones et al 2016 SleepDuration | SLEEP | 128266 |
| Psychiatric-neurologic | PGC ADHD EUR 2017 | ADHD | 53293 |
| Psychiatric-neurologic | pgc.scz2 | SCZ | 150064 |
| Psychiatric-neurologic | SSGAC Depressive Symptoms | DEPR | 180866 |
| Psychiatric-neurologic | SSGAC Education Years Pooled | EDU | 293723 |
| Psychiatric-neurologic | UKB 1160 Sleep duration | SLEEP_UKB | 337119 |
| Psychiatric-neurologic | UKB 1180 Morning or evening person chronotype | CHRONO_UKB | 337119 |
| Psychiatric-neurologic | UKB 1200 Sleeplessness or insomnia | INSOMN_UKB | 337119 |
| Psychiatric-neurologic | UKB 20002 1243 self reported psychological or psychiatric problem | PSY_UKBS | 337119 |
| Psychiatric-neurologic | UKB 20002 1261 self reported multiple sclerosis | MS_UKBS | 337119 |
| Psychiatric-neurologic | UKB 20002 1262 self reported parkinsons disease | PD_UKBS | 337119 |
| Psychiatric-neurologic | UKB 20002 1265 self reported migraine | MIGR_UKBS | 337119 |
| Psychiatric-neurologic | UKB 20002 1289 self reported schizophrenia | SCZ_UKBS | 337119 |
| Psychiatric-neurologic | UKB 20002 1616 self reported insomnia | INSOMN_UKBS | 337119 |
| Psychiatric-neurologic | UKB 20016 Fluid intelligence score | FIS_UKB | 337119 |
| Psychiatric-neurologic | UKB 20127 Neuroticism score | NEUROT_UKB | 337119 |
| Psychiatric-neurologic | UKB G40 Diagnoses main ICD10 G40 Epilepsy | EPI_UKB | 337119 |
| Psychiatric-neurologic | UKB G43 Diagnoses main ICD10 G43 Migraine | MIGR_UKB | 337119 |
| Anthropometric | EGG BW3 EUR | BW | 143677 |
| Anthropometric | ENIGMA Intracraneal Volume | ICV | 30717 |
| Anthropometric | GEFOS Forearm | BMD | 49988 |
| Anthropometric | GIANT HEIGHT | HEIGHT | 253288 |
| Anthropometric | UKB 20022 Birth weight | BW_UKB | 337119 |
| Anthropometric | UKB 21001 Body mass index BMI | BMI_UKB | 337119 |
| Anthropometric | UKB 23099 Body fat percentage | FAT_UKB | 337119 |
| Anthropometric | UKB 50 Standing height | HEIGHT_UKB | 337119 |
| Cardiometabolic | CARDIoGRAM C4D CAD ADDITIVE | CAD | 184305 |
| Cardiometabolic | MAGIC FastingGlucose | FG | 46186 |
| Cardiometabolic | MAGIC ln FastingInsulin | INSUL | 38238 |
| Cardiometabolic | MAGNETIC CH2.DB.ratio | CH2 | 24154 |
| Cardiometabolic | MAGNETIC HDL.C | HDLC | 19270 |
| Cardiometabolic | MAGNETIC IDL.TG | IDL | 21559 |
| Cardiometabolic | MAGNETIC LDL.C | LDLC | 13527 |
| Cardiometabolic | UKB 20002 1065 self reported hypertension | HPT_UKBS | 337119 |
| Cardiometabolic | UKB 20002 1094 self reported deep venous thrombosis dvt | DVT_UKBS | 337119 |
| Cardiometabolic | UKB 20002 1223 self reported type 2 diabetes | T2D_UKBS | 337119 |
| Cardiometabolic | UKB 20002 1473 self reported high cholesterol | HC_UKBS | 337119 |
| Cardiometabolic | UKB 6150 1 Vascular or heart problems diagnosed by doctor Heart attack | MI_UKB | 337119 |
| Cardiometabolic | UKB 6152 5 diagnosed by doctor Blood clot in the leg DVT | DVT_UKB | 337119 |
| Cardiometabolic | UKB 6152 7 diagnosed by doctor Blood clot in the lung | PE_UKB | 337119 |
| Blood | Astle et al 2016 Eosinophil counts | EC | 173480 |
| Blood | Astle et al 2016 Granulocyte count | GC | 173480 |
| Blood | Astle et al 2016 High light scatter reticulocyte count | HRET | 173480 |
| Blood | Astle et al 2016 Lymphocyte counts | LC | 173480 |
| Blood | Astle et al 2016 Monocyte count | MC | 173480 |
| Blood | Astle et al 2016 Myeloid white cell count | MWBC | 173480 |
| Blood | Astle et al 2016 Neutrophil count | NC | 173480 |
| Blood | Astle et al 2016 Platelet count | PLT | 173480 |
| Blood | Astle et al 2016 Red blood cell count | RBC | 173480 |
| Blood | Astle et al 2016 Reticulocyte count | RET | 173480 |
| Blood | Astle et al 2016 Sum basophil neutrophil counts | BNC | 173480 |
| Blood | Astle et al 2016 Sum eosinophil basophil counts | EBC | 173480 |
| Blood | Astle et al 2016 Sum neutrophil eosinophil counts | NEC | 173480 |
| Blood | Astle et al 2016 White blood cell count | WBC | 173480 |
| Cancer | BCAC ER negative BreastCancer EUR | ERNBC | 120000 |
| Cancer | BCAC ER positive BreastCancer EUR | ERPBC | 120000 |
| Cancer | BCAC Overall BreastCancer EUR | BC | 120000 |
| Allergy | EAGLE Eczema | ECZ | 116863 |
| Allergy | UKB 20002 1111 self reported asthma | ATH_UKBS | 337119 |
| Allergy | UKB 20002 1452 self reported eczema or dermatitis | ECZ_UKBS | 337119 |
| Immune | IBD.EUR.Crohns Disease | CD | 20833 |
| Immune | IBD.EUR.Inflammatory Bowel Disease | IBD | 34652 |
| Immune | IBD.EUR.Ulcerative Colitis | UC | 27432 |
| Immune | IMMUNOBASE Systemic lupus erythematosus hg19 | SLE | 23210 |
| Immune | RA OKADA TRANS ETHNIC | RA | 80799 |
| Immune | UKB 20002 1222 self reported type 1 diabetes | T1D_UKBS | 337119 |
| Immune | UKB 20002 1313 self reported ankylosing spondylitis | ASP_UKBS | 337119 |
| Immune | UKB 20002 1453 self reported psoriasis | PSO_UKBS | 337119 |
| Immune | UKB 20002 1461 self reported inflammatory bowel disease | IBD_UKBS | 337119 |
| Immune | UKB 20002 1462 self reported crohns disease | CD_UKBS | 337119 |
| Immune | UKB 20002 1463 self reported ulcerative colitis | UC_UKBS | 337119 |
| Immune | UKB 20002 1464 self reported rheumatoid arthritis | RA_UKBS | 337119 |
| Immune | UKB 6152 8 diagnosed by doctor Asthma | ATH_UKB | 337119 |
| Immune | UKB 6152 9 diagnosed by doctor Hayfever allergic rhinitis or eczema | HAY_UKB | 337119 |
| Aging | UKB 1807 Fathers age at death | FAD_UKB | 337119 |
| Aging | UKB 3526 Mothers age at death | MAD_UKB | 337119 |
| Digestive system disease | UKB 20002 1154 self reported irritable bowel syndrome | IBS_UKBS | 337119 |
| Endocrine system disease | UKB 20002 1225 self reported hyperthyroidism or thyrotoxicosis | HYPERTHY_UKBS | 337119 |
| Endocrine system disease | UKB 20002 1226 self reported hypothyroidism or myxoedema | HYPOTHY_UKBS | 337119 |
| Skeletal system disease | UKB 20002 1309 self reported osteoporosis | OST_UKBS | 337119 |
| Skeletal system disease | UKB 20002 1466 self reported gout | GOUT_UKBS | 337119 |
| Morphology | UKB 2395 2 Hair or balding pattern Pattern 2 | BLDP2_UKB | 337119 |
| Morphology | UKB 2395 3 Hair or balding pattern Pattern 3 | BLDP3_UKB | 337119 |
| Morphology | UKB 2395 4 Hair or balding pattern Pattern 4 | BLDP4_UKB | 337119 |

**Table S4. GWAS datasets.**

#### 12.2 Summarizing across phenotypes and tissues

Many of our analyses generate one statistic for each of the 4,263 (87 $\times$ 49) phenotype/tissue pairs. These can have a complex error structure and a wide range of standard errors largely driven by variation in sample size and correlation between tissues. Thus, the usual iid (independent and identically distributed) assumption behind common statistical tests is not appropriate. When summarizing across phenotypes for a given tissue, we assume independence across phenotypes but take into account their different standard errors. When summarizing across phenotype/tissue pairs, we allow both correlation between tissues, and correlation between phenotypes, and correct for different standard errors. More specifically, let $S_{tp}$ be some statistic estimated in phenotype $p$ and tissue $t$ with standard error se$(S_{tp})$.

In order to summarize across phenotypes for a given tissue, for each tissue $t$, we summarize $S_{t1}, \cdots, S_{tP}$ by fitting the the following linear model:

$$S_{tp} = \mu_S^t + \epsilon_{tp} \tag{1}$$

$$\epsilon_{tp} \sim N(0, \text{se}(S_{tp})^2 \times \sigma_t^2) \tag{2}$$

so that we obtain $\hat{\mu}_S^t$ and se$(\hat{\mu}_S^t)$ as the summary of $S_{t1}, \cdots, S_{tP}$ estimates aggregated across traits, which is in essence a weighted average across phenotypes.

In order to summarize across phenotypes and tissues pairs, similarly, we summarize $S_{11}, \cdots, S_{tp}, \cdots, S_{TP}$ by fitting the following linear model:

$$S_{tp} = \mu_S + \mu_S^t + \mu_S^p + \epsilon_{tp} \tag{3}$$

$$\mu_S^t \sim N(0, \sigma_T^2) \tag{4}$$

$$\mu_S^p \sim N(0, \sigma_P^2) \tag{5}$$

$$\epsilon_{tp} \sim N(0, \text{se}(S_{tp})^2 \times \sigma^2), \tag{6}$$

where $\mu_S^t$ is tissue-specific random intercept (this accounts for tissue specific features common across phenotypes) and $\mu_S^p$ is phenotype-specific random intercept (this accounts for phenotype-specific characteristics and thus accounts for the correlation between tissues for a given phenotype). The estimated $\hat{\mu}_S$ and se$(\hat{\mu}_S)$ is the average $S_{tp}$ across all phenotype/tissue pairs accounting for the complex error structure.

Finally, to test whether two statistics measured across all phenotype/tissue pairs have a different mean, we proceed as follows. First, we form the test statistic $T^{tp} := S_{1,tp} - S_{2,tp}$ which, under the null $\mathcal{H}_0 : \mu_{S_1} = \mu_{S_2}$, has $T^{tp} \sim N(0, \text{se}(S_{1,tp})^2 + \text{se}(S_{2,tp})^2)$. Then, we summarize $T^{tp}$ across all phenotype/tissue pairs (by the procedure described in the previous paragraph) where tissue/trait-specific intercepts are introduced to account for complex correlation structure among $T^{tp}$'s. The resulting statistic $T$ follows $T \sim N(0, \text{se}(T))$ under the null.

#### 12.3 Enrichment of complex trait associated variants among e/sVariants

We examined whether eVariants and sVariants (e/sVariants for short) were enriched among GWAS significant variants (either GWAS catalog variants or belonging to the 87 GWAS traits) using multiple non-overlapping lines of evidence.

##### 12.3.1 Overrepresentation of eVariants/sVariants among GWAS catalog variants

We first investigated e/sVariant enrichment in GWAS catalog variants, comparing the proportion of significant *cis*-e/sVariants and *trans*-eVariants (FDR<10%) among GWAS catalog variants to the proportion among all tested variants.

First, we extracted all GWAS catalog variants which were identified as GTEx v8 variants (by matching rsID). These variants were defined as GWAS catalog variants in this analysis. Next, we defined a variant to be *cis*-e/sVariant if it was identified as *cis*-e/sQTL in at least one gene/intron and one tissue. In the same manner, we defined a variant to be *trans*-eVariant if it was identified as *trans*-eQTL in at least one gene in some tissue. We limited this analysis to autosomal chromosomes. The counts of e/sVariants in the GWAS catalog and among all GTEx v8 variants are shown in **table S5**.

| variant | type | tested | signif | fraction | jackknife_est | jackknife_se |
|---|---|---|---|---|---|---|
| *cis*-eVariant | GWAS | 44,137 | 27,819 | 0.63 | 0.63 | 0.011 |
| *cis*-eVariant | All | 10,390,085 | 4,495,884 | 0.43 | 0.43 | 0.009 |
| *cis*-sVariant | GWAS | 44,029 | 16,197 | 0.37 | 0.37 | 0.013 |
| *cis*-sVariant | All | 10,299,184 | 2,027,766 | 0.2 | 0.2 | 0.007 |
| *trans*-eVariant | GWAS | 34,406 | 10 | 0.00029 | 0.00029 | $9 \times 10^{-5}$ |
| *trans*-eVariant | All | 5,154,755 | 215 | $4.2 \times 10^{-5}$ | $4.2 \times 10^{-5}$ | $3.6 \times 10^{-6}$ |

**Table S5. GWAS catalog variant overlap with e/sVariants.** The number of tested variants is shown under the column named **tested**. The jackknife estimates of fraction for e/sVariants are shown in the **jackknife_est** and **jackknife_se** columns.

To compute the standard error of the observed fraction/fold-enrichment (defined below) of e/sVariant in GWAS catalog, we accounted for the fact that variants included in this analysis were correlated with each other due to LD. So, we performed block jackknife to obtain the standard error of the fraction/fold-enrichment estimate. Specifically, we ordered the list of variants by the genomic position and divided them into 200 consecutive blocks. Then, we calculated the statistic of interest by removing the variants in $i$th block. This procedure gave rise to 200 delete-one statistics, $\hat{\theta}_{(1)}, \cdots, \hat{\theta}_{(200)}$. Then, we calculated the jackknife estimate and standard error as follows:

$$\hat{\theta}_{(\cdot)} = \frac{1}{K}\sum_{i=1}^{K}\hat{\theta}_{(i)} \tag{7}$$

$$\hat{\theta}_{jack} = K\hat{\theta} - (K-1)\hat{\theta}_{(\cdot)} \tag{8}$$

$$\text{Var}(\hat{\theta}_{jack}) = \frac{K-1}{K}\sum_{i=1}^{K}(\hat{\theta}_{(\cdot)} - \hat{\theta}_{(i)})^2, \tag{9}$$

where $K$ was the number of blocks, which was set to 200 in this analysis. Specifically, we were interested in the cases for measuring the fraction of e/sVariants among a list of variants and/or the enrichment fold of e/sVariants among GWAS catalog variants versus the baseline. The statistics for each case is specified below:

- Fraction for variant list A:

$$\hat{f}_{\text{A}} = \frac{\text{number of variants in A that are e/sVariant}}{\text{number of variants in A}}$$

- Enrichment of GWAS catalog variants against all variants tested:

$$\frac{\hat{f}_{\text{GWAS}}}{\hat{f}_{\text{All}}}$$

The jackknife estimates of enrichment are shown in **Fig. 5A** and **table S6**. Moreover, the jackknife estimate of the fraction for e/sVariants among GWAS catalog variants is shown and compared with the fraction among all tested variants in **fig. S21** (with numbers reported in **table S5**).

To rule out the possibility that the enrichment was driven by shared functional features of e/sVariants and GWAS variants we corrected for them using QTLEnrich and stratified LDSC regression approaches as described below. Due to the sparsity of *trans*-QTL data, these approaches were applied to *cis*-QTLs only.

| variant | enrichment | jackknife_est | jackknife_se |
|---|---|---|---|
| *cis*-eVariant | 1.46 | 1.46 | 0.021 |
| *cis*-sVariant | 1.87 | 1.87 | 0.0627 |
| *trans*-eVariant | 6.97 | 6.95 | 2.12 |

**Table S6. GWAS catalog enrichment among e/sVariants.**

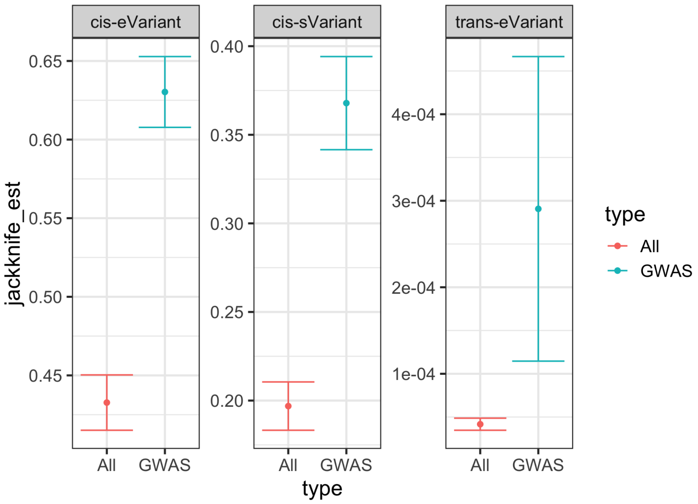

**Fig. S21. Enrichment of GWAS catalog variants among e/sVariants.** We compared the fraction of *cis*-e/sVariant, and *trans*-eVariant among all GTEx variants being tested (labelled as `All`) and among GWAS catalog variants which overlapped with GTEx variants (labelled as `GWAS`). Here fraction estimate and 95% confidence interval based on jackkife are shown.

##### 12.3.2 QTLEnrich: overrepresentation of complex trait associations among *cis*-QTLs

To test whether *cis*-e/sVariants in each of the 49 tissues were enriched for genetic associations with various complex diseases and traits, compared to expectation, we applied an updated version of QTLEnrich (v2) [43] ( `https://github.com/segrelabgenomics/QTLEnrich`) that accounts for increase in number of genes with e/sVariants discovered in GTEx v8 compared to v6p. QTLEnrich.v2 assesses the enrichment of top ranked trait associations (GWAS p-value<0.05 used here) amongst a set of significant *cis*-e/sVariants in a given tissue, accounting for potential confounding factors.

Briefly, an enrichment p-value was computed for each tissue/phenotype pair tested, as the fraction of 1,000 to 100,000 randomly sampled sets of null variants (of equal size to that of the e/sVariant set), matched on three potential confounding factors relative to the significant e/sVariants, whose fold-enrichment of top ranked phenotype associations is equal to or larger than that of the significant set of e/sVariants. An adaptive permutation scheme was used. The confounding factors included: (i) minor allele frequency that was computed per tissue, (ii) distance of each e/sVariant variant to the transcription start site (TSS) of the nearest expressed gene in a given tissue, and (iii) number of LD proxy variants ($r^2 \geq 0.5$), computed using the European subset of GTEx WGS samples. The null variant sets were randomly sampled from all variants within $\pm$1Mb around the TSS of all genes expressed in the tested tissue, excluding variants that were significant eVariant or sVariant in any of the 49 tissues analyzed in v8. Sampling was applied to decile bins of the confounding factors with replacement. Fold-enrichment was computed as the number of e/sVariants or null variants with a GWAS p-values below 0.05 divided by 5% of the variant set size. An adjusted fold-enrichment was computed for each *cis*-e/sVariant set, as the fold-enrichment of the *cis*-e/sVariant set divided by the median fold-enrichment of 1,000 randomly sampled sets of confounder-matched null variants. The most significant *cis* eVariant or sVariant per eGene or sGene, respectively (at FDR<0.05) was used to reduce potential inflation of enrichment due to local LD. QTLEnrich.v2 was applied to 87 GWAS by 49 tissues (4,263 trait-tissue pairs). Bonferroni correction was used to determine significant tissue/phenotype pairs ($P < 1.17 \times 10^{-5}$), and the adjusted fold-enrichment was used as the test statistic to rank significant tissues based on their enrichment, as it corrects for enrichment of trait associations amongst matched null variants.

##### 12.3.3 Stratified LDSC regression-based enrichment

Finally, we examined the enrichment of heritability through stratified LD-score regression (S-LDSC) in *cis*-e/sVariant relative to the rest of the genome, in order to corroborate the other enrichment estimates and to analyze how incorporation fine-mapping information affects the enrichments.

We applied three e/sVariant annotations as proposed in [44]. In particular, for each tissue, the *cis*-e/sVariant annotation was defined as the set of *cis*-e/sVariants identified in that tissue. The fine-mapped *cis*-e/sVariant annotation was defined as the set of variants within the 95% credible set in DAPG [45] with posterior inclusion probability (PIP) greater than 0.01 in eGene or sGene. Moreover, we defined the continuous annotation, MaxCPP, as described in [44], based on DAPG PIP. Specifically, DAPG MaxCPP was defined as maximum PIP of the variant over all DAPG 95% credible sets for eGene and sGene respectively in each tissue. All annotations were lifted over to hg19 for this analysis. For each tissue, S-LDSC was calculated using 1000G phase 3 genotypes shared in `https://data.broadinstitute.org/alkesgroup/LDSCORE/1000G_Phase3_plinkfiles.tgz`

(without MAF filter). S-LDSC regression for each tissue was performed within the list of HapMap3 SNPs shared in https://data.broadinstitute.org/alkesgroup/LDSCORE/w_hm3.snplist.bz2 with regression weights shared in https://data.broadinstitute.org/alkesgroup/LDSCORE/1000G_Phase3_weights_hm3_no_MHC.tgz. We analyzed all 87 traits in all 49 tissues. For each trait/tissue pair, S-LDSC regression was performed with baselineLD model (version 1.1) [46] along with either e/sVariant annotation or finemapped e/sVariant annotation or DAPG MaxCPP. Furthermore, we removed trait/tissue pairs with heritability $\hat{h}_g^2$ not significantly greater than 0 (at $\alpha = 0.05$) in subsequent analysis. For each tissue, we calculated an average enrichment signal across all traits by the procedure described in Section 12.2 and the results are shown in **figure S22**.

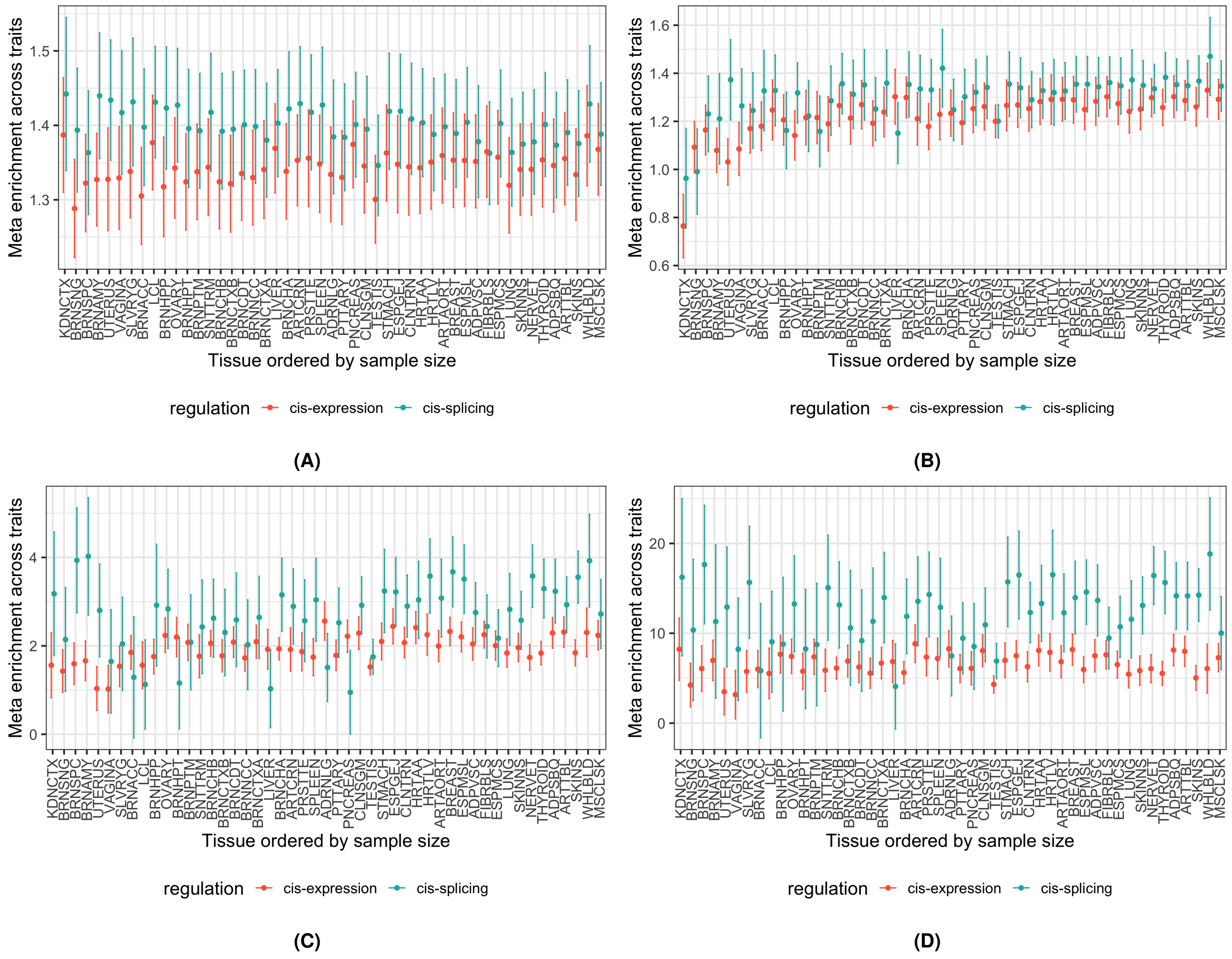

**Fig. S22. Enrichment of GWAS associations across tissues.** Enrichment estimates by tissue are summarized across traits (on y-axis) with error bars representing 95% confidence intervals. Tissues are listed on the x-axis, ordered by sample size with abbreviation described in **table S8**. *Cis*-expression results are shown in red and *cis*-splicing results are shown in green. A) QTLEnrich; B) S-LDSC on e/sVariant annotation; C) S-LDSC on finemapped QTL annotation (DAPG); D) S-LDSC on MaxCPP annotation (DAPG).

##### 12.3.4 Summary statistics of enrichment analysis

The fold enrichment estimates calculated in Section 12.3.2 and Section 12.3.3 were meta-analyzed across tissues and traits by method as described in Section 12.2. The resulting statistics are listed in **table S7** and **Fig. 5A**.

| regulation | enrichment | enrichment_se | method |
|---|---|---|---|
| *cis*-expression | 1.43 | 0.0382 | QTLEnrich. |
| *cis*-splicing | 1.52 | 0.0447 | QTLEnrich. |
| *cis*-expression | 1.44 | 0.0502 | S-LDSC.QTL |
| *cis*-splicing | 1.56 | 0.0655 | S-LDSC.QTL |
| *cis*-expression | 2.54 | 0.199 | S-LDSC.Finemapped |
| *cis*-splicing | 3.1 | 0.354 | S-LDSC.Finemapped |
| *cis*-expression | 11.1 | 1.21 | S-LDSC.MaxCPP |
| *cis*-splicing | 14.2 | 2.41 | S-LDSC.MaxCPP |

**Table S7. Enrichment of GWAS signal among *cis*-e/sVariants.**

#### 12.4 Calculating the joint contribution of e/sVariants to phenotype heritability

The previous analyses showed that when analyzed separately, both *cis*-eQTLs and *cis*-sQTLs contribute to GWAS. Next, we wanted to analyze these in a joint model to evaluate their independent relative contributions to GWAS heritability, using S-LDSC regression. The annotation-specific regression coefficient $\tau_c$ in S-LDSC measures the contribution of each annotation class $c$ (e.g., *cis*-eVariant or -sVariant) to the heritability of the complex trait in joint analysis — i.e., after accounting for the contribution of the other classes. Since the magnitude of $\tau_c$ also depends on the total heritability, here we scaled $\tau_c$ by chip heritability. Specifically, we defined $\mathcal{R}^{\text{ldsc}}$ as the proportion of chip heritability that could be attributed to eVariant or sVariant annotation. Namely, for $c$th binary annotation in $j$th tissue and $k$th trait,

$$\mathcal{R}^{\text{ldsc}}_{c,jk} = \frac{\tau_{c,jk} p_{c,j}}{h^2_{jk}/M} \tag{10}$$

$$\text{se}(\mathcal{R}^{\text{ldsc}}_{c,jk}) = \frac{\text{se}(\tau_{c,jk}) p_{c,j}}{h^2_{jk}/M}, \tag{11}$$

where $h^2_{jk}/M$ was the per-SNP heritability estimated by S-LDSC, $p_{c,j}$ was the proportion of SNPs lying in the annotation $c$ for tissue $j$ and $\tau_{c,jk}$ was the coefficient estimate of $c$th annotation in the corresponding trait/tissue pair.

In order to examine the independent contribution of *cis*-e/sVariants to heritability, we used only eVariant, sVariant, and the 'base' annotation including all SNPs to run S-LDSC. The rationale was that other functional annotations (genomic and epigenomic annotations) were highly correlated with e/sVariant annotation and we were not interested in partitioning heritability among these annotations. For each tissue/trait pair, we performed S-LDSC regression using *cis*-e/sVariant annotation (as described in Section 12.3.3). We computed $\mathcal{R}^{\text{ldsc}}$ for e/sVariant annotations in each trait/tissue pair. Trait/tissue pairs with $\hat{h}^2_g$ not significantly greater than 0 (at $\alpha = 0.05$) were excluded. $\mathcal{R}^{\text{ldsc}}$ estimates were aggregated across traits using the procedure described in Section 12.2. The resulting aggregated $\mathcal{R}^{\text{ldsc}}$ along with 95% confidence interval are shown in **fig. S23**.

#### 12.5 Causal gene prioritization

The goal here is to identify and prioritize candidate causal genes affecting a complex trait through transcriptome regulation in *cis*. We employed two classes of methods to identify the target genes of GWAS loci. One is based on the colocalization of GWAS and *cis*-QTL loci - i.e., studying whether the causal variant for the phenotype is the same as the causal variant for the molecular trait. The other class is based on the association between the genetically regulated component of gene expression (or splicing) with the phenotype.

##### 12.5.1 *cis*-QTL-GWAS colocalization: ENLOC

To identify target genes and GWAS loci of interest we performed *cis* colocalization analysis on 87 GWAS traits across 49 tissues. After extensive comparison of colocalization methods we chose ENLOC [18] as our primary approach. The main factors leading to this decision were the need to account for allelic heterogeneity in expression/splicing traits (**Fig. 2, fig. S8**), and the high sensitivity of the methods to the prior on the enrichment of GWAS probability of causality given the *cis*-e/sQTL's posterior inclusion probability (PIP). See details in [42]. In short, ENLOC yields regional colocalization probabilities (rcp) for (GWAS region, trait, tissue, gene) or (GWAS region, trait, tissue, intron) tuples.

First, for protein-coding and lincRNA genes in the *cis*-eQTL analysis, the expression levels and genotypes in the *cis*-window were processed with DAPG [45]. We used individuals of European ancestry in the GTEx study and variants with MAF $> 0.01$,

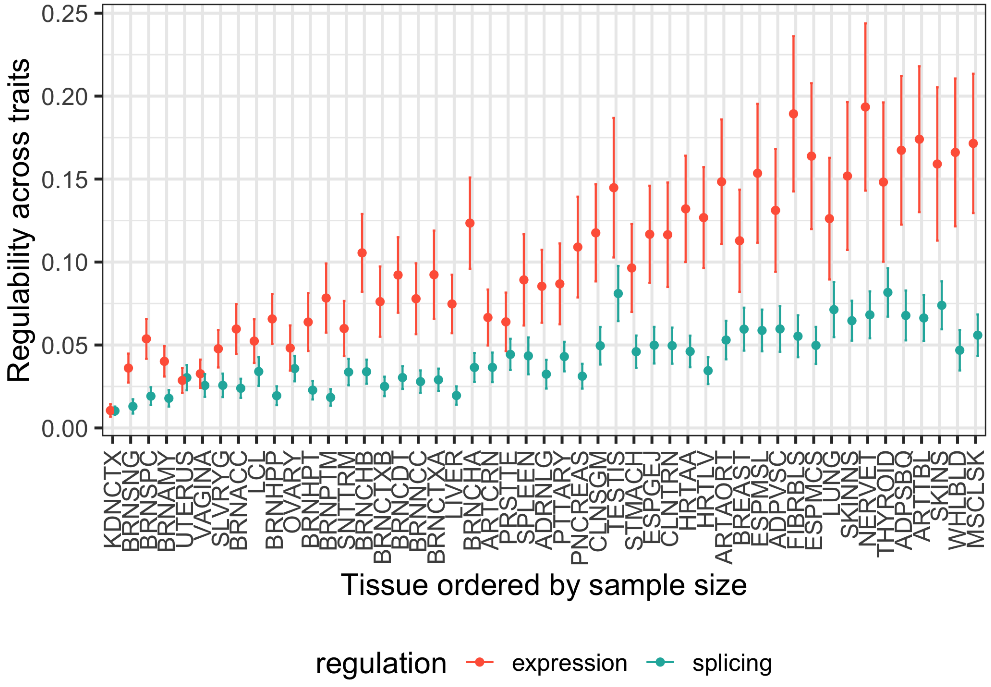

**Fig. S23. Independent GWAS contribution of *cis*-eQTLs and sQTLs** The partitioned contributions of *cis*-eVariant (red) and *cis*-sVariant (green) to complex trait heritability were measured by $\mathcal{R}^{\text{ldsc}}$ in S-LDSC regression. Here, for each tissue (on x-axis ordered by sample size with abbreviation described in **table S8**), the meta-analyzed $\mathcal{R}^{\text{ldsc}}$ (across traits) are shown on y-axis, where error bar indicates the 95% confidence interval.

to generate eVariant enrichment estimates and variants' posterior inclusion probabilities (PIP). The main motivation in using individuals of European ancestry was to match the LD structure as close as possible to the ones from the GWAS studies (mostly European-based). Expression levels were corrected by the same covariates as in the main eVariant analysis.

Second, the imputed GWAS summary statistics were split into approximately LD-independent regions [41], each region defining a GWAS locus. Lastly, ENLOC was performed for all *cis*-eVariant regions and overlapping GWAS loci for each trait, yielding colocalization results for 12,662,634 $(\text{tissue}, \text{gene}, \text{GWAS locus}, \text{trait})$ tuples.

We found good agreement between ENLOC results based on all individuals compared to the results using Europeans only as shown in **figure S24-A)**; when using the best rcp value across tissues for a gene, most colocalized genes can be detected through both approaches.

For each trait, we counted how many GWAS loci contained a GWAS significant hit, and also contained a gene with ENLOC colocalization $rcp > 0.5$. As shown in **fig. S25C**, across traits, a median 21% of loci with a GWAS signal contain an ENLOC colocalized signal. Given ENLOC's conservative nature, we caution that $rcp < 0.5$ does not mean that there is no causal relationship between the molecular trait and the complex trait; rather, it should be interpreted as a lack of sufficient evidence with current data.

We repeated this process for splicing events' excision ratios from Leafcutter [13], for all 87 traits and 49 tissues; we did not correct for covariates in this case. There were a few key differences with the eQTL data: the ratios within a cluster were correlated (since they added up to 1), and the *cis*-sQTL summary statistics contained a significantly larger number of splicing events than genes (4,337,796 intron-tissue pairs versus 1,207,976 gene-tissue pairs). We obtained analogous RCP and PIP values for 82,301,739 $(\text{tissue}, \text{intron}, \text{GWAS locus}, \text{trait})$ tuples. We summarized the findings in **fig. S26**. We observed a smaller proportion of GWAS loci containing a colocalized intron, (median 11% across traits).

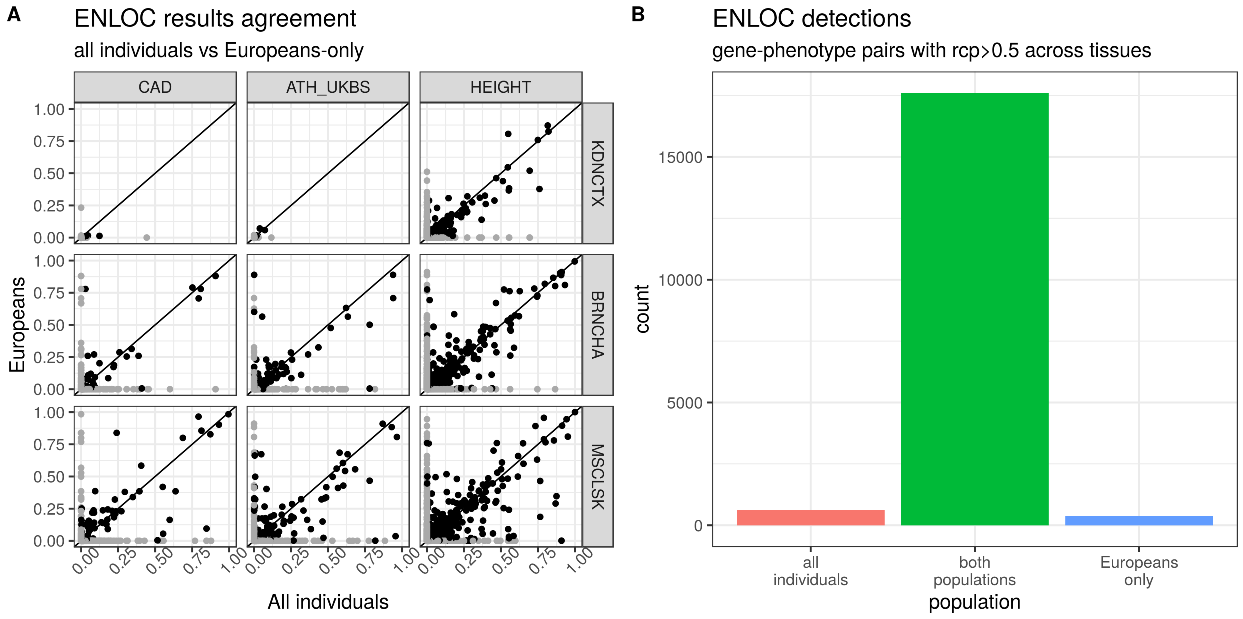

**Fig. S24. ENLOC on different populations.** A) ENLOC regional colocalization probabilities (rcp) for genes at representative trait-tissue combinations (CAD: Coronary Artery Disease, ATH_UKBS: Athma in UK Biobank, HEIGHT: Height in GIANT Consortium; KDNCTX: Kidney - Cortex, BRNCHA: Brain - Cerebellum, MSCLSK: Muscle - Skeletal). We observe a general agreement for genes achieving rcp$> 0.5$. B) Number of gene-phenotype pairs achieving rcp $> 0.5$ in at least one tissue. Most such pairs are detected by both methods (17598), with only a few detected using all individuals (623) or European individuals (378).

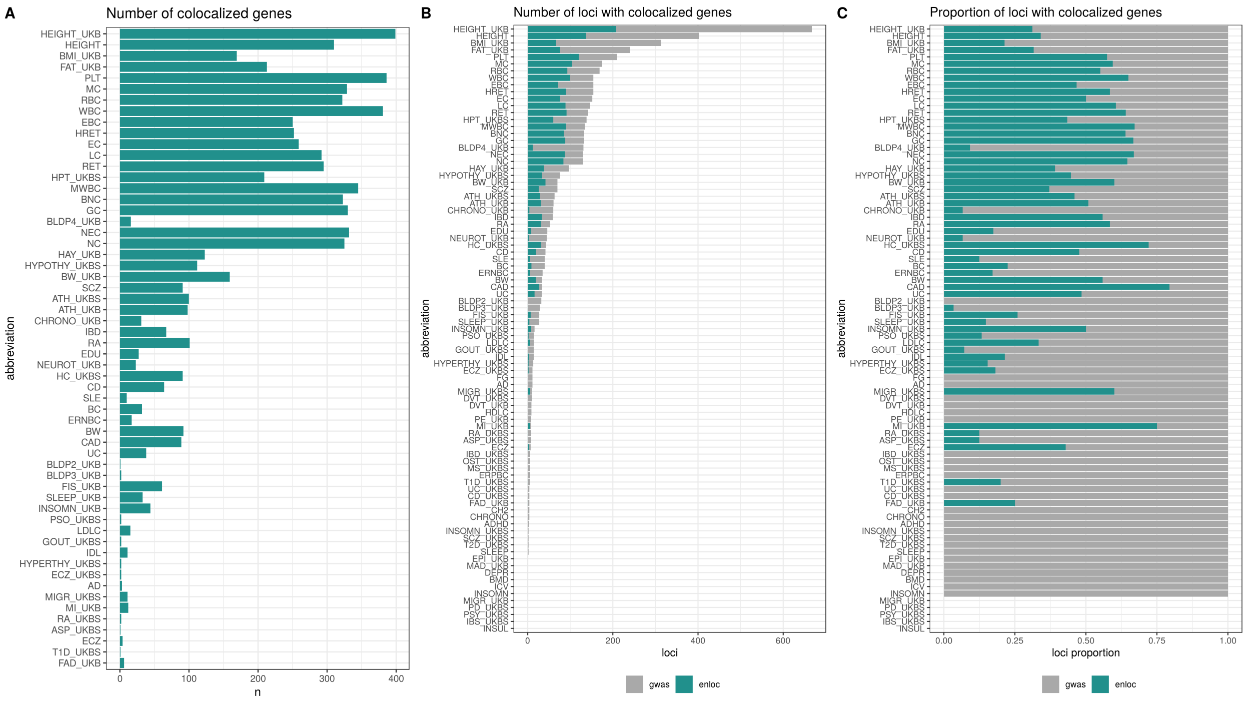

**Fig. S25. GWAS colocalization with *cis*-eQTLs** by ENLOC for each of the 87 GWAS traits aggregated across the 49 tissues. The traits are ordered by the number of GWAS-significant loci (approximately independent LD regions from [41]). A) Number of colocalized genes achieving ENLOC $rcp > 0.5$ in at least one tissue, for each GWAS trait. The number of colocalized results tends to increase with the number of GWAS-significant loci. B) Number of loci with at least one GWAS-significant variant (dark gray), and among them those with at least one gene reaching $rcp > 0.5$ (dark green). The bottom five traits have no loci with GWAS-significant associations. C) Proportion of loci with at least one GWAS-significant hit that contain at least one colocalized gene. Across traits, a median of 21% of the GWAS loci contain colocalized results. See phenotype abbreviation list in **table S4**.

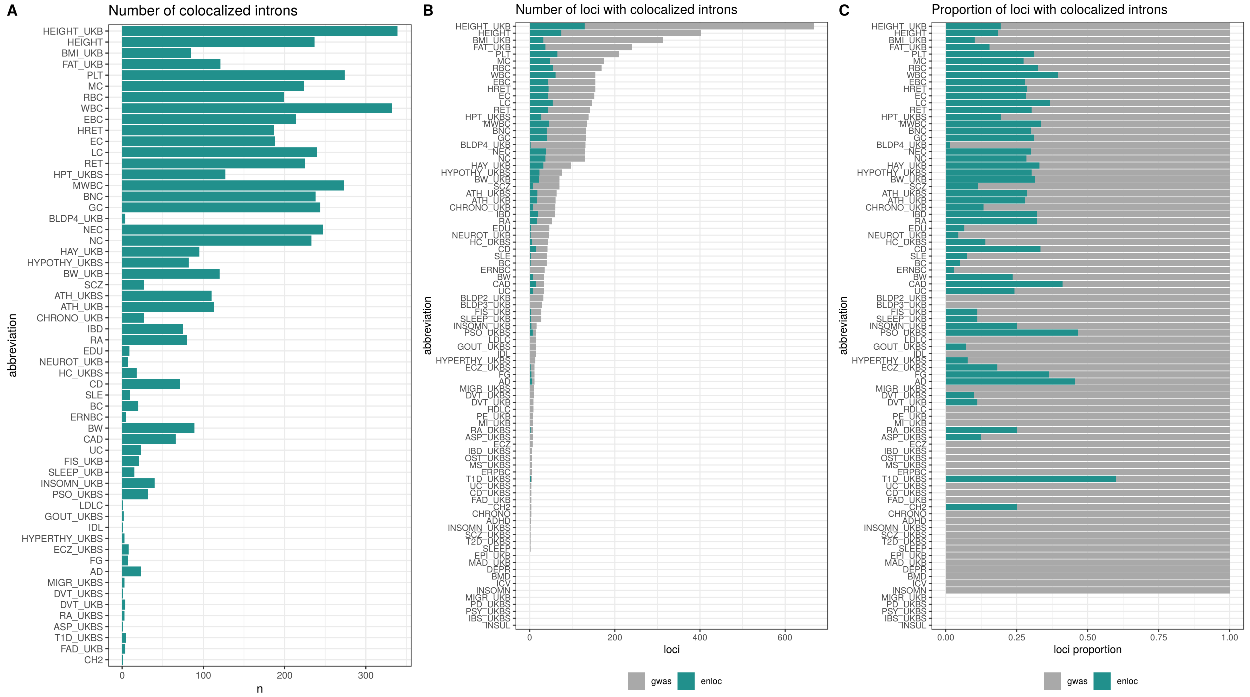

**Fig. S26. GWAS colocalization with *cis*-sQTLs** by ENLOC for each of the 87 GWAS traits aggregated across the 49 tissues. The traits are ordered by the number of GWAS-significant loci (approximately independent LD regions from [41]). A) Number of colocalized introns, achieving ENLOC $rcp > 0.5$ in at least one tissue, for each GWAS trait. As with gene expression results, the number of colocalized results tends to increase with the number of GWAS-significant loci. B) Number of loci with at least one GWAS-significant variant (dark gray), and among them those with one intron achieving $rcp > 0.5$ (dark green). The bottom five traits have no loci with GWAS-significant associations. C) Proportion of loci with at least one GWAS-significant hit loci with at least one colocalized intron. Across traits, a median of 11% of the GWAS loci contain a colocalized result, lower than the gene expression counterpart (21%), indicating a decreased power in the sQTL study. See phenotype abbreviation list in **table S4**.

###### 12.5.2 Trait association with predicted expression and splicing in *cis*: PrediXcan

To identify target genes and GWAS loci of interest with the association approach, we trained prediction models for expression and splicing in 49 tissues, performed S-PrediXcan [47] analysis and aggregated the evidence across all tissues with S-MultiXcan [48].

We generated prediction models following methods described in [49] and [47]. The analysis was restricted to genes that were annotated as protein coding or lincRNA. For each gene/tissue pair, we trained models via the Elastic Net algorithm [50], for expression levels and genotypes in individuals of European ancestry. We used variants in the eQTL *cis*-window with MAF $> 0.01$ belonging to the HapMap [51] CEU track. Expression levels were corrected by the same covariates as in the main eQTL analysis. We employed a cross-validated strategy, and kept only models that achieved cross-validated correlation $\rho > 0.1$ and cross-validated prediction performance p-value $p < 0.05$. For each gene/tissue pair, we compiled the covariance matrix of variants present in the model, to be used as an LD reference panel for GWAS summary statistics. For every gene, we also computed the covariance of all the variants present in the different tissue models, compiling a cross-tissue LD panel to be used with S-MultiXcan. Analogous models were built for intron splicing ratios instead of expression, using a window from 1Mb upstream of the intron start to 1Mb downstream of the intron end, as well as matching LD reference panels. **Table S8** summarizes the number of models generated, for a total of 281,902 gene-tissue pairs and 525,823 intron-tissue pairs.

| name | abbreviation | european samples | expression models | splicing models |
|---|---|---|---|---|
| Adipose - Subcutaneous | ADPSBQ | 491 | 8606 | 15068 |
| Adipose - Visceral (Omentum) | ADPVSC | 401 | 7336 | 13765 |
| Adrenal Gland | ADRNLG | 200 | 4866 | 9073 |
| Artery - Aorta | ARTAORT | 338 | 7619 | 12588 |
| Artery - Coronary | ARTCRN | 180 | 4000 | 9264 |
| Artery - Tibial | ARTTBL | 489 | 8657 | 14364 |
| Brain - Amygdala | BRNAMY | 119 | 2772 | 5549 |
| Brain - Anterior cingulate cortex (BA24) | BRNACC | 135 | 3568 | 6758 |
| Brain - Caudate (basal ganglia) | BRNCDT | 172 | 5026 | 7978 |
| Brain - Cerebellar Hemisphere | BRNCHB | 157 | 5818 | 11105 |
| Brain - Cerebellum | BRNCHA | 188 | 6824 | 12051 |
| Brain - Cortex | BRNCTXA | 184 | 5447 | 8827 |
| Brain - Frontal Cortex (BA9) | BRNCTXB | 158 | 4580 | 7680 |
| Brain - Hippocampus | BRNHPP | 150 | 3723 | 6354 |
| Brain - Hypothalamus | BRNHPT | 157 | 3659 | 6932 |
| Brain - Nucleus accumbens (basal ganglia) | BRNNCC | 181 | 4909 | 8204 |
| Brain - Putamen (basal ganglia) | BRNPTM | 153 | 4419 | 6571 |
| Brain - Spinal cord (cervical c-1) | BRNSPC | 115 | 3229 | 6690 |
| Brain - Substantia nigra | BRNSNG | 101 | 2542 | 5167 |
| Breast - Mammary Tissue | BREAST | 337 | 6460 | 13827 |
| Cells - Cultured fibroblasts | FIBRBLS | 300 | 8887 | 14004 |
| Cells - EBV-transformed lymphocytes | LCL | 116 | 2940 | 9375 |
| Colon - Sigmoid | CLNSGM | 274 | 6145 | 11677 |
| Colon - Transverse | CLNTRN | 306 | 6295 | 11534 |
| Esophagus - Gastroesophageal Junction | ESPGEJ | 281 | 6346 | 11632 |
| Esophagus - Mucosa | ESPMCS | 423 | 8513 | 12472 |
| Esophagus - Muscularis | ESPMSL | 399 | 8242 | 13247 |
| Heart - Atrial Appendage | HRTAA | 322 | 6653 | 10704 |
| Heart - Left Ventricle | HRTLV | 334 | 5997 | 8327 |
| Kidney - Cortex | KDNCTX | 65 | 1635 | 4577 |
| Liver | LIVER | 183 | 3810 | 6478 |
| Lung | LUNG | 444 | 7925 | 15494 |
| Minor Salivary Gland | SLVRYG | 119 | 2954 | 8050 |
| Muscle - Skeletal | MSCLSK | 602 | 7618 | 11625 |
| Nerve - Tibial | NERVET | 449 | 10006 | 16809 |
| Ovary | OVARY | 140 | 3564 | 9139 |
| Pancreas | PNCREAS | 253 | 5923 | 7778 |
| Pituitary | PTTARY | 219 | 5711 | 12119 |
| Prostate | PRSTTE | 186 | 4298 | 10077 |
| Skin - Not Sun Exposed (Suprapubic) | SKINNS | 440 | 8597 | 14756 |
| Skin - Sun Exposed (Lower leg) | SKINS | 517 | 9265 | 15440 |
| Small Intestine - Terminal Ileum | SNTTRM | 144 | 3633 | 8875 |
| Spleen | SPLEEN | 186 | 5805 | 10303 |
| Stomach | STMACH | 269 | 5149 | 9179 |
| Testis | TESTIS | 277 | 9941 | 32916 |
| Thyroid | THYROID | 494 | 9665 | 16855 |
| Uterus | UTERUS | 108 | 2573 | 7909 |
| Vagina | VAGINA | 122 | 2512 | 7933 |
| Whole Blood | WHLBLD | 573 | 7240 | 8724 |
| total | | | 281902 | 525823 |

**Table S8. Summary of prediction models** for each tissue, for expression levels and splicing ratios.

We executed S-PrediXcan [47] on the 87 traits for all gene expression models to obtain (gene,tissue,trait) associations, which were aggregated across tissues with MultiXcan. This process was repeated for intron models. Since we had GWAS summary results available, we used the summary versions of the methods: S-PrediXcan and S-MultiXcan. In [48] we have shown that MultiXcan integrates predictions from multiple-tissues simultaneously while accounting for correlation across tissues, achieving higher power than single-tissue approaches like S-PrediXcan.

Finally, we counted the GWAS loci explained by PrediXcan as follows. For each trait, we split the GWAS study in approximately independent LD regions [41] and counted which of these loci contained a GWAS significant hit ($b < 0.05/$n_variants $\sim 5 \times 10^{-9}$). Among them, we counted which contained a gene achieving S-MultiXcan significance (Bonferroni threshold $b < 0.05/$n_genes $\approx 2 \times 10^{-6}$), and which of these also contained evidence of colocalization via ENLOC $rcp > 0.5$. As shown in **fig. S27C**, around 67% of loci with a GWAS signal contain an S-MultiXcan significant signal, and about 20% is also colocalized via ENLOC. For splicing models, we summarize the results in **fig. S28**; panel C shows that the proportion of GWAS loci containing an S-MultiXcan association is also about 60% similar to expression but only about 11% is colocalized.

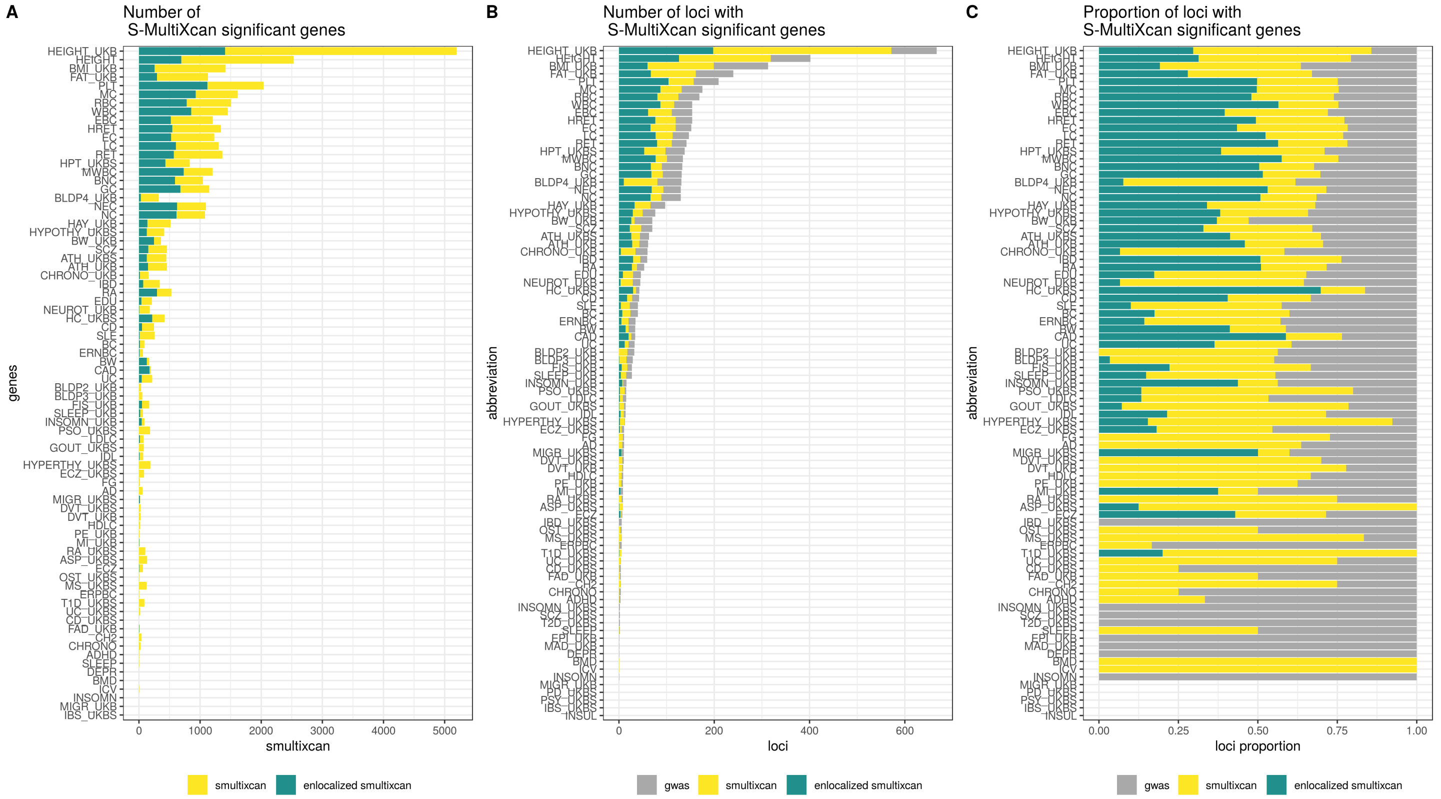

**Fig. S27. Expression associations by S-MultiXcan.** Summary of S-MultiXcan detections for each of the 87 traits using gene expression models. The traits are ordered by the number of GWAS-significant variants. A) Number of S-MultiXcan significant genes (yellow) and subset also achieving ENLOC $rcp > 0.5$ in any tissue (dark green). B) Number of loci (approximately independent LD regions [41]) with a significant GWAS association (gray), a significant S-MultiXcan association (yellow), and a significant S-MultiXcan association that is colocalized (dark green). Anthropometric and Blood traits tend to present the largest number of associated loci, with Height from two independent studies leading the number of associations. C) Proportions of loci with significant GWAS associations (gray) that contain S-MultiXcan (yellow) and colocalized S-MultiXcan associations (dark green). S-MultiXcan has a high power for detecting associations: across traits, a median of 67% of GWAS-associated loci show a S-MultiXcan detection, while 17% show a colocalized S-MultiXcan detection. See phenotype abbreviation list in **table S4**.

#### 12.6 Concordance of downstream phenotype effects of multiple variants affecting the same gene

Many genes have multiple independent *cis*-eQTLs, independent *cis*-eQTLs and sQTLs, as well as both common QTLs and rare coding variants. This creates an opportunity to test if these different variants have concordant effects on downstream GWAS phenotypes when the gene appears to contribute to traits.

##### 12.6.1 Concordance of downstream effects on phenotype between independent *cis*-eQTLs

We first asked whether the downstream GWAS effects of primary eQTLs differed from secondary eQTLs in *cis*. We defined downstream gene-level effect size of independent *cis*-eVariants on phenotypes as the ratio of the GWAS and the *cis*-eQTL effect sizes $\beta_{\text{gene}} = \delta_{\text{GWAS}}/\gamma_{\text{QTL}}$. To compare them, we calculated the correlation between the downstream effect sizes of secondary vs primary *cis*-eQTLs.

To make sure that the correlation was not driven by any residual LD between independent *cis*-eQTL signals, we used the correlation among genes not contributing to GWAS as the null. We would not expect concordance between primary and secondary *cis*-eQTLs of these non-causal genes, and thus any correlation would be attributable to LD between the variants. We used the colocalization probability as a proxy to select the putatively causal and non-causal null gene sets.

**Fig. 5C** shows the median correlation between primary and secondary eQTLs for the 87 phenotypes for subsets with increasing colocalization probability threshold. As the threshold for colocalization increases, enriching for causal genes, the correlation increases, and the difference relative to low colocalization probability also increases. These observations suggest that the mediated effect of independent *cis*-eVariants are concordant.

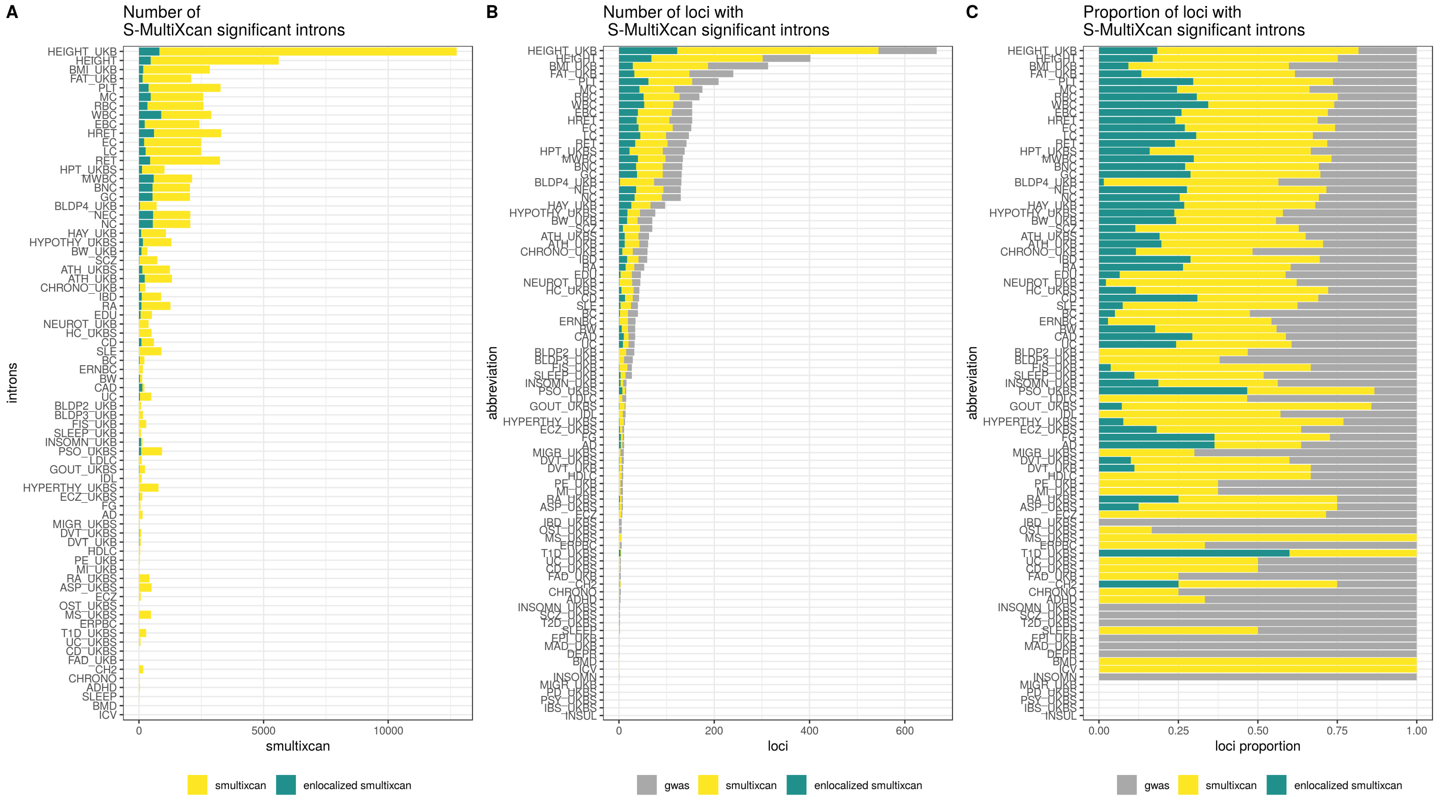

**Fig. S28. Splicing associations by S-MultiXcan.** Summary of S-MultiXcan detections for each of the 87 traits using splicing models. The traits are ordered by the number of GWAS-significant variants. A) Number of S-MultiXcan significant introns (yellow) and subset also achieving ENLOC $rcp > 0.5$ in any tissue (dark green). B) Number of loci (approximately independent LD regions [41]) with a significant GWAS association (gray), a significant S-MultiXcan association (yellow), and a significant S-MultiXcan association that is colocalized (dark green). As in the case of expression models, Anthropometric and Blood traits tend to present the largest number of associated loci. C) Proportion of loci with significant GWAS associations (gray) that contain S-MultiXcan (yellow) and colocalized S-MultiXcan associations (dark green). Across traits, a median of 63% of GWAS-associated loci show an S-MultiXcan detection, while 11% show a colocalized S-MultiXcan detection. These proportions are lower than those for expression (67% and 17% respectively). See phenotype abbreviation list in **table S4.**

##### 12.6.2 Concordance of downstream effects of independent eQTLs and sQTLs in *cis*

Next, we investigated whether the GWAS effects for non-shared *cis*-eQTLs and *cis*-sQTLs affecting the same target e/sGene are concordant. Since the direction of sQTL effects is not directly comparable to eQTL effects, the effect size concordance approach above was not applicable. Thus, we analyzed this by colocalization concordance. We restricted our analyses to genes that have independent *cis*-eQTLs and *cis*-sQTLs, i.e., max $r^2$ between independent eQTLs and sQTLs for that gene $<$ 0.2. Across the approximately LD-independent GWAS regions, we estimated the proportion of *cis*-eQTLs and *cis*-sQTLs that are colocalized in at least one tissue, respectively. Then for colocalized *cis*-sQTLs, we estimated again the proportion of colocalized *cis*-eQTLs, and vice versa, to determine if one type of the molecular *cis*-QTL colocalizing increases the chance of the other molecular *cis*-QTL also colocalizing in the same locus (**fig. S29**). We used Fisher's exact test to estimate the significance of the association.

##### 12.6.3 Concordance of GWAS effects of rare variants and *cis*-eQTLs

Finally, we investigated whether the GWAS effects of rare variants and *cis*-eQTLs are concordant. We used the results of a genome-wide rare variant association analysis based on 50K exomes from UK Biobank from [52], and colocalized GTEx *cis*-eQTLs. Out of 3,166 phenotypes analyzed in [52], 50 were included in the list of phenotypes studied by GTEx (37 UKB traits and 13 other traits where the trait of interest was the same). We counted if a rare variant associated gene is colocalized with the given trait in at least one tissue in GTEx, using separately rare variant gene-based associations with the non-benign coding variants (coding model) and loss of function (LoF) variants (LoF model). We summarized the results for three rare variant significance categories - $P \geq 10^{-3}, P \in (10^{-3}, 10^{-6}], P < 10^{-6}$ (**Fig. 5D**). To estimate if there is an association between the rare variant significance category and the count of colocalized gene-trait pairs, we used Pearson's chi-squared test with Monte Carlo simulations (B = 10,000 replicate samples).

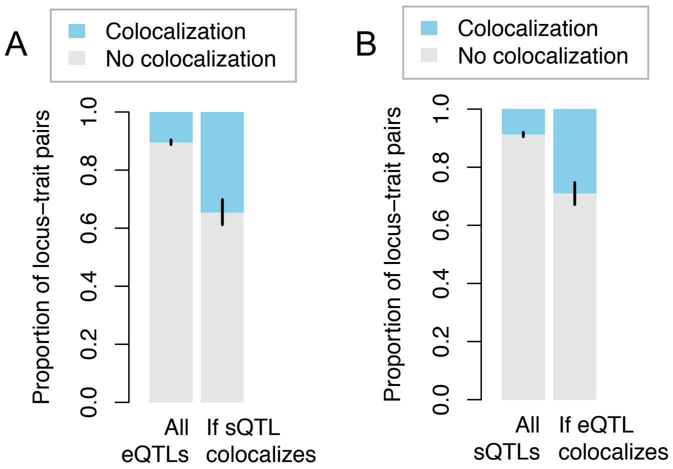

**Fig. S29. Concordant colocalization of *cis*-eQTLs and *cis*-sQTLs.** For GWAS loci that have an independent (LD-pruned) *cis*-eQTL and an *cis*-sQTL, A) shows the proportion of colocalising *cis*-eQTLs in all loci and in loci with a colocalizing *cis*-sQTL; and B) shows the proportion of colocalising *cis*-sQTL in all loci and in loci with a colocalizing *cis*-eQTL.

#### 12.7 Downstream phenotypic effect of regulatory pleiotropy and tissue sharing

We examined how variants affecting multiple genes in *cis* — or regulatory pleiotropy, sometimes called coregulation — in single tissues and across tissues contributes to their phenotypic consequences. To this end, we first compared how regulatory pleiotropy and cross tissue sharing associates with downstream GWAS association z-scores. We also compared regulatory pleiotropy with complex trait horizontal pleiotropy as defined in [53]. Briefly, for each *cis*-eVariant, we defined a score representing regulatory pleiotropy as the number of significantly associated genes. The tissue sharing status was defined as the tissue sharing pattern that most closely matched the *cis*-eVariant association vector across tissues. Next, we provide a detailed description of the procedure.

##### 12.7.1 Quantifying regulatory pleiotropy

First, we built a representative set of *cis*-eVariants filtered for potential confounding factors. To select likely causal variants, we used posterior inclusion probabilities calculated with the fine-mapping method DAPG [45] to select the most probable variant within each credible set of potentially causal *cis*-eQTL variants. The union of variants in all tissues were used. To remove any remaining correlation between selected variants, we performed LD pruning with $R^2 > 0.1$ cutoff inside a 100kb window (PLINK1.9 command `-indep-pairwise 100 10 0.1`). To reduce the bias introduced by sample size and also to increase the power for tissues with a low sample size, we applied MASH [54] on all possible variant/*cis*-gene pairs among the variants of interest across 49 tissues. Variants regulating no gene at MASH local false sign rate (LFSR) less than 0.05 were removed. For each eVariant in the set, we defined a score of regulatory pleiotropy, $P_n$, as the number of genes with LFSR < 0.05. We calculated $P_n$ in each tissue and also calculated an 'Aggregated' $P_n$, as the total number of genes the variant regulated across tissues. We found that regulatory pleiotropy was widespread, as shown in **fig. S30A**, which shows the fraction of variants regulating more than one gene within each tissue. To investigate how regulatory pleiotropy co-occurs across tissues, for each variant of interest, we counted the number of tissues in which the variant was regulating more than one gene. The distribution of this quantity, which measures the level of tissue-sharing for regulatory pleiotropy, is shown in **fig. S30B**. We observed that regulatory pleiotropy is extensively shared across tissues.

##### 12.7.2 Regulatory pleiotropy and GWAS associations

To examine the phenotypic consequences of regulatory pleiotropy, we regressed the significance of GWAS associations (z-score) on the regulatory pleiotropy score. We present the results using the dichotomized version of the regulatory pleiotropy score aggregated across tissues but verified that the substance of the results did not change when using the scores directly. The linear regression

$$z_{gwas}^2 \sim \mathbb{I}\{P_n > 1\} + \text{LD-score} + 1$$

was performed for each of the 87 traits. LD-score was calculated using genotypes of European individuals on GTEx variants at MAF $\geq$ 0.05 to account for high LD being a potential confounder of both regulatory pleiotropy and GWAS association. **Fig. S30C** shows the distribution of the coefficients of the regression. 66 out of 86 of the traits yielded a positive coefficient estimate, indicating a positive association between regulatory pleiotropy level and GWAS significance. Overall, we observed significantly positive correlation between regulatory pleiotropy level and GWAS significance at the genomic locus ($p = 6.1 \times 10^{-7}$, two-sided t-test with 87 traits) and the positive correlation remained after removing the set of blood cell count traits ($p = 2.9 \times 10^{-3}$, same test) and further removing one outlier ($p = 8.5 \times 10^{-5}$, same test). Thus, variants affecting multiple genes are more likely to contribute to downstream traits.

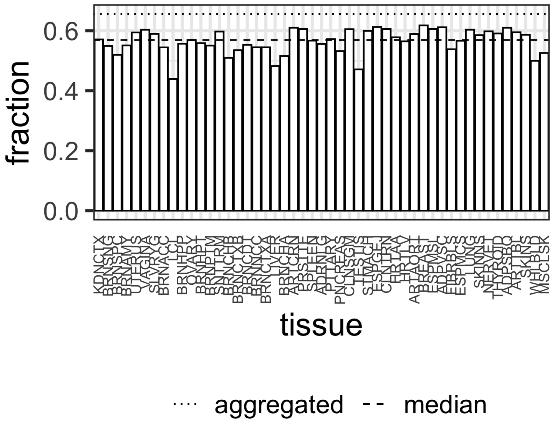

**(A)**

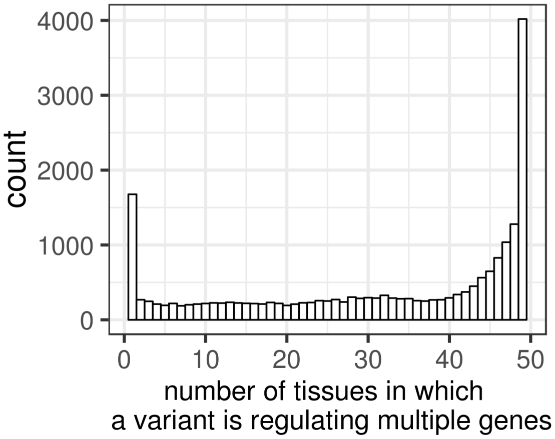

**(B)**

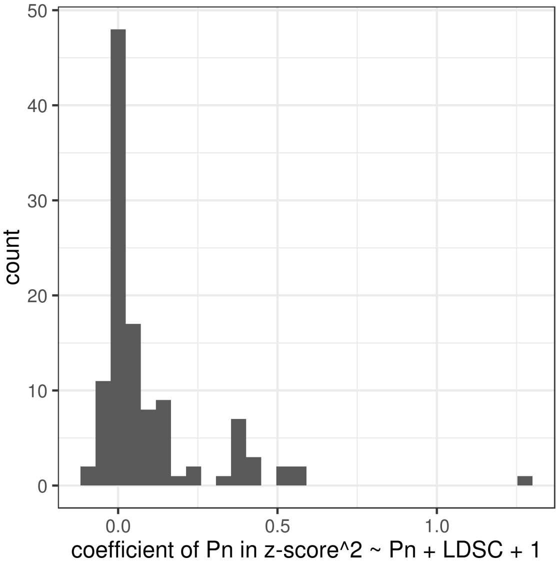

**(C)**

**Fig. S30. Regulatory pleiotropy of *cis*-eVariants.** A) Fraction of variants regulating multiple genes among a list of independent fine-mapped variants across 49 tissues (on x-axis ordered by sample size with abbreviations described in **table S8**). The dashed line indicates the median fraction among all tissues and the dotted line shows the fraction of variants regulating multiple genes in any tissue (aggregated across tissues). B) Number of tissues in which a variant is regulating multiple genes, which represents the level of tissue-sharing of regulatory pleiotropy. C) Distribution of regression coefficient for regulatory pleiotropy status when regressed against GWAS significance.

##### 12.7.3 Regulatory pleiotropy and trait pleiotropy

We hypothesized that when variants affect the expression of multiple genes, the downstream cellular effects of these genes may drive pleiotropic effects also at the level of complex traits. We compared regulatory pleiotropy to trait pleiotropy, which was measured by trait pleiotropy score for a set of 372 heritable traits from UK Biobank, $P_n$ as defined in [53]. The comparison was limited to the intersection between HapMap3 SNPs and the set of DAPG variants with MASH LFSR < 0.05. Among these variants, we contrasted the level of trait pleiotropy between those regulating only one gene and those regulating multiple genes (aggregated across all tissues) in **Fig. 5E**.

In order to analyze how regulatory pleiotropy across tissues contributes to trait pleiotropy, we used the FLASH approach [55] to classify *cis*-eQTLs into tissue-shared and tissue-specific sets (see details in [42]). Briefly, for every gene that was tested across all 49 tissues, the top associated variant was selected and used to find global patterns of tissue sharing using FLASH. One of the factors represented the broadly shared pattern (tissue-shared) and *cis*-eQTL's were considered to be tissue-shared if the projection of the vector of association effect sizes was large (proportion of variance explained > 20%).

To examine the relationship between tissue-sharing pattern and complex trait pleiotropy, we compared the distribution of trait pleiotropy (as stated above) between variants without any tissue-shared *cis*-eQTLs and variants with at least one tissue-shared *cis*-eQTL. The result is shown in **Fig. 5E** where the distribution for the latter set of variants shifts towards higher trait pleiotropy level. It suggests that variants with tissue-shared eQTLs tend to have higher GWAS pleiotropy.

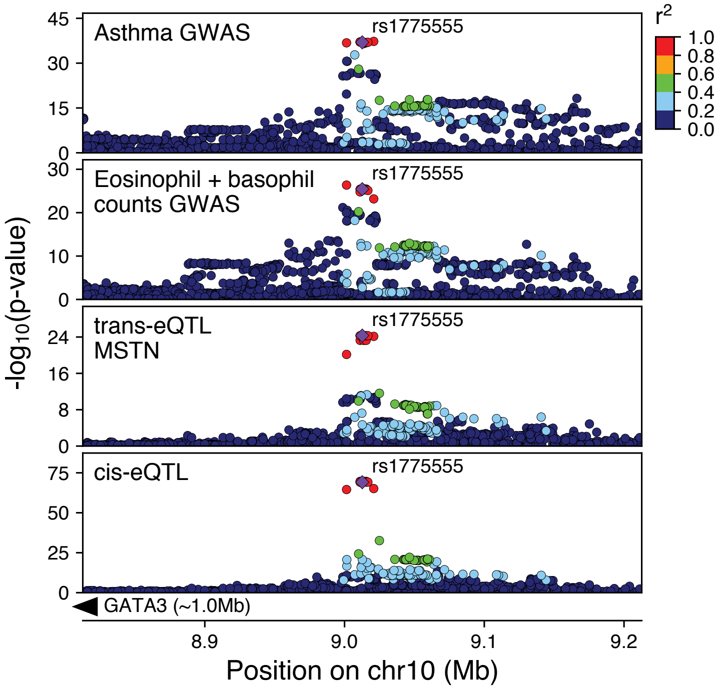

**Fig. S31. Associations at the *GATA3* locus.** Association landscapes for colocalizing associations for two GWAS traits, *trans*-eQTL for *MSTN*, and *cis*-eQTL for *GATA3* (gene location not shown due to it being nearly a megabase away).

### 13 Tissue sharing

#### 13.1 Estimating cross tissue activity of QTLs

In order to analyze tissue-specificity of QTLs, we used MashR [54] for every top *cis*-QTL per gene per tissue across all tissues where expression quantification was available. R v3.4.1 was used with mashr v0.2-6 and ashr v2.2-7. MashR was run using Z-scores as input, and 250,000 randomly selected SNP-gene pairs that were tested across all tissues were used to fit the mash model. Missing Z-score values (gene or splice quantification absent) were set to 0 and standard error to 1e6. Effect size estimates and local false sign rate (LFSR) outputted by MashR were used as metrics of QTL magnitude and activity respectively. A LFSR < 0.05 was used as a threshold for significant QTL activity unless otherwise noted.

We observed the previously shown [2, 15] U-shaped curve with *cis*-eQTLs and *cis*-sQTLs being typically highly shared or highly specific. In order to understand how these patterns are affected by LFSR threshold as well as differences in sensitivity of quantifying expression and splicing, we varied these thresholds (**fig. S32**). In particular, differences between expression and splice quantification sensitivity could result in power differences when calling *cis*-QTLs, and this might cause the observed results where *cis*-sQTLs have much higher cross-tissue activity than eQTLs when analyzing only those junctions/genes that were able to be tested in all tissues. Such a result could arise if there are substantial power differences caused by the quantification thresholds used for *cis*-sQTL vs *cis*-eQTL calling. While finding an equivalent threshold with respect to power between expression TPM and LeafCutter quantifications is challenging, an alternative option is to repeat the analysis using increasing levels of expression TPM thresholds for *cis*-eQTLs (**fig. S32**). While there is a clear pattern of increased tissue sharing at higher thresholds, even at the highest *cis*-eGene TPM threshold tested (TPM $>$ 100), which includes only 0.37% of *cis*-eQTLs, *cis*-sQTLs tested in all tissues (using the default thresholds) still showed much higher tissue sharing, indicating that the result is unlikely purely driven by a difference in detection power.

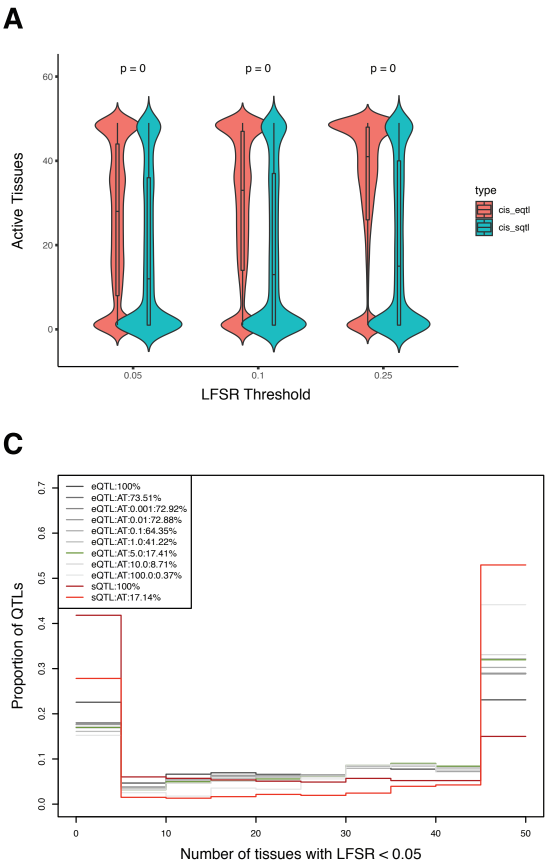

**Fig. S32. Thresholds in *cis*-QTL tissue sharing analysis.** A-B) Distributions of the number of active tissues per QTL for all *cis*-eQTLs and *cis*-eQTLs significant (FDR < 5%) in at least one tissue (A) and only those significant *cis*-QTLs that could be tested across all 49 tissues (B) at three different local false sign rate (LFSR, equivalent to FDR) thresholds. P-values are generated using a two-sided Wilcoxon signed rank test. Only *cis*-QTLs active in at least 1 tissue in the MashR meta-analysis were used. C) Tissue-specificity at different inclusion thresholds for gene expression and splicing. AT = tested in all tissues (using the standard thresholds from Sections 3.4.2 & 3.4.3), AT:0.001 = median TPM > 0.001 in all tissues, AT:0.01 = median TPM > 0.01 in all tissues, etc. The % of total QTLs that pass the threshold is listed in the legend.

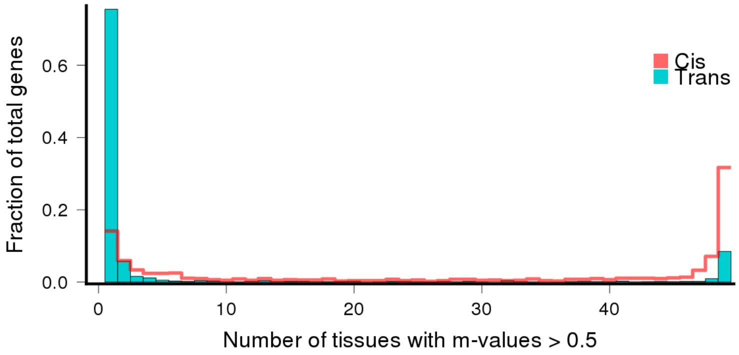

**Fig. S33. Tissue sharing of *cis*- and *trans*-eQTLs.** Distribution of the number of tissues having Meta-Tissue m>0.5 for the top variant for each *trans*-eGene at 50% FDR, and FDR-matched, randomly selected *cis*-eGenes (also 50% FDR). *cis*-eGenes were matched for discovery tissue to the *trans*-eGenes.

In order to estimate tissue-specificity of *trans*-eQTLs, and to ensure that the MashR results are robust *cis*-eQTL tissue-specificity estimates, we used the previously proposed linear mixed model approach that accounts for multiple tissues from the same individual and incomplete overlap of samples across tissues [56]. Briefly, for the number of tissues, T, we assume that there are $N$ incompletely overlapping individuals for each tissue. Then the following linear mixed model can be used to assess the statistical significance between gene expression $g$ and SNP $j$:

$$Y^g = 1\alpha + X_j\beta + u + e, \tag{12}$$

where $\beta$ is the effect coefficient of the SNP $j$ in $T$ tissues, and $u$ is the random effect of the mixed model accounting for the incompletely overlapping individuals across tissues. We estimated the two variance components using a linear mixed model approach as implemented in GEMMA [57].

Given the estimate $\hat{\beta}$, we combine the information from multiple tissues using meta-analytic approaches as implemented in metasoft [58], accounting for the covariance structure estimated as described above. We used the posterior probability that the effect exists in each study, m-value, as our estimate of tissue-specificity.

#### 13.2 Comparison of tissue clustering across data types

Estimates of pairwise tissue similarity were generated as similarly as possible across the 7 data types: (*cis*-eQTLs, *trans*-eQTLs, *cis*-sQTLs, ASE, splicing, expression, and cell type composition (see **fig. S34** for descriptions; *trans*-eQTLs were too few to be included). Only tissues that had pairwise similarity estimates across all data types were used, which excluded sex-specific organs since their similarity could not be estimated within individuals using ASE data. As shown in **fig. S34** and **Fig. 6a**, tissue clustering appears very similar across data types. In order to quantify this, we compared the clustering similarity. The fossil v0.3.7 package was used to perform k-means clustering using between 2 and 22 clusters with 50,000 iterations to ensure stability. At each cluster number a pairwise Rand index was calculated between each of the data types as a measure of clustering similarity. The median Rand index per pairwise data types across all cluster numbers was reported (**Fig 6B**).

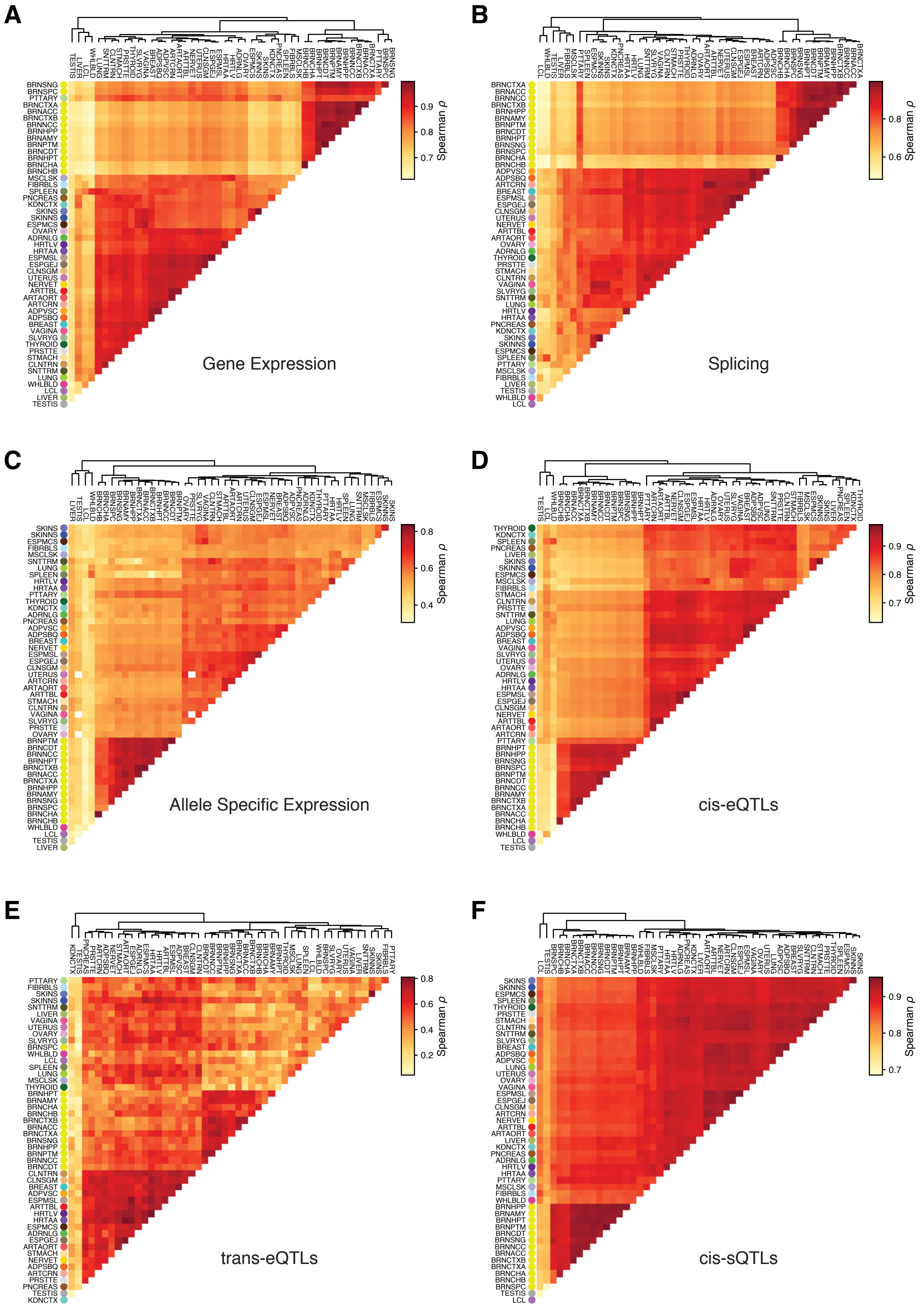

**Fig. S34. Pairwise tissue sharing.** Tissue clustering based on pairwise Spearman correlation for A) median tissue level gene expression (all genes with > 0 expression in at least one of the two pairwise tissues). B) median tissue level STAR splice junction counts (all junctions with > 0 counts in at least one of the two pairwise tissues). C) allelic expression (gene level AE for all genes with at least 8 reads in both of the pairwise tissues. In this analysis, the correlation is calculated intra-individual). D) *cis*-eQTL effect sizes (LD pruned set of all tissue level significant (FDR < 5%) top eQTLs per gene with MashR LFSR < 0.05 in at least one of the two pairwise tissues. E) *trans*-eQTL effect sizes (all tissue level significant (FDR < 5%) top *trans*-eQTLs per gene with Metasoft M-value > 0.50 in at least one of the two pairwise tissues. F) *cis*-sQTL effect sizes (LD pruned set of all tissue level significant (FDR < 5%) top eQTLs per gene with MashR LFSR < 0.05 in at least one of the two pairwise tissues.

#### 13.3 Allelic expression across tissues and tissue clustering

Allelic expression data from multiple tissues of the same individual can be used to evaluate the similarity of *cis*-regulatory genetic effects. We used a beta-binomial mixture to model allelic expression across tissues, with each component corresponding to one distinct mode of allelic ratio for a gene in an individual. Haplotypic counts were derived for each gene using phASER (see Section 5). Genes with at least 20 reads in at least six tissues were included in the analysis. In total, 6,614,043 gene-individual pairs were modeled using up to 10 modes of allelic ratio. Let us assume vector $\boldsymbol{x}$ to be the number of reads associated with the first haplotype and $\boldsymbol{y}$ be the total number of allelic reads in a given individual (haplotype labels are arbitrary). The data likelihood associated for a $K$-component mixture model is:

$$L(\boldsymbol{x}, \boldsymbol{y}|\lambda, \rho, \sigma) = \prod_i \sum_k \lambda_k BB(x_i, y_i, \rho_k, \sigma),$$

where $\lambda_k$ is the mixture weight for the $k^{\text{th}}$ component of the mixture ($\sum_k \lambda_k = 1$), and $x_i$, $y_i$ are the observed counts in the $i^{\text{th}}$ tissue. The Beta-binomial likelihood function, $BB(x_i, y_i, \rho_k, \sigma)$, was alternatively parameterized using a component-specific allelic ratio, $\rho_k$, and an over-dispersion parameter $\sigma$ shared by all $K$ mixture components (equivalent to standard parameterization using shape parameters $\alpha$ and $\beta$ for $\rho = \alpha/(\alpha + \beta)$, $\sigma = \alpha + \beta$).

A generic gradient-based optimization procedure (Matlab function `fmincon`) was used for maximizing the data likelihood with respect to $\lambda$, $\rho$, and $\sigma$. The optimization was done using four independent initial values for the parameters and the solution with the highest likelihood was selected. To select the number of components in the mixture model, $K$, we used a greedy approach: we started with a single mode mixture ($K = 1$), and increased $K$ by one at each step as long as the more complex model with $K + 1$ modes was significantly better ($p<0.01$) than the model with $K$ modes. Since the two models are nested and the overdispersion parameter is shared across components, we performed a likelihood ratio test using a chi-squared distribution with one degree of freedom to evaluate the goodness of fit at each step.

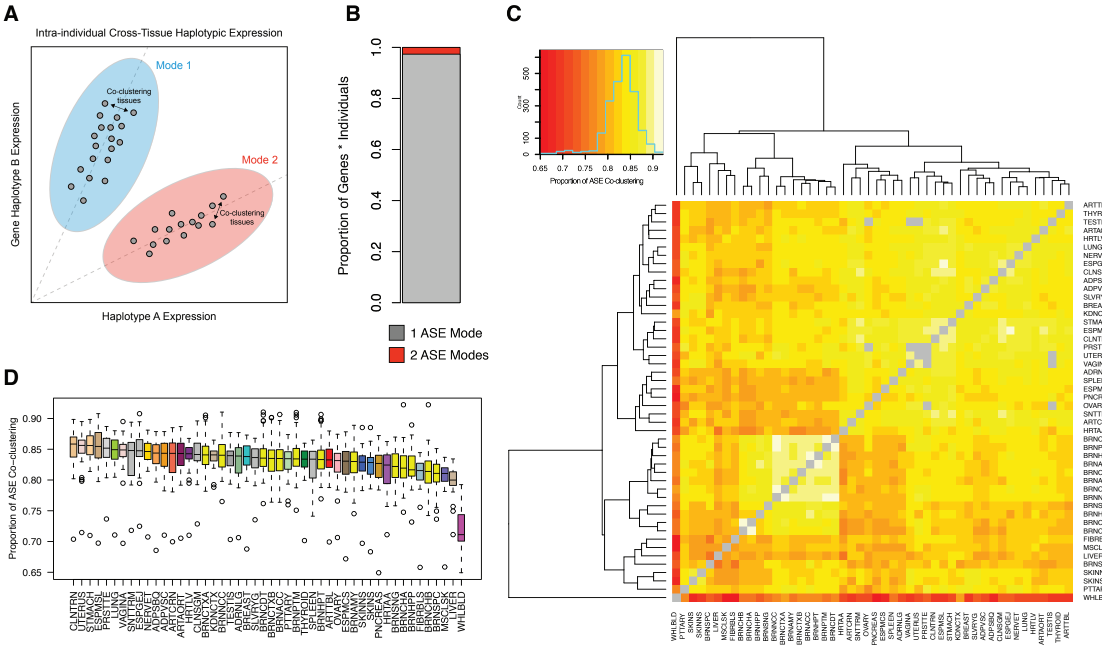

**Fig. S35. Tissue-specificity of allelic expression.** A) Within individuals, haplotypic expression across tissues was measured per gene and clustered to identify tissue specific patterns of *cis*-regulatory effects. Across individuals, tissues that often fall within the same cluster (co-cluster) share a higher degree of *cis*-regulatory architecture, as compared to those that do not. B) Proportion of gene x individual analyses with either one mode of allelic expression (grey, 97.34%), or two (red, 2.61%). Genes with three (0.04%) and four (6.35e-4%) modes are too few to be seen. C) Heatmap generated using the 2.61% of cases where two modes of allelic expression were observed and calculating what proportion of time each tissue appeared in the same cluster as other tissues. High co-clustering with other tissues indicates shared *cis*-regulatory architecture. D) Boxplot showing the distribution of co-clustering proportions per tissue. Each box represents a row (or column) from the heatmap.

#### 13.4 Correlation of *cis*-eQTL effect size and *cis*-eGene expression

Both tissue-specific and tissue-shared eQTLs may have different effect sizes across tissues. Thus, we examined the tissue variability of *cis*-eQTL effect sizes across tissues using the allelic fold change (aFC) statistic [25] (Section 6.1). We first explored the relationship of aFC with *cis*-eGene expression across all genes, and then in GWAS loci.

For each *cis*-eGene, we looked at all tissues with a significant *cis*-eQTL and selected the "top eVariant" with the largest effect size, and the tissue that it was found in was called the "discovery tissue." For each top eVariant, we calculated aFC across all tissues.

We then examine if there is a relationship between *cis*-eQTL effect size and *cis*-eGene expression level. For each top eVariant, we calculated the Spearman correlation of tissue aFC with *cis*-eGene median transcripts per million (TPM) for all tissues that had a median TPM greater than zero. We chose to focus on *cis*-eQTLs where the effect size does not flip from positive to negative between tissues, since those may be enriched for LD artefacts and the biological mechanism for opposite molecular effects is unclear. Thus, we filtered out *cis*-eQTL correlations that had "unclear" directions based on any of the following conditions: 1) over half of the tissue effect sizes were in the opposite direction of the discovery tissue effect; 2) any tissue effect size was in the opposite direction of the discovery tissue effect and had a magnitude of at least half the discovery tissue magnitude; or 3) the discovery effect size was 6.64 or higher, which is the maximum possible calculated aFC and often corresponds to *cis*-eQTLs with low allele frequency and unstable effect sizes (**fig. S36**). Next, in order to make the sign of the correlation coefficient interpretable for both positive and negative effect size *cis*-eQTLs, we flipped the sign of the effect sizes (multiplied by -1) if the discovery tissue *cis*-eQTL effect size was negative. This ensured that the discovery effect size was always labeled as positive and correlations could be interpreted the same way for both positive and negative effect size *cis*-eQTLs.

Of those *cis*-eQTLs where at least half of the GTEx tissues have a non-zero median *cis*-eGene expression, *cis*-eQTL effect size and *cis*-eGene expression level are significantly correlated across tissues for 2,637 top *cis*-eQTLs (5% Benjamini-Hochberg FDR; N=26,499; **fig. S36**). Of these, 666 are filtered out because of an unclear correlation direction. The remaining correlations are split nearly evenly to among positive/negative correlations with a positive/negative discovery tissue effect size (**fig. S36**). Positive correlations represent increasing effect size magnitude with increasing *cis*-eGene expression, and negative correlations represent decreasing effect size magnitude with increasing *cis*-eGene expression. The sign of the discovery tissue effect size represents whether the eVariant is associated with an increase (positive) or decrease (negative) in *cis*-eGene expression. *trans*-eGenes tended to have higher expression in their discovery tissue (**fig. S36**), but this may be due to better power in highly expressed tissues.

These results show that *cis*-eGene expression and *cis*-eQTL effects are not independent phenomena. Furthermore, it highlights the complicated nature of the classical question of in which tissues and cell types a given regulatory variant acts to cause downstream cellular, physiological, and disease phenotypes. Genes are generally believed to have the strongest functional role in tissues where they are highly expressed, but when this is combined with small molecular effect of the *cis*-eQTL (as in our negatively correlated genes), interpretation of the tissue(s) where that *cis*-eQTL might have a downstream effect becomes complicated.

#### 13.5 Cross-tissue *cis*-eQTL effect size and *cis*-eGene expression for GWAS genes

Previous analysis of GWAS loci has shown an enrichment of genes expressed in disease-relevant tissues [59], and the informativeness of *cis*-eQTLs in pinpointing causally relevant tissues has been debated [60, 61]. Thus, we sought to shed light on this question by analyzing colocalized and nearest genes in GWAS loci to examine whether aFC or *cis*-eGene expression are higher in potentially disease-relevant tissues.

##### 13.5.1 GWAS locus, tissue and gene selection

In order to study the cross-tissue *cis*-eQTL effect size and *cis*-eGene expression patterns in GWAS loci, we analyzed colocalized GWAS and *cis*-eQTL data from ENLOC as in Section 12.5.1. For a subset of GWAS traits we assigned a tissue group (blood, brain, immune, or metabolic) to each GWAS trait, and we then performed literature research to select hypothesized trait-relevant tissues for each trait (**table S9**). For each filtered, colocalized GWAS/*cis*-eQTL locus, we also built a set of the nearest protein-coding or lincRNA genes based on the absolute distance to the transcription start site (TSS). In order to remove redundancy in our dataset, we removed duplicated colocalized genes and nearest genes for each GWAS trait by first choosing one colocalized *cis*-eGene with the highest rcp per each nearest gene-GWAS trait pair, and then choosing one nearest gene with the closest TSS per each colocalized eGene-GWAS trait pair. This resulted in two gene sets, colocalized *cis*-eGenes and nearest genes, with each gene associated with one or more GWAS traits.

##### 13.5.2 Normalization across tissues

We next explored the tissue properties of colocalized and nearest GWAS genes, with the hypothesis that tissues with a potential causal role in disease should be enriched for high *cis*-eQTL effect size, high gene expression, or both. In this analysis, the colocalized *cis*-eGenes are compared to the "control" set of nearest genes. In order to achieve a fair comparison of tissues that differ in their overall expression profiles or regulatory effects (**fig. S37**), we used additional genome-wide background sets of tissue expression and *cis*-eQTL effects in all protein-coding and lincRNA genes. First, we determined the tissue with the highest significant *cis*-eQTL effect size (absolute aFC) and the tissue with the maximum median expression (TPM) for each gene in the

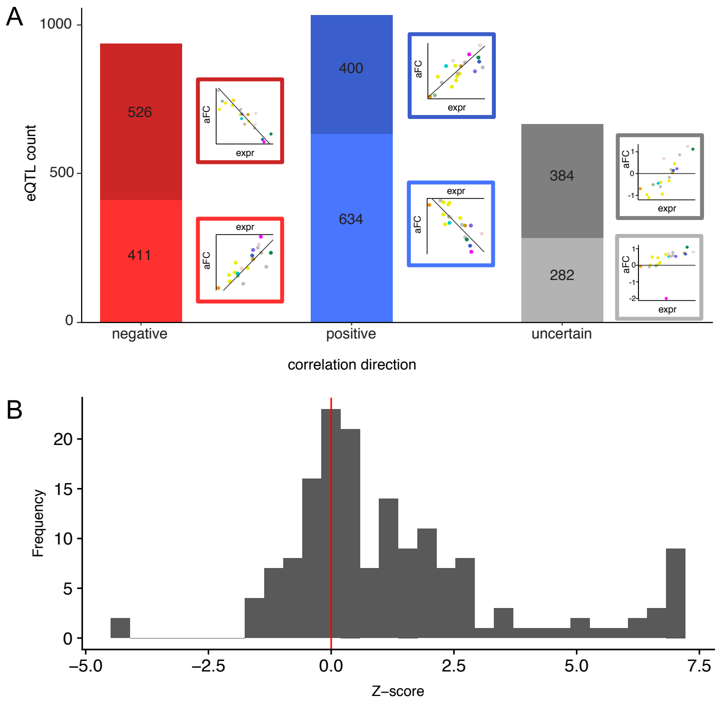

**Fig. S36. Correlation between *cis*-eQTL effect size and *cis*-eGene expression.** A) Top *cis*-eQTLs with a significant correlation between *cis*-eQTL effect size and median *cis*-eGene expression across tissues (5% Benjamini-Hochberg FDR; N=26,499). Cartoon examples of each type of correlation are shown, where each dot is a tissue and *cis*-eQTL effect size is plotted versus median *cis*-eGene expression. Note that for the uncertain direction correlations, there were additional patterns observed in the data that are not depicted here. B) Distribution of tissue-specific gene expression levels of *trans*-eGenes. For each *trans*-eGene, the Z-score is computed for its expression level in the corresponding tissue based on the empirical distribution of its expression levels in all other tissues, in each tissue assessing the mean across individuals. Mean gene expression in a tissue is measured by mean of log10(TPM + 1) across samples in this tissue. Among 162 *trans*-eGenes, There are 116 *trans*-eGenes with z-score > 0, and 46 *trans*-eGenes with z-score < 0, representing a strong excess of positive z-scores (p-value 3.7e-8, two-tailed binomial test).

GWAS gene sets and the background gene set. Next, we analyzed properties of GWAS genes in GWAS-trait-relevant tissues. We calculated tissue aFC (a) and expression (e) ranks for tissue t in gene i using rank statistics:

$$\textit{aFC rank statistic} = \mathrm{rank}(|a_i t| \text{ in } |A_i|)/N_i$$
$$\textit{Expression rank statistic} = \mathrm{rank}(e_i t \text{ in } E_i)/N_i$$

$A_i$ and $E_i$ are vectors of gene aFC and expression, respectively, in all tissues, and the ranks were normalized by N, the number of tissues that were not NA for the given measurement. Tissues that had a median TPM of 0 were assigned an expression rank statistic of 1/N.

We examined the distributions of aFC and expression rank statistics for trait-relevant tissues for colocalized *cis*-eGenes and nearest genes. To account for the tissue-specific aFC and gene expression patterns (e.g. blood has low expression levels for most genes), we performed paired Wilcoxon signed-rank tests between trait-relevant tissue rank statistics for colocalized and nearest genes and tissue-specific null rank statistics (the median rank statistic of a given tissue in our background gene set).

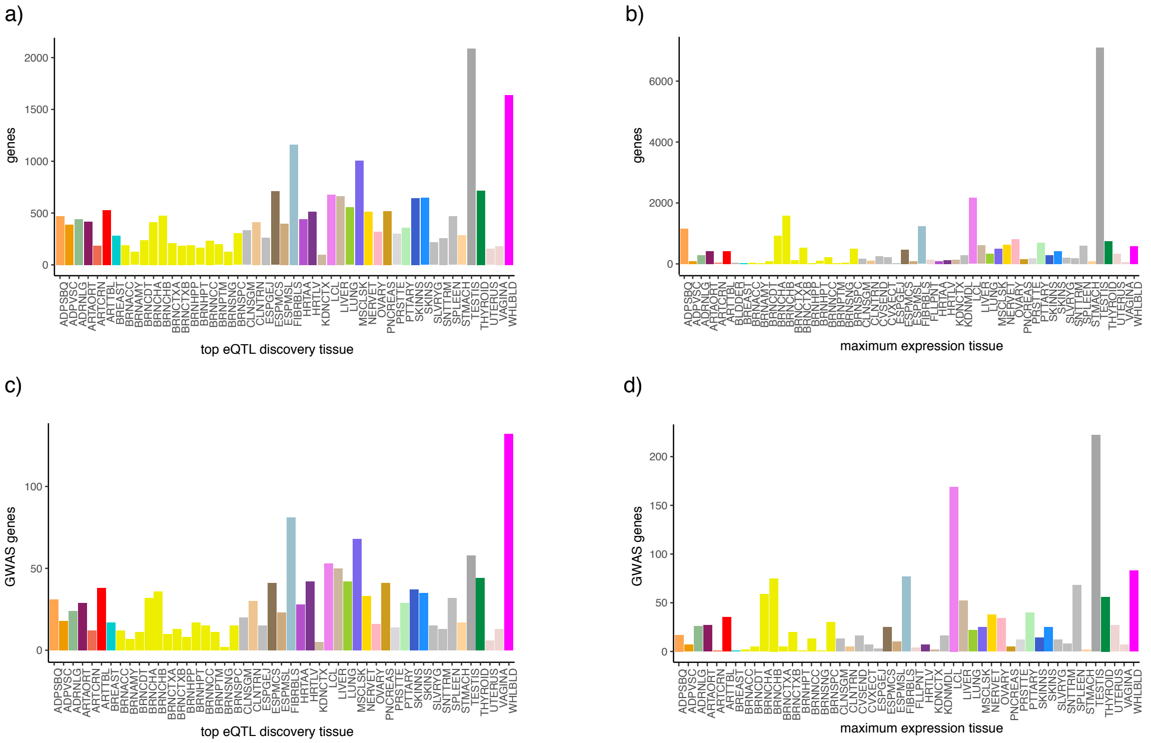

**Fig. S37. Tissue statistics for effect size and expression.** A) Tissue with the highest significant effect size for all protein-coding and lincRNA genes with a significant *cis*-eQTL (N=22,298). B) Tissue with the highest median expression for all protein-coding and lincRNA genes (N=26,724). C) Tissue with the highest significant effect size for each GWAS gene with a significant *cis*-eQTL (N=1,391). D) Tissue with the highest median expression for each GWAS gene (N=1,431). Note that the GWAS gene tissue patterns do not represent a generalizable pattern but rather reflect the traits that were available and selected for the analysis.

##### 13.5.3 Patterns of GWAS genes across tissues

We identified 2,157 filtered GWAS-eQTL ENLOC loci, corresponding to 1,110 colocalized and 1,096 nearest genes and 42 GWAS traits (**table S12**). 697 of these loci have different colocalized and nearest genes, and the union of all colocalized and nearest genes is referred to as "GWAS genes." The tissues with the highest significant *cis*-eQTL effect size and the tissues with the highest median TPM for GWAS genes and for genes in our background gene set (all protein-coding and lincRNA genes) are depicted in **fig. S37**. The distribution of tissues differs between the *cis*-eQTL effect size and the gene expression metrics.

Next, we analyzed how tissues potentially relevant for the GWAS traits ranked. For colocalized and nearest genes, the distributions of aFC and expression rank statistics for potentially trait-relevant tissues are shown in **fig. S38**. A rank statistic of one means the gene's highest effect size or expression level is in the potentially trait-relevant tissue, while 1/49 means the gene's lowest is in the tissue. When examining the rank statistic distributions, it is important to keep in mind the overall rank statistic distribution for each tissue, as observed in our background gene set (**fig. S38**). We found that GWAS genes have significantly higher effect sizes and expressions levels in the trait-relevant tissues than expected based on tissue median ranks in the background gene set (Paired Wilcoxon sign test, $p<1e-4$; **table S9**). This is observed both for colocalized and nearest genes.

These results suggest that both *cis*-eQTL effect size and expression level carry relevant information about the tissue that mediates downstream GWAS phenotype effects of genetic variant. Thus, examining both may be the best approach for understanding tissues that are causally relevant for human GWAS loci, even in the absence of gold-standards of causal tissues (or cell types) for a given trait, or specific loci.

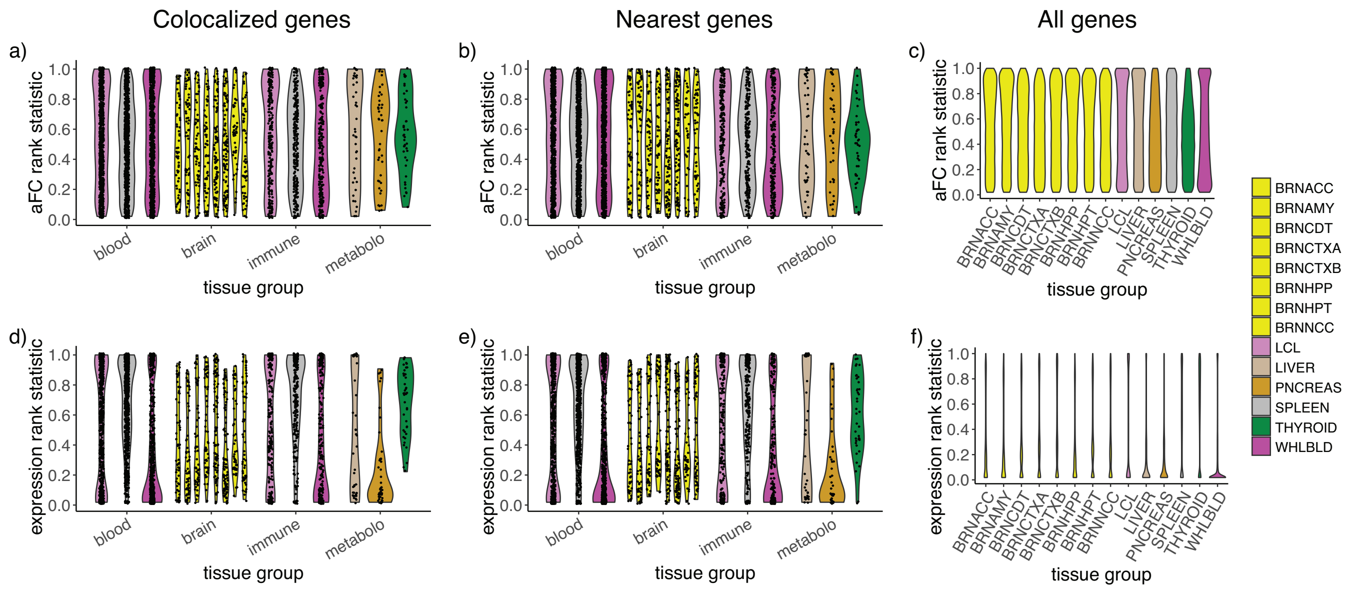

**Fig. S38. Tissue enrichment of effect size and expression for GWAS genes.** Tissue rank statistics in GWAS genes and all genes. In GWAS gene plots (a,b,d,e), each dot represents the rank statistic of a tissue in a GWAS gene, and violin plots are included to summarize the data. GWAS genes are split along the x-axis by the tissue group of the GWAS trait they are related to (blood, brain, immune, and metabolic). Rank statistics for each tissue in that group are then plotted separately for all genes that colocalize with (a,d) or are nearest to (b,e) the GWAS trait locus. Panels (a) and (b) display aFC tissue rank statistics in GWAS genes, while panel (c) depicts the aFC rank statistic distribution for those tissues across all protein-coding and lincRNA genes. Figures d and e display expression tissue rank statistics in GWAS genes, while figure c depicts the expression rank statistic distribution for those tissues across all protein-coding and lincRNA genes. The rank statistic distributions for all genes are included for reference, as some tissues tend to always have high or low relative aFC or expression across tissues.

| Gene selection | Rank method | Tissue-gene pairs | Median rank GWAS | Median rank null | P-value |
|---|---|---|---|---|---|
| colocalization | aFC | 3492 | 0.531 | 0.510 | 1.52e-04 |
| nearest | aFC | 3341 | 0.542 | 0.510 | 3.58e-06 |
| colocalization | expr | 3503 | 0.519 | 0.222 | 2.62e-294 |
| nearest | expr | 3460 | 0.481 | 0.222 | 4.57e-255 |

**Table S9.** aFC and expression rank statistics of GWAS genes

#### 13.6 Modeling determinants of QTL Tissue Specificity

In order to understand the factors that contribute to tissue-shared *cis*-eQTLs and *cis*-sQTLs, a logistic regression model of *cis*-QTL tissue activity was built to predict whether a QTL identified in a given discovery tissue is active in a given replication tissue, given a set of predictors: genomic annotations, tissue specific gene expression, and chromatin states. *cis*-QTL activity was defined as MashR LFSR $< 0.05$ in a replication tissue. Basic QC on the *cis*-QTL data used to build the model was performed as follows: expression / splice quantification $> 0$ in both discovery and replication tissues, *cis*-QTL MAF $> 0.005$ in both discovery and replication tissues, difference in expression level or splice quantification $>$ quantile(0.005) and $<$ quantile (0.995) to exclude the most extreme cases of expression or splice difference. R v3.5.1 was used with speedglm v0.3-2. When plotting model predictor coefficients (**Fig. 6**), they were standardized using the standardize R package v0.2.1 so that they could be plotted on the same scale. When reporting AUCs for the model including different sets of features, it was trained on *cis*-QTLs spanning chromosomes 1-20 and tested on QTLS from chromosomes 21 and 22. Otherwise, tissue level AUCs were generated by holding out holding out individual tissues and predicting the activity of *cis*-QTLs found in the other 21 tissues in the held-out tissue.

In total, 22 tissues were used for the analyses, which were chosen based on having appropriately paired epigenomic state predictions from the ROADMAP Epigenomics Project (**table S3**). In cases where there were two extremely similar tissues (defined by pairwise tissue gene expression clustering), the tissue with the higher sample size was used. Chromatin state sharing was defined as either shared on not shared based on if the ROADMAP Chromatin state prediction was the same (shared) or different (not shared) between the pairwise tissues. In total the following predictors were used in the model: distance between variant and TSS, variant MAF in GTEx, effect size in discovery tissue (aFC), global gene expression correlation between discovery and replication tissue, variant effect prediction, linsight conservation score [62], variant is INDEL, Roadmap state and sharing between discovery and replication tissue, variant overlaps Ensembl Regulatory Build TF binding site (in any Ensembl tissue), variant overlaps Ensembl Regulatory Build CTCF binding site (in any Ensembl tissue), variant overlaps Ensembl Regulatory Build DHS site (in any Ensembl tissue), variant overlaps Ensembl Regulatory Build predicted motif site.

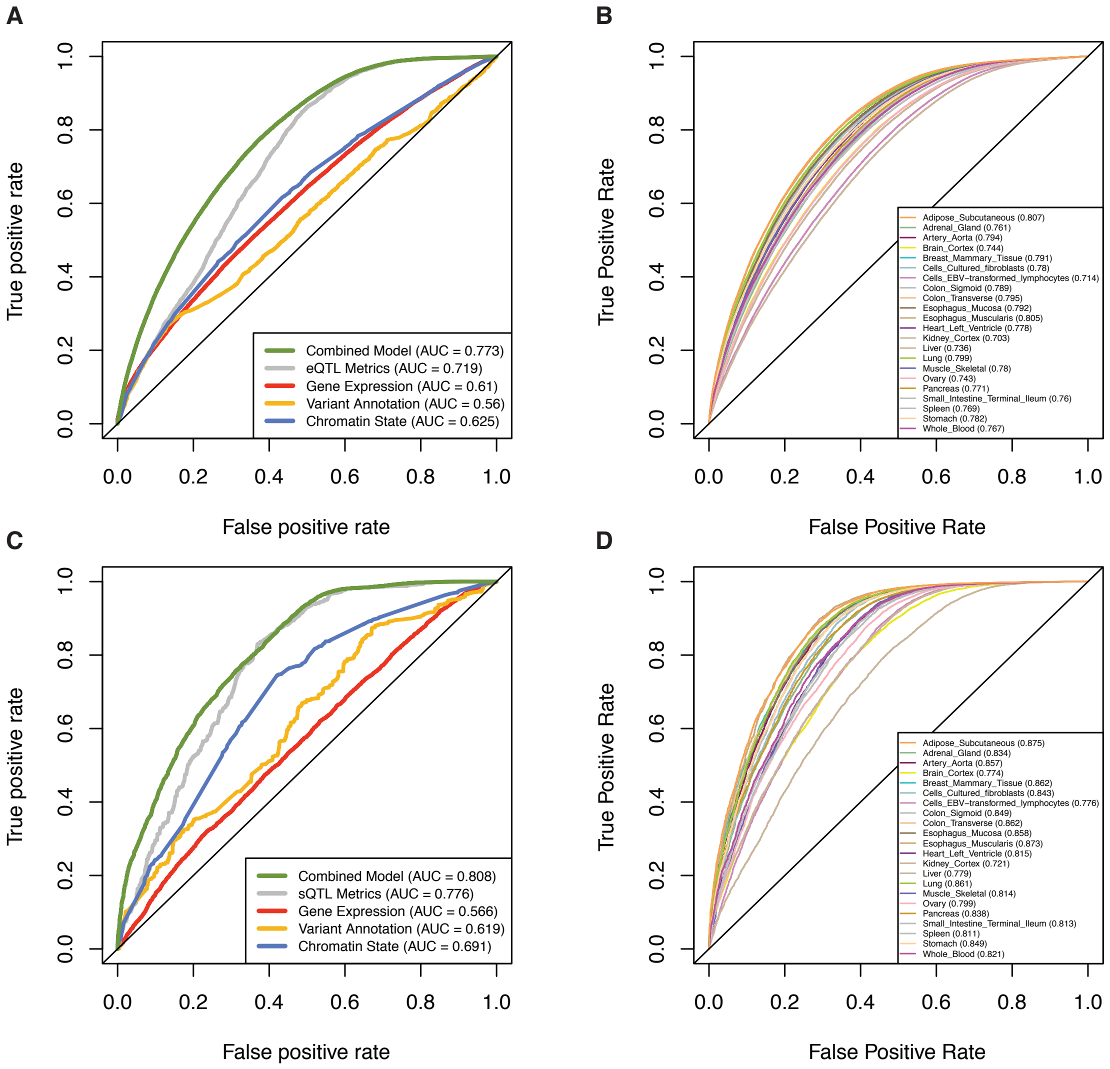

**Fig. S39. Predicting *cis*-eQTL and *cis*-sQTL activity in another tissue.** Receiver operating characteristic (ROC) curves for a logistic regression model of *cis*-eQTL and sQTL tissue sharing, defined as MashR LFSR $< 0.05$, using 22 GTEx tissues with paired Roadmap Epigenomics tissues and Ensembl Regulatory Build annotations. A) ROC curves for model trained on eQTLs from chromosomes 1-20 and tested on eQTLs from chromosomes 21-22. The performance of the model including different groups of predictors is shown: eQTL metrics (GTEx MAF + eQTL aFC + |tss_distance|), Gene Expression (|$\rho$ expr $\sim$ aFC| + $\rho$ Disc expr $\sim$ Rep expr + |$\Delta$ expr|), Variant Annotation (VEP effect + Variant INDEL + LINSIGHT Conservation), and Chromatin State (ROADMAP predicted chromatin state in discovery and replication tissues, and Ensembl Regulatory Build annotations). The performance of the full combined model is also indicated. B) ROC curves for model trained holding out individual tissues and then tested by predicting the activity of eQTLs found in the other 21 tissues in the held-out tissue. AUC is indicated in parenthesis beside each held out tissue name in the legend. C) ROC curves for model trained on *cis*-sQTLs from chromosomes 1-20 and tested on eQTLs from chromosomes 21-22. The performance of the model including only different groups of predictors is shown: sQTL metrics (GTEx MAF + sQTL beta + |tss_distance|), Gene Expression and Splicing ($\rho$ Disc splice junctions Rep splice junctions + |$\Delta$ expr|), Variant Annotation (VEP effect + Variant INDEL + LINSIGHT Conservation), and Chromatin State (ROADMAP predicted chromatin state in discovery and replication tissues, and Ensembl Regulatory Build annotations). The performance of the full combined model is also indicated. D) ROC curves for model trained holding out individual tissues and then tested by predicting the activity of *cis*-sQTLs found in the other 21 tissues in the held-out tissue. AUC is indicated in parenthesis beside each held out tissue name in the legend.

### 14 Cell type composition

#### 14.1 Estimation of cell type enrichment with xCell

Cell type enrichment scores were computed from the gene expression TPM matrix of the 17,382 samples in the analysis freeze with the `xCell` R package [63] using the `xCellAnalysis` function. The full matrix of expression data was used to maximize tissue heterogeneity. For QTL mapping, the enrichment scores corresponding to the subset of samples with available genotypes were inverse normal transformed.

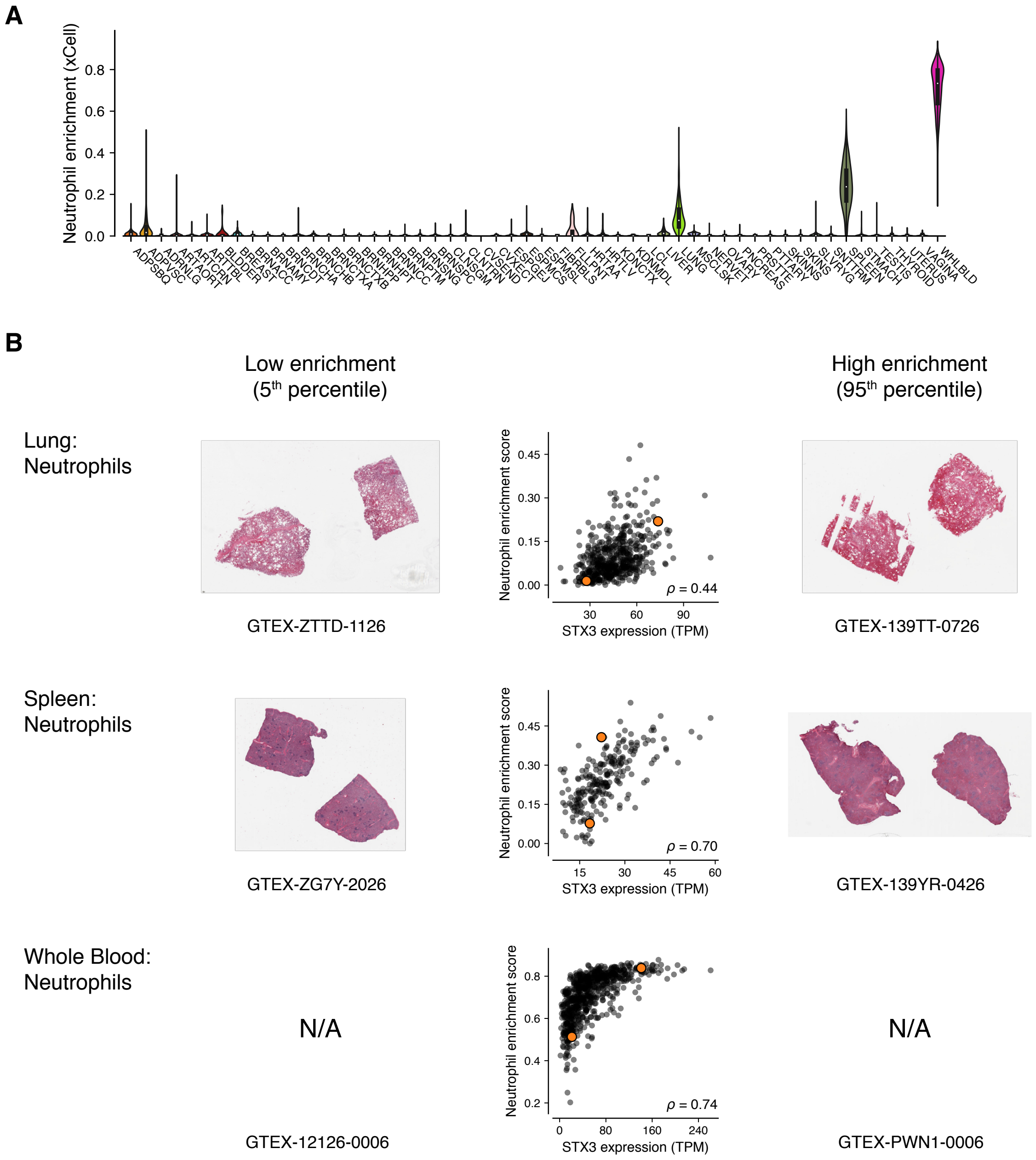

**Fig. S40. Neutrophil enrichment across GTEx tissues**. A) Distribution of xCell enrichment score for neutrophils across all samples of each tissue. B) Correlation between xCell neutrophil enrichment score and expression of *STX3*, a marker gene for neutrophils [64]. Orange dots indicate the samples closest to the 5th and 95th percentile of the neutrophil enrichment score, respectively, and the corresponding histology images are shown.

**Fig. S40** shows the distribution of neutrophil enrichment in GTEx samples, with the highest scores in whole blood, spleen and lung, which make physiological sense (lung tissue contains substantial amounts of blood cell types). Furthermore, there is substantial inter-individual variation in neutrophil enrichment within these tissues. The enrichment scores were highly correlated with expression of *STX3*, a marker gene for neutrophils [64], and histology images provided additional support that the enrichment scores represent true inter-individual variation in cell type composition. The median cell type enrichment per tissue was highly correlated between tissues, following general patterns of tissue relatedness (fig. **S41**).

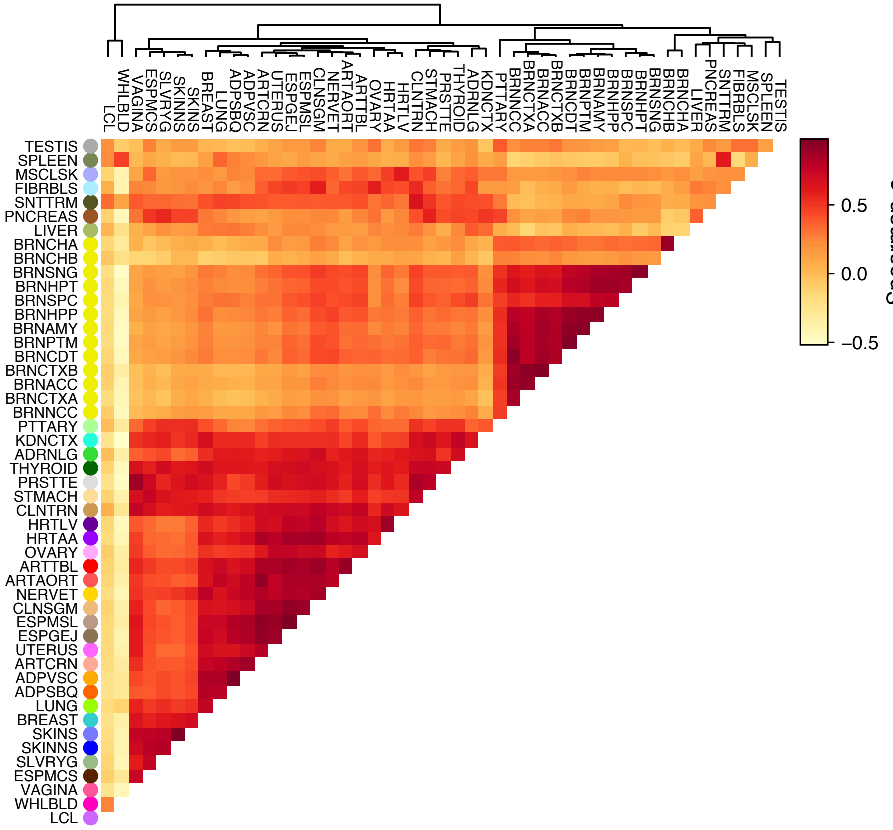

**Fig. S41. CT_TISSUE_HEATMAP Pairwise tissue sharing of cell type composition.** Tissue clustering generated using pairwise Spearman correlation on median xCell enrichment estimates across 64 cell types using cell types with $>0$ enrichment in at least one of the two pairwise tissues.

#### 14.2 Interaction QTL mapping

Cell type interaction eQTLs and sQTLs (ieQTLs and isQTLs, respectively) were mapped using a linear regression model with an interaction term accounting for interactions between genotype and cell type enrichment:

$$\boldsymbol{p} \sim \boldsymbol{g} + \boldsymbol{i} + \boldsymbol{g} \circ \boldsymbol{i} + \mathbf{C} \tag{13}$$

where $\boldsymbol{p}$ is the phenotype vector (e.g., gene expression or intron excision ratio), $\boldsymbol{g}$ is the genotype vector, $\boldsymbol{i}$ is the inverse normal transformed xCell enrichment score, and the interaction term $\boldsymbol{g} \circ \boldsymbol{i}$ corresponds to point-wise multiplication of genotypes and cell type enrichment scores. The same covariates, denoted by $\mathbf{C}$, were used as in regular QTL mapping (see Section 4.1). Interaction QTLs were mapped by testing for the significance of the interaction term, using tensorQTL [20]. TensorQTL computes regression coefficients and p-values for all terms in the model, enabling comparisons of interaction and main effects. Variants within $\pm$1Mb of the TSS of each gene were tested, as for regular QTL mapping. To avoid potential regression outlier effects, we restricted ieQTL mapping to variants with MAF $\geq$ 0.05 in the samples belonging to each of the top and bottom halves of the enrichment score distribution, for each tissue-cell type combination (using the `--maf_threshold_interaction 0.05` option in tensorQTL). For isQTL mapping, this threshold was set to MAF $\geq$ 0.1. The same filtered and normalized gene expression and splicing phenotype matrices used for regular QTL mapping were used for interaction QTL mapping. To identify genes with at least one significant ieQTL or isQTL (ieGenes or isGenes, respectively), the top nominal p-values for each gene or phenotype was corrected for multiple testing at the gene level using eigenMT [65]. Significance across genes was computed by adjusting the eigenMT-corrected p-values using Benjamini-Hochberg, and applying a 0.05 FDR threshold. For isQTLs, the p-value corresponding to the top splicing phenotype was selected for each gene-variant pair, and corrected by the number of phenotypes tested ($\tilde{p} = \min(n * p, 1)$, where $n$ is the number of splicing phenotypes for the gene) prior to running eigenMT. QTL mapping and FDR correction were performed using expression and splicing phenotypes for all biotypes in the GENCODE v26 annotation, but downstream analyses are based on protein coding and lincRNA genes only, as for regular QTLs. Due to high correlation of cell types in a given tissue (shown in **fig. S43** for blood), cell type interaction QTLs should not be interpreted as cell-type specific QTLs since they can represent regulatory effects in an (anti)-correlated cell type as well. Enrichment of ieQTLs and isQTLs in functional elements of the genome was calculated with Torus [33] (Section 11.1; **fig. S44**).

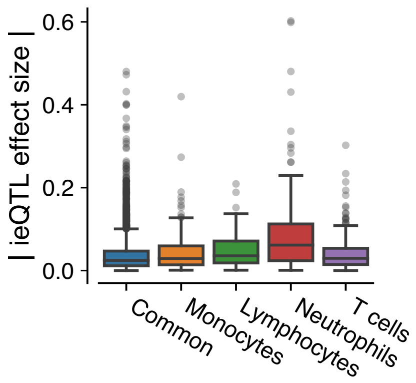

**Fig. S42. Replication of neutrophil ieQTLs in purified blood cell types.** eQTLs from purified neutrophils have higher median neutrophil ieQTL effect sizes than eQTLs from other cell types or eQTLs shared across cell types ('common'). eQTLs from purified blood cell types were obtained from [66].

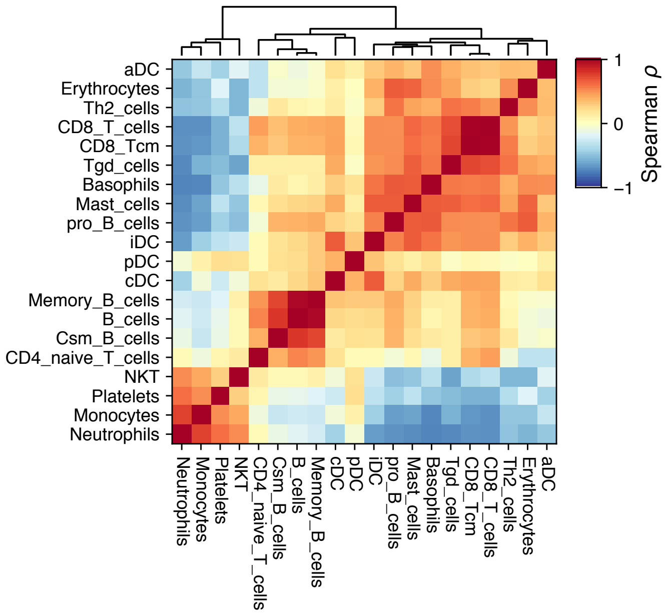

**Fig. S43. Correlation of blood cell types.** Spearman correlation between xCell enrichment scores for blood cell types in whole blood.

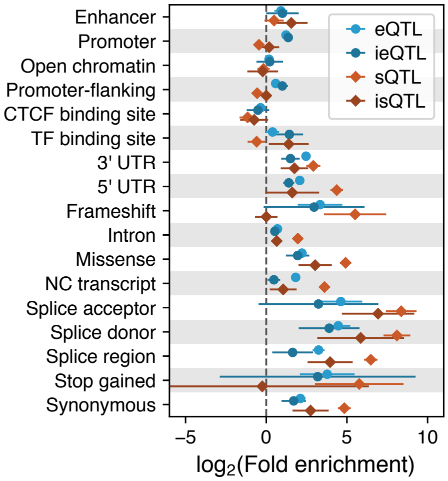

**Fig. S44. Functional enrichment of ieQTLs and isQTLs.** Enrichment in functional annotations of the genome for epithelial cell ieQTLs and isQTLs compared to cis-eQTLs and cis-sQTLs from transverse colon.

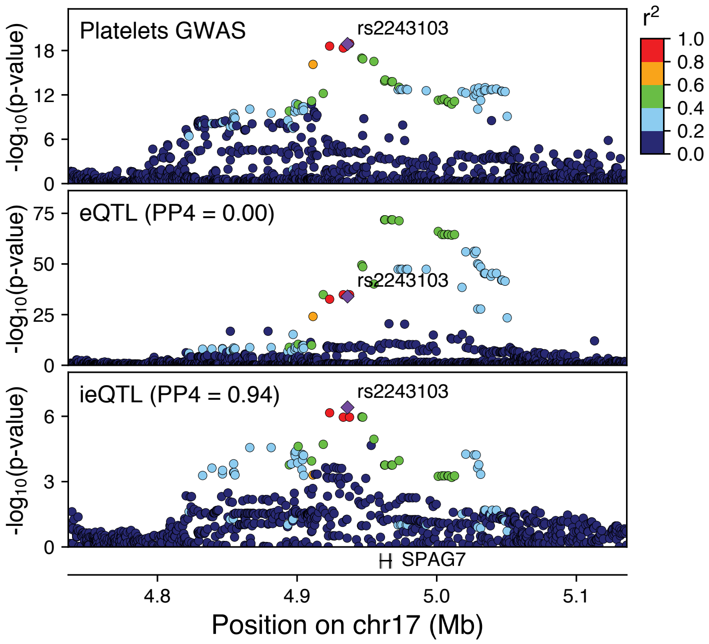

**Fig. S45. *SPAG7* ieQTL GWAS colocalization** P-value landscape in the SPAG7 locus for a platelet count GWAS [67], for bulk Whole Blood cis-eQTL associations (PP4 = 0.00), and neutrophil ieQTL associations (PP4 = 0.94). The top ieQTL variant (rs2243103) is highlighted.

#### 14.3 Colocalization of neutrophil ieQTLs and GWAS traits

Colocalization analysis was conducted using the `coloc` R package [21]. ENLOC, which was used in Section 12.5.1, could not be applied to cell type ieQTLs as DAP-G/ENLOC does not provide the option to include an interaction term when modelling the association (inputs are individual-level genotype and expression data sets). However, coloc priors can be computed from ENLOC estimates [18], and we therefore used coloc with model-based ENLOC priors (see below). Coloc uses summary statistics from QTL and GWAS studies in a Bayesian framework to identify GWAS signals that colocalize with QTLs. We ran coloc for all 1,120 neutrophil ieGenes at FDR $\leq$ 0.05 and 87 GWAS traits. All variants of the cis-QTL region ($\pm$1 Mb of the TSS of an ieGene) that were available for both the QTL and the GWAS trait were used in the function `coloc.abf()` with either cis-ieQTL or corresponding cis-eQTL p-values and GWAS effect size estimates and their variances. Given the high sensitivity of colocalization results to the choice of priors, we use model-based priors computed with *ENLOC* [18] (See Section 12.5.1). The same model-based priors used for cis-eQTLs were used for cis-ieQTLs assuming that regular cis-eQTLs reflect the average signal of all ieQTLs for a particular gene. The corresponding prior can thus be interpreted as the average prior of all ieQTLs for that gene and can be used as an approximated prior for each individual cell type ieQTL. We defined an ieGene or eGene as having evidence of colocalization when the posterior probability of colocalization (PP4) was higher than 0.5.

### 15 Supplementary Table legends

**Table S10. Population-biased eQTLs**. Statistics about high confident pb-eQTLs, including summary statistics for $\text{eVariant}_{\text{EA}}$ and $\text{eVariant}_{\text{AA}}$ (lead eVariant in an eQTL mapping analysis separately for individuals of European or African ancestry), and validation using allele-specific expression data.

**Table S11. GWAS Metadata** with relevant information about each GWAS study used. Columns are: **Tag**: Internal name to identify the study, **PUBMED_Paper_Link**: PUBMED entry, **Pheno_File**: name of downloaded file, **Source_File**: actual name of GWAS summary statistics (i.e. downloaded files might contain several traits), **Portal**: URL to GWAS study portal, **Consortium**: Name of consortium if any, **Link**: download link for the file, **Notes**: any special comment on the GWAS trait, **Header**: GWAS summary statistics header in case the file is malformed, **EFO**: Experimental Factor Ontology [68] entry if applicable, **HPO**: Human Phenotype Ontology [69] entry if applicable, **Description**: optional description of the study, **Phenotype**: phenotype name, **Sample_Size**: number of individuals included in the study, **Population**: types of populations present (EUR for European, AFR for African, EAS for East Asian, etc), **Date**: Date the file was downloaded, **Declared_Effect_Allele**: column specifying effect allele, **Genome_Reference**: Human Genome release used as reference (i.e. hg19, hg38), **Binary**: whether the trait is dichotomous, **Cases**: number of cases if binary trait, **abbreviation**: short string for figure and table display, **new_abbreviation**: additional abbreviation, **new_Phenotype**: additional phenotype name, **Category**: type of trait.

**Table S12. Pairing of GWAS traits and putatively relevant tissues**. GWAS traits were selected if they could be reasonably assigned to an affected tissue group: blood, brain, immune, or metabolic. The assigned tissue group and the literature-search-based trait-relevant tissues are displayed, as well as the final number of filtered loci included for each trait.

**Table S13. Per-tissue trans-eQTLs**. Trans-eQTLs across 49 tissues at gene-level FDR $< 0.05$ within each tissue. For each eGene in a tissue, the variant with the most extreme P value is selected as the lead variant.

**Table S14. Per-tissue trans-sQTLs**. Trans-sQTLs across 49 tissues at gene-level FDR $< 0.05$ within each tissue. For each eGene in a tissue, the variant with the most extreme P value is selected as the lead variant.

**Table S15. Trans-eQTL colocalization analysis**. Colocalization analysis of trans-eQTLs that overlap GWAS traits. Of the 25 trans-eQTLs that overlapped with a GWAS trait ($P<5e\text{-}8$), 10 colocalized with at least one trait at $PP4>0.9$.

### 16 Author Contributions

- François Aguet : Contributed to study design; Co-led the cell type composition analysis team with S.K-H.; Generated and QC'd the expression and splicing data; Contributed to genotyping pipeline development and QC; Developed the cis-QTL mapping pipelines and generated the cis-QTL data; Developed and ran the allelic expression pipelines with S.E.C; Developed the trans-sQTL mapping approach and performed related analyses; Contributed to fine-mapping and tissue-specificity analyses; Performed functional enrichment analyses; Performed the colocalization analyses between cis- and trans-eQTLs; Developed the cell type interaction QTL mapping approach and performed related analyses with S.K.-H.; Contributed text and supplementary materials for RNA-seq data processing and normalization, cis-QTL and trans-sQTL mapping, enrichment analyses, and other sections; Made all figures in the main text; Contributed to overall editing of the manuscript and supplement in the manuscript writing group.

- Kristin G Ardlie : Contributed to study design; Led the GTEx LDACC; Oversaw study design for processing, batching, generation and QC of all raw data; Supervised trainees who developed and ran QTL and expression pipelines; Led the GTEx portal efforts; Contributed to drafting and editing of the manuscript in the manuscript writing group.

- Alvaro N Barbeira : Harmonized and imputed public GWAS summary traits to hg38; Participated in colocalization and fine-mapping analyses; Performed DAP-G and ENLOC GWAS-QTL analyses; Generated PrediXcan transcriptomic prediction models, S-PrediXcan and S-MultiXcan results; Contributed to text, figures, and supplementary materials for the GWAS section.

- Alexis Battle : Contributed to study design; Co-led the trans-eQTL analysis team with B.E.E.; Helped design and oversee the trans-eQTL analysis pipeline and QC; Helped oversee downstream analysis of trans-eQTL characterization and examples; Contributed to the writing of the trans-eQTL sections of the manuscript and supplement; Supervised the work of trainees in her lab on trans-QTL characterization; Participated in the manuscript writing group.

- Andrew Brown : Co-led the fine mapping analysis team; Performed the CaVEMaN fine-mapping analysis, Coordinated and collated results across the three fine mapping methods; Contributed text, figures, and supplementary materials for the fine-mapping section.

- Christopher D Brown : Provided advice on GWAS analyses; Supervised the work of trainees in his lab on GWAS; Provided advice to the writing group.

- Rodrigo Bonazzola : Contributed to harmonization and imputation of GWAS summary statistics; Contributed to colocalization methods benchmarking; Performed the analysis of primary and secondary eQTL contribution to GWAS signals; Performed the analysis of mediating effects of eQTLs and sQTLs on complex traits; Contributed to text, figures, and supplementary materials for the GWAS section.

- Stephane E Castel : Led the tissue specificity analysis team; Developed the pipeline for generation and quality control of ASE data; Performed descriptive analysis of ASE data; Assisted with generation of ASE data for the purposes of validating sex, population, and cell type specific eQTLs; Calculated and analyzed cross tissue activity estimates for cis-e/s-QTLs; Analyzed tissue clustering across all core data types; Modeled determinants of cis e/s-QTL tissue specificity using genomic annotations; Contributed text, figures, and supplementary materials for the tissue specificity section.

- Nancy Cox : Contributed to study design; Supervised the work of trainees in her lab.

- Sayantan Das : Performed genotype imputation for dap-g analysis.

- Emmanouil T Dermitzakis : Contributed to study design; Supervised the work of trainees in his lab on fine-mapping and eQTL replication.

- Barbara E Engelhardt : Co-led the trans-eQTL analysis team with A.B.; Helped design and overseee the trans-eQTL analysis pipeline and QC. Supervised the work of trainees in her lab on trans-eQTLs.

- Elise Flynn : Performed analysis of the relationship between eQTL effect sizes and eGene expression across tissues; Analyzed tissue properties of GWAS-colocalized eQTLs; Mapped and characterized the CBX8 fine-mapping example. Contributed text, figures, and supplementary materials for the corresponding sections.

- Laure Fresard : Performed the annotation of non-eQTL genes; Contributed supplementary materials for non-eQTLs.

- Eric R Gamazon : Performed GWAS enrichment and heritability analyses; Contributed to interpretation of GWAS analyses.

- Diego Garrido-Martín : Contributed to benchmarking of cell type enrichment estimates.

- Nicole R Gay : Performed analyses based on population structure.
- Gad Getz : Contributed to study design; Oversaw computational and analysis pipeline development and optimization team in the LDACC.
- Roderic Guigó : Contributed to study design; Supervised the work of trainees in his lab on cell type composition and splicing.
- Andrew R Hamel : Developed QTLEnrich; Performed GWAS-QTL enrichment analyses with QTLenrich.
- Robert E Handsaker : Performed WGS analysesto identify aneuploidies, atypical sex chromosomes, large germline CNVs and mosaic CNVs.
- Yuan He : Performed enrichment analysis of cis-QTLs among trans-QTLs; Performed cis-trans mediation analysis; Contributed to analysis of sample size and cis-eQTL discovery underlying trans-eQTL discovery; Contributed supplementary materials for the corresponding sections.
- Paul J Hoffman : Contributed to effect size and interaction QTL pipeline development; Assisted with manuscript formatting.
- Farhad Hormozdiari : Led the functional annotation analysis team; Performed S-LDSC analysis for GWAS traits including cross trait genetic correlation; Performed fine-mapping by CAVIAR; Contributed to text, figures, and supplementary materials of the corresponding sections.
- Hae Kyung Im : Contributed to study design; Led the GWAS analysis team; Coordinated and led the design, analysis, resource sharing, and interpretation of the GWAS results; Supervised the work of trainees in her lab on diverse GWAS analyses; Contributed to the writing of the GWAS sections of the manuscript and supplement; Participated in the manuscript writing group.
- Brian Jo : Mapped the trans-eQTLs; Performed the cis-QTLs - trans-QTL mediation analysis.
- Silva Kasela : Mapped and analyzed the pb-eQTLs; Performed ASE replication of sex, population and cell type interaction eQTLs; Performed the analysis of concordance of GWAS effect for rare variants and eQTLs; Performed the analysis of concordance of GWAS effect for eQTL and sQTLs. Provided text, figures, and supplementary materials for these sections.
- Seva Kashin : Performed WGS analysesto identify aneuploidies, atypical sex chromosomes, large germline CNVs and mosaic CNVs.
- Sarah Kim-Hellmuth : Co-led the cell type composition analysis team with F.A.; Developed pipelines and performed cell type enrichment estimation and cell type ieQTL mapping; Supervised benchmarking of cell type deconvolution methods. Led the downstream analyses of ieQTL properties; Developed pipelines and assisted in multi-tissue cis-eQTL analysis and pairwise tissue-sharing analysis; Developed pipelines and assisted in torus enrichment analysis; Contributed to sex-biased eQTL analysis; Contributed to cis-/trans-eQTL interpretation; Contributed to colocalization methods testing; Contributed text, figures, and supplementary materials for the cell type section.
- Alan Kwong : Performed genotype imputation for dap-g analysis.
- Tuuli Lappalainen : Contributed to study design; Led and coordinated the work of the manuscript working group and the manuscript writing group; Performed the TAD enrichment analysis; Supervised the work of trainees in her lab on cell type composition, tissue-specificity, pb-eQTLs, shared rare variant and eQTL-sQTL contribution to GWAS signals, ASE, and effect size; Wrote the first draft of the manuscript; Contributed figures and text for the supplement; Led the writing and editing of the manuscript and supplement.
- Xiao Li : Developed and applied the sample and variant quality control pipeline to the WGS and WES data; Performed variant annotation of the WGS and WES data; Assisted in the long copy number variation analysis; Evaluated different phasing methods on the WGS genotype data.Contributed supplementary materials for these sections.
- Yanyu Liang : Performed the analysis on the enrichment of GWAS in e/sQTLs; Performed the analysis of regulatory pleiotropy and their downstream effect on complex phenotypes; Performed the analysis of GWAS effects of primary and secondary QTLs; Performed the analysis of mediating effects of eQTLs and sQTLs on complex traits; Developed mixed effects models for aggregation across phenotypes and tissues; Contributed to text, figures, and supplementary materials for the GWAS section.
- Daniel G MacArthur : Contributed to the generation and quality control of genotyping and DNA sequencing data.

- Pejman Mohammadi : Performed the analysis of ASE sharing across tissues; Contributed to ASE validation of sex-biased eQTLs; Contributed supplementary materials for ASE tissue-sharing.
- Stephen B Montgomery : Supervised the work of trainees in his lab on population structure and non-eQTL genes; Provided advice to the writing group.
- Manuel Muñoz-Aguirre : Contributed to benchmarking of cell type deconvolution methods. Contributed to pairwise tissue sharing based on expression and cell type composition.
- Meritxell Oliva : Co-led the sex analysis team with B.E.S.; Mapped and analyzed the sb-eQTLs; Optimized, benchmarked and replicated cell type enrichment and ieQTL analyses with F.A. and S.K.-H.; Contributed text, figures, and supplementary materials for sb-eQTLs.
- YoSon Park : Contributed to colocalization analyses and methods benchmerking; Performed DAP-G/ENLOC enrichment analyses; Performed cis-eQTL tissue-sharing analysis by Meta-Tissue; Contributed to GWAS data processing; Contributed supplementary materials for tissue-sharing and GWAS.
- Princy Parsana : Performed analysis of trans-eQTL - GWAS overlap; Performed analyses of potential trans-eQTL vignettes; Contributed to text and supplementary materials of the GATA3 locus.
- John M Rouhana : Programmed the new permutation module for QTLEnrich; Contributed to GWAS-QTL enrichment analysis.
- Ashis Saha : Participated in trans-eQTL pipeline development and QC.
- Ayellet V Segrè : Supervised the whole genome sequencing data processing and sample and variant QC and the development of the WGS QC pipeline; Contributed to optimization of the QTL pipeline; Supervised the work of trainees in her lab, including the development of QTLEnrich; Contributed text, figures, and supplementary materials for WGS and QTLEnrich sections.
- Matthew Stephens : Provide advice and software support for multi-tissue eQTL analyses by MashR; Supervised the work of trainees in his lab.
- Barbara E Stranger : Co-led the sex analysis team with M.O.; Supervised the work of trainees in her lab on cell type composition and sex-biased eQTLs.
- Benjamin J Strober : Developed and applied Meta-Tissue analysis pipelines for tissue-specificity and sharing of cis- and trans-eQTLs.
- Ellen Todres : Coordinated all sample processing and sequencing.
- Ana Viñuela : Contributed to replication of ieQTLs and trans-eQTLs.
- Gao Wang : Contributed to multivariate analysis for cis-eQTLs and interaction eQTLs; Contributed to eQTL fine-mapping.
- Xiaoquan Wen : Co-led the fine-mapping analysis team with A.B.; Contributed to colocalization and PrediXcan analyses; Provided advice on the GWAS analyses.
- Valentin Wucher : Contributed to benchmarking of cell type deconvolution methods; Contributed to pairwise tissue sharing based on expression and cell type composition.
- Yuxin Zou : Provided advice for MashR analysis.
